# Distributable, Metabolic PET Reporting of Tuberculosis

**DOI:** 10.1101/2023.04.03.535218

**Authors:** R.M. Naseer Khan, Yong-Mo Ahn, Gwendolyn A. Marriner, Laura E. Via, Francois D’Hooge, Seung Seo Lee, Nan Yang, Falguni Basuli, Alexander G. White, Jaime A. Tomko, L. James Frye, Charles A. Scanga, Danielle M. Weiner, Michelle L. Sutphen, Daniel M. Schimel, Emmanuel Dayao, Michaela K. Piazza, Felipe Gomez, William Dieckmann, Peter Herscovitch, N. Scott Mason, Rolf Swenson, Dale O. Kiesewetter, Keriann M. Backus, Yiqun Geng, Ritu Raj, Daniel C. Anthony, JoAnne L. Flynn, Clifton E. Barry, Benjamin G. Davis

## Abstract

Tuberculosis remains a large global disease burden for which treatment regimens are protracted and monitoring of disease activity difficult. Existing detection methods rely almost exclusively on bacterial culture from sputum which limits sampling to organisms on the pulmonary surface. Advances in monitoring tuberculous lesions have utilized the common glucoside [^18^F]FDG, yet lack specificity to the causative pathogen *Mycobacterium tuberculosis* (*Mtb*) and so do not directly correlate with pathogen viability. Here we show that a close mimic that is also positron-emitting of the non-mammalian *Mtb* disaccharide trehalose – 2-[^18^F]fluoro-2-deoxytrehalose ([^18^F]FDT) – can act as a mechanism-based enzyme reporter in vivo. Use of [^18^F]FDT in the imaging of *Mtb* in diverse models of disease, including non-human primates, successfully co-opts *Mtb*-specific processing of trehalose to allow the specific imaging of TB-associated lesions and to monitor the effects of treatment. A pyrogen-free, direct enzyme-catalyzed process for its radiochemical synthesis allows the ready production of [^18^F]FDT from the most globally-abundant organic ^18^F-containing molecule, [^18^F]FDG. The full, pre-clinical validation of both production method and [^18^F]FDT now creates a new, bacterium-specific, clinical diagnostic candidate. We anticipate that this distributable technology to generate clinical-grade [^18^F]FDT directly from the widely-available clinical reagent [^18^F]FDG, without need for either bespoke radioisotope generation or specialist chemical methods and/or facilities, could now usher in global, democratized access to a TB-specific PET tracer.

## Introduction

Tuberculosis (TB) caused by *Mycobacterium tuberculosis* (*Mtb*) still remains a serious global health challenge causing an estimated 1.5 million deaths worldwide in 2020 following the first year-on-year increase since 2005.^1^ Prompt, short-term diagnoses of TB are crucial for public health infection control measures, as well as for ensuring appropriate treatment for infected patients and controls.^2^ However, 2019-2020 saw a 1.3 million (18% of total) drop in diagnoses globally despite estimated increasing disease levels.^1^ Additionally, long-term accurate monitoring of chronic disease burden and the effectiveness of treatment is critically important in trials of new antitubercular agents and regimens. Sensitive and TB-specific reporters with the potential for ready democratization are therefore urgently required to address the development of new antituberculosis agents and regimens with the potential to shorten the duration of therapy.

Positron emission tomography (PET) integrated with computed tomography (CT) now routinely provides a prominent method in some (and increasing^3^) national healthcare systems for noninvasively imaging the whole body whilst diagnosing, staging and assessing response to therapy in diseases such as cancer and inflammation. The analysis^4^ and internationally-agreed, comprehensive monitoring of access to PET-CT (e.g. by the **I**AEA **M**edical im**AGI**ng and **N**uclear m**E**dicine global resources database, IMAGINE^5^) helps, in part, to drive global equity of access to such diagnostic imaging. However, this would be importantly aided by development of (i) novel reporters specific to other (e.g. communicable) diseases of similar, or even greater, relevance to the developing world, such as TB, and (ii) strategies and methods for their ready, distributed implementation.

Existing *metabolic* PET probes allow pharmacological, immunologic and microbiological aspects of TB lesions to be correlated with anatomic information derived from PET/CT.^6,7^ In particular, 2-[^18^F]fluoro-2-deoxy-D-glucose ([^18^F]FDG, often shortened to “FDG”, **Figure 1**), as the most widespread, organic ^18^F-containing probe, has allowed useful diagnosis and monitoring of the response to treatment of TB.^8,9^ Specifically, the high metabolic uptake of [^18^F]FDG into the host cells around pulmonary and extra-pulmonary lesions of active TB can allow PET/CT imaging of TB granulomas by inference and so attempts to indirectly assess disease extent and progression or resolution.^10^ [^18^F]FDG PET exploits enhanced glucose uptake into the anaerobic glycolytic pathway both in tumor cells and in immune cells – [^18^F]FDG PET in imaging of infections relies therefore typically on enhanced glucose uptake of inflammatory cells as a result of associated respiratory burst. However, by virtue of this physiological mechanism, [^18^F]FDG is also taken up and retained by *any* ‘metabolically active’ tissue and so as a generic ‘marker’ of more active metabolism has a limitation in its lack of specificity and inability to clearly distinguish granulomatous TB disease from other inflammatory conditions, including cancer. This ‘inferred imaging’ mode therefore frequently gives rise to false-positive diagnosis in patients evaluated for active TB.^11-13^ This diagnostic confusion when using [^18^F]FDG has been further exacerbated by the wide-spread emergence of lung pathology in COVID-19 patients.^14^

**Figure 1.**
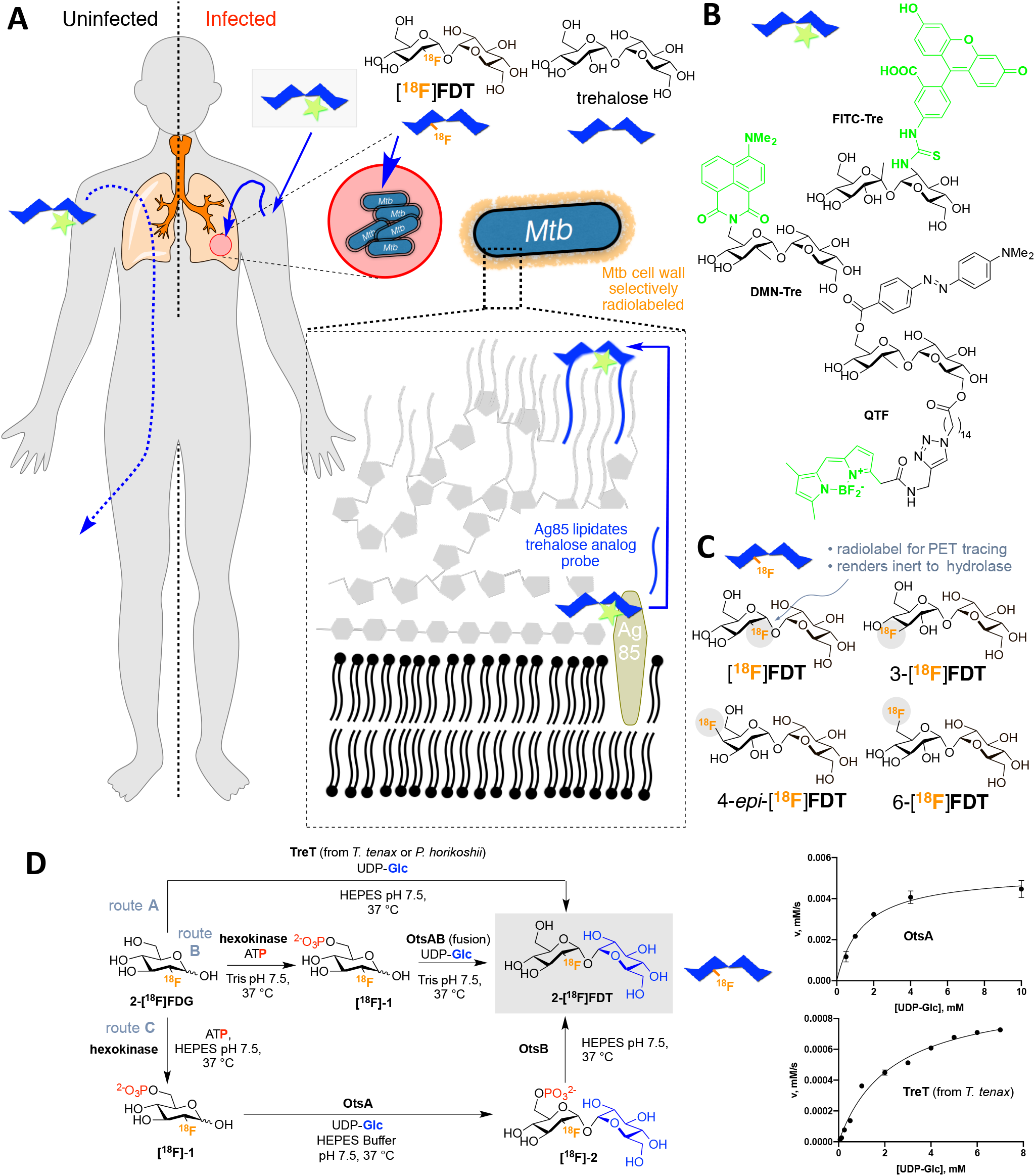
A Strategy for Non-Invasive Imaging Reporters using Trehalose-based, TB-specific Probes Derived Directly from [^18^F]FDG. **A)** Trehalose (blue) in *Mtb* is found in the outer portion of the mycobacterial cell envelope as its corresponding mycolate glycolipids. The biosynthesis of trehalose mycolates (lipidation) is catalyzed specifically in *Mtb* by abundant membrane-associated Antigen 85 (Ag85a, Ag85b and Ag85c) enzymes. The lack of naturally occurring trehalose in mammalian hosts as well as the uptake of exogenous trehalose by *Mtb* suggests that it could function as both a highly specific and sensitive probe, allowing here the development of an *in vivo* TB-specific, PET-radiotracer analogue **[^18^F]FDT** that selectively labels lesions (red) in infected organisms (right). Uninfected organisms (left) do not process trehalose and so probe is not retained. **B)** Prior work has established that Ag85s are sufficiently plastic in the substrate scope that they can process, for example, fluorescent analogues of trehalose, allowing them to be metabolically incorporated into the mycobacterial outer membrane *in vitro* for labeling. Fluorescence-based methods are not yet amenable to effective, non-invasive imaging *in vivo*. **C)** Four ^18^F-labeled variants of trehalose were tested in which each of the available hydroxyls (OH-2, 3, 4, 6) were converted in turn to ^18^F. **D)** Enzymatic synthesis of [^18^F]FDT. Three parallel routes (A, B and C, gray) were evaluated for efficiency, rate and yield. Route A: TreT-mediated synthesis was evaluated using enzymes from two different sources (from *T. tenax* or *P. horikoshii*). Both proved functional but gave lower turnover under a range of conditions (see pseudo-single substrate plot for TreT (from *T.* tenax), lower right – See **Supplementary Figure S1** for further details of kinetics). [Reagents and Conditions: 50 mM HEPES, 100 mM NaCl and 10 mM MgCl_2_, pH 7.5]. Route B: OtsAB-fusion enzyme-mediated synthesis was explored to allow use of fewer biocatalysts. A OtsAB fusion protein was constructed but proved to be more difficult to express and less stable under typical reaction conditions [Reagents and Conditions: 50 mM HEPES, 100 mM NaCl and 10 mM MgCl_2_, pH 7.5 C] Route C: Although a three-step, three-enzyme route, Route C proved to be more flexible and reliable. The selectivity of the biocatalysts allowed this to be performed in a convenient one-pot manner. The stability of enzyme components and the ability to vary catalyst amounts to control flux led to its choice over Route B. The higher turnovers and efficiencies led to its choice over Route A (see pseudo-single substrate plot for OtsA, upper right – See **Supplementary Figure S1** for further details of kinetics). [Reagents and Conditions: 50 mM HEPES, 100 mM NaCl and 10 mM MgCl_2_, pH 7.5.]

With the aim of improving detection specificity, other [^18^F]-labeled PET tracers have been investigated for imaging of TB such as 2-[^18^F]fluoroisonicotinic acid hydrazide^15^ and [^18^F]deoxy-fluoro-L-thymidine (FLT)^16^. However, the sensitivity and specificity of these radiotracers is either only similar to (or worse than) [^18^F]FDG and so only can only help to distinguish TB from malignancy when also combined with [^18^F]FDG. Moreover, neither probes are yet readily accessible from common precursors; both require specialist generation of radioisotopes (e.g. [^18^F]fluoride via cyclotron-mediated proton (^1^H) irradiation of H_2_^18^O) combined with specific chemical technologies (e.g. appropriate synthetic laboratories and methods).

Trehalose, a nonreducing disaccharide in which the two glucose units are linked in an α,α-1,1-glycosidic linkage (**Figure 1a**) has, in the last decade, gathered interest as an agent for selectively imaging TB. Trehalose is found in *Mtb*, especially in the outer portion of the mycobacterial cell envelope – primarily as its corresponding glycolipids trehalose monomycolate (TMM) and trehalose dimycolate (TDM). These glycolipids appear to play a critical structural, and perhaps pathogenic role, as an essential cell wall component,^17,18^ as has long been noted (first referred to as ‘cord-factor’ in the 1950s^19^). The biosynthesis of trehalose mycolates is catalyzed specifically in *Mtb* by a family of abundant TB-specific, membrane-associated enzymes – the Antigen 85 proteins (Ag85a, Ag85b and Ag85c).^20^

The lack of naturally-occurring trehalose in mammalian hosts as well as the uptake of exogenous trehalose by *Mtb*^21^ has suggested that trehalose-based probes could function as both a highly specific and sensitive reporters (**Figure 1a**). Prior work has established that the Ag85 family enzymes (Ag85s) are somewhat plastic in the substrates that they can process – not only can exogenous trehalose be processed by Ag85s but also analogues of trehalose. In this way, they can be metabolically incorporated into the mycobacterial outer membrane.^21^ This allowed selective fluorescent labeling and monitoring of *Mtb* in vitro, not only in bacilli but also within infected mammalian macrophages using reagents such as FITC-Tre (**Figure 1b**).^21^ More recently, elegant design has created yet more powerful fluorescent variants, such as the fluorogenic QTF,^22^ and FRET-TDM^23^ and solvatochromic DMN-Tre^24^ (**Figure 1b**), some of which enable detection of *Mtb* detection in sputum,^24^ as well as photosensitized variants for possible photodynamic therapy.^25^ However, fluorescence-based methods are not yet amenable to effective, non-invasive clinical *in vivo* imaging and, to our knowledge, no trehalose-based probe or method has yet demonstrated efficacy *in vivo*. Here we show that one of the simplest ^18^F-analogs of trehalose, 2-[^18^F]fluoro-2-deoxy-trehalose **[^18^F]FDT** can be generated as an *in vivo* TB-reporter using one-pot, automatable, pyrogen-free, chemoenzymatic synthesis from readily-available [^18^F]FDG in a validated manner that does not require specialist expertise (**Figure 1c**).^26,27,28^ Toxicological and preclinical testing in diverse species shows that this now creates access to a safe probe that is effective and specific in multiple preclinical models both to visualize TB lesions and to monitor their treatment non-invasively. **[^18^F]FDT** is therefore now suggested as suitable candidate for clinical use, enabled by a distributable synthetic strategy that could help lower global health technology imbalances.

## Results

### Comparison of four candidate ^18^F-fluoro-deoxy analogs of trehalose as probes identifies [^18^F]FDT as a likely radiotracer reporter of TB

To test the effectiveness of different minimal alterations at different sites in the trehalose scaffold (**Figure 1c**), we synthesized four radiolabeled ^18^F-fluoro-analogs of trehalose. We aimed to use complementary, straightforward chemical and enzymatic approaches that were initially developed via semi-analogous ‘cold’ routes (see **Supplementary Figure S2**). In each fluoro-trehalose analog a single hydroxyl at each of the four different sites in the trehalose scaffold was replaced by a fluoro group – specifically at positions 2-, 3-, 4 and 6-giving the corresponding deoxy-[^18^F]fluoro analogs: **[^18^F]FDT**, 3-[^18^F]FDT, 4-*epi*-[^18^F]FDT and 6-[^18^F]FDT (**Figure 1c**).

In our design (**Figure 1c**) of these minimally-altered trehalose analogues, we considered not only tolerance for the PET-tracing label (due to minimal structural changes) for co-opted processing by *Mtb* but also we were mindful of the potential inhibitory properties that such compounds might have upon putative host degrading enzymes^29^ – 2-fluoro-2-deoxy-sugar substrates, for example, have been elegantly shown to act as irreversible, mechanism-based inhibitors of corresponding sugar hydrolases.^30^

Specifically, in brief, 4-*epi*-[^18^F]FDT and 6-[^18^F]FDT novel radiochemical syntheses used nucleophilic fluorination protocols that generated product from corresponding triflate precursors in good yields and with high purity (**Supplementary Fig. S3 A and B, Fig. S4**). As such, these synthetic approaches were strategically similar to typical syntheses of, for example, commercially-available [^18^F]FDG, where aqueous [^18^F]-fluoride is used as the source of ^18^F^31^. Protocols for **[^18^F]FDT**, 3-[^18^F]FDT utilized enzymatic methods (see the **Supplementary Methods** and following sections for details).

Whilst the syntheses of **[^18^F]FDT**, 4-*epi*-[^18^F]FDT and 6-[^18^F]FDT proved sufficiently rapid (complete within one half-life of ^18^F, t_1/2_ ∼110 min), that of 3-[^18^F]FDT was slower (consistent with results for ‘cold’ analogues) and less efficient (3-[^18^F]FDT, 10% conversion (n = 2), 60 min to 4h). Moreover, whilst 4-*epi*-[^18^F]FDT and 6-[^18^F]FDT could be prepared from [^18^F]-fluoride used at a later stage (substitution then deprotection, see **Supplementary Figs S2-4**), 3-[^18^F]FDT required the prior preparation of deprotected 3-[^18^F]fluoro-3-deoxy-glucose before enzymatic processing, which was itself relatively inefficient^32^. The particular difficulties in the synthesis of 3-[^18^F]FDT (length of reaction times, poor conversion and need for early use of radionuclide) led to low specific activities and so this candidate was dismissed at an early stage.

The metabolic stabilities of the remaining three analogs, **[^18^F]FDT**, 4-*epi*-[^18^F]FDT and 6-[^18^F]FDT were tested. Although, human trehalose-degrading trehalase activity is low and restricted primarily to the brush-border of the kidney where it is thought to be GPI-anchored,^29,33^ it can be elevated in some diseases,^29^ therefore we also tested degradation of the analogs *in vitro* using high concentrations of mammalian (porcine) trehalase (0.3 units/mL, 10 mM substrate, **Supplementary Figure S5**). Under conditions that led to complete degradation of trehalose, ‘cold’ analogs [**^19^F]FDT**, 4-*epi*-[^19^F]FDT showed little or no degradation, whereas 6-[^19^F]FDT showed low levels (14%).

Finally, putative degradation was also probed directly *in vivo* using both *Mtb*-infected and naïve rabbits (see **Supplementary Information**). Consistent with the *in vitro* observations, radiotracer 6-[^18^F]FDT was less stable than [**^18^F]FDT** and 4-*epi*-[^18^F]FDT; 6-[^18^F]FDT was entirely metabolized to 6-[^18^F]FDG within 90 min post-injection, (see **Supplementary Figure** (**Supplementary Figure S3E,F**)). Whilst 4-*epi*-[^18^F]FDT proved more stable, some metabolism (albeit slower) was observed (30% after 3h (**Supplementary Figure S3C,D**). However, consistent with design (**Figure 1c**), [**^18^F]FDT** showed no apparent degradation (see below).

Taken together, the rapid degradation of 6-[^18^F]FDT and both the partial degradation and less efficient (lower RCY and a need for more stringent purification) synthesis of 4-*epi*-[^18^F]FDT led to their dismissal as candidate tracers; [**^18^F]FDT** was evaluated further.

### Comparison of synthetic routes suggests a flexible One-Pot, biocatalytic route to **[^18^F]FDT**

Three parallel strategies were envisaged for the synthesis of [**^18^F]FDT** (**Figure 1d**) and these were tested in the synthesis of [^19^F]FDT. We aimed from the outset to devise routes that would ultimately utilize [^18^F]FDG, as this is the most readily-available organic source of ^18^F, even to those without specialist sources of ^18^F (e.g. cyclotron facilities). Global access via commercial sources would therefore, in principle, allow the development of synthesis that could implemented in many locations.

[^18^F]FDG is typically supplied in aqueous solution and so all routes adopted a protecting-group-free approach that would not only obviate the need for additional (e.g. protection-deprotection) steps, but, through the use of biocatalysts, allow the use of such solutions directly. Given the often highly (chemo-, regio- and stereo-) selective nature of biocatalysts we envisaged that all three routes would have potential for application in a ‘one-pot’ operation for ease-of-use. Four biocatalyst systems were evaluated for mediating the critical formation of the α,α-1,1-*bis*-glycosidic linkage that is found in [**^18^F]FDT**. Due to the nature of its dual *cis*-1,2-linkage,^34^ this is a particularly difficult linkage to form and control selectively using chemical methods, further highlighting another potential advantage of the use of biocatalysis. Two homologues of the, so-called, trehalose glycosyltransferring synthase enzyme, TreT,^35^ (from *T. tenax*^36,37^ and *P. horikoshii*^38^) were tested via Route A; and two variant constructs containing the trehalose 6-phosphate synthase domain OtsA,^39^ one in Route B as a fusion to the dephosphorylating enzyme OtsB and one as a single enzyme construct via Route C.

First, for initial syntheses, all enzymes were expressed in and purified from *E. coli*. The yield for OtsA enzyme as an individual construct was far superior to the OtsAB fusion construct and/or TreT homologues. OtsB from *E. coli* did not express in the soluble fraction but could either be purified from inclusion bodies or its solubility improved through the addition of a fused-*N*-terminal-MBP-His-tag (see **Supplementary Information and Supplementary Figures S6-S8**).

Next, we determined the kinetic parameters of all of the enzyme systems (**Supplementary Figure S1** and **Table 1**) using a variety of complementary (continuous or endpoint; coupled-enzyme^40^, spectrophotometric, HPLC or ^1^H NMR) methods to monitor substrate consumptions and/or product formation, according to conditions. Again, individual OtsA and OtsB constructs proved superior catalysts (**Table 1**): not only to the corresponding OtsAB fusion but also to the two TreT enzymes with OtsA catalytic efficiencies some >2-fold to 4-fold higher, as judged by k_cat_/K_M_. The relative efficiency of OtsB was yet higher (∼4-fold to 10-fold higher even than OtsA) allowing its use in lower concentration, further highlighting the advantage of this separate construct (see below).

**Table 1:**
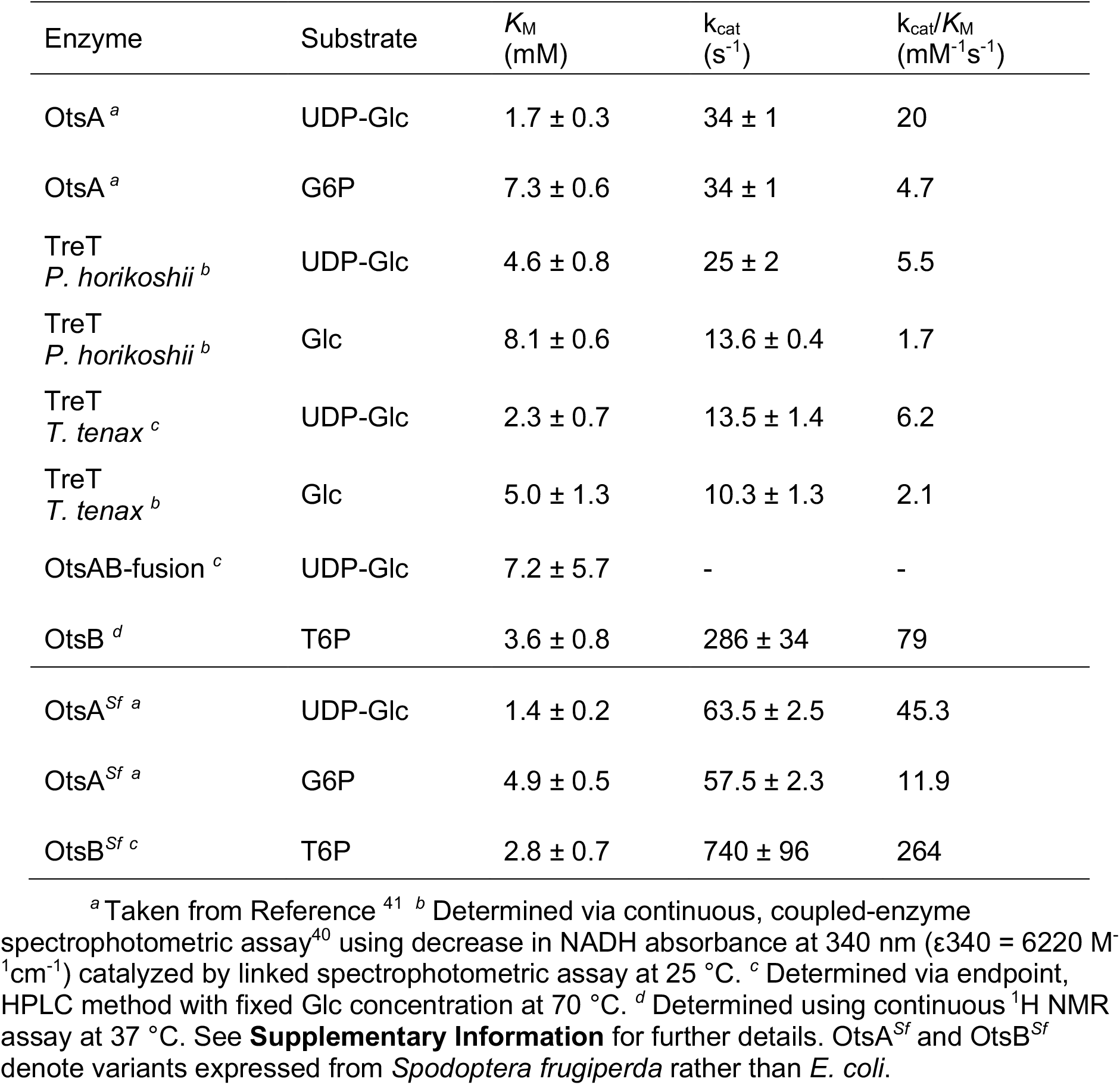
Kinetic Parameters of Enzymes used in [^19^F]FDT Synthesis.

| Enzyme | Substrate | $K_M$<br>(mM) | $k_{cat}$<br>(s<sup>-1</sup>) | $k_{cat}/K_M$<br>(mM<sup>-1</sup>s<sup>-1</sup>) |
| --- | --- | --- | --- | --- |
| OtsA<sup>a</sup> | UDP-Glc | 1.7 ± 0.3 | 34 ± 1 | 20 |
| OtsA<sup>a</sup> | G6P | 7.3 ± 0.6 | 34 ± 1 | 4.7 |
| TreT<br>*P. horikoshii*<sup>b</sup> | UDP-Glc | 4.6 ± 0.8 | 25 ± 2 | 5.5 |
| TreT<br>*P. horikoshii*<sup>b</sup> | Glc | 8.1 ± 0.6 | 13.6 ± 0.4 | 1.7 |
| TreT<br>*T. tenax*<sup>c</sup> | UDP-Glc | 2.3 ± 0.7 | 13.5 ± 1.4 | 6.2 |
| TreT<br>*T. tenax*<sup>b</sup> | Glc | 5.0 ± 1.3 | 10.3 ± 1.3 | 2.1 |
| OtsAB-fusion<sup>c</sup> | UDP-Glc | 7.2 ± 5.7 | - | - |
| OtsB<sup>d</sup> | T6P | 3.6 ± 0.8 | 286 ± 34 | 79 |
| OtsA<sup>Sf a</sup> | UDP-Glc | 1.4 ± 0.2 | 63.5 ± 2.5 | 45.3 |
| OtsA<sup>Sf a</sup> | G6P | 4.9 ± 0.5 | 57.5 ± 2.3 | 11.9 |
| OtsB<sup>Sf c</sup> | T6P | 2.8 ± 0.7 | 740 ± 96 | 264 |

<sup>a</sup> Taken from Reference <sup>41</sup> <sup>b</sup> Determined via continuous, coupled-enzyme spectrophotometric assay<sup>40</sup> using decrease in NADH absorbance at 340 nm ( $\epsilon_{340} = 6220 \text{ M}^{-1}\text{cm}^{-1}$ ) catalyzed by linked spectrophotometric assay at 25 °C. <sup>c</sup> Determined via endpoint, HPLC method with fixed Glc concentration at 70 °C. <sup>d</sup> Determined using continuous <sup>1</sup>H NMR assay at 37 °C. See **Supplementary Information** for further details. OtsA<sup>Sf</sup> and OtsB<sup>Sf</sup> denote variants expressed from *Spodoptera frugiperda* rather than *E. coli*.

These catalytic efficiencies had a direct bearing on synthetic utility. In comparative 10 mM reactions: via Route A (**Figure 1D**) TreT from *P. horikoshii* performed better (40% conversion over 2 h) than the TreT from *T. tenax* (10% conversion over 2 h); *P. horikoshii* TreT also dipslayed reversibility under certain conditions that would contaminate product FDT by converting it back to starting FDG. Via Route B using OtsAB, synthesis failed (even over prolonged periods). In striking contrast, via Route C using OtsA and OtsB full conversion was observed within 45 min (**Supplementary Figure S9**).

Following these evaluations Route C (**Figure 1**) was selected over both Routes A and B on the basis of higher biocatalyst expression yields, flexibility of enzyme usage (differential loadings), stability and superior kinetic parameters, which, together, gave rise to more efficient synthesis in our hands.

### Creation of a pyrogen-free synthetic biocatalyst system enables a safe route to **[^18^F]FDT**

Pyrogen or endotoxin presence due to lipopolysaccharide contamination caused, for example, by carryover from *E. coli* production is prominent and of serious concern in any potential clinical method^42^. Whilst we were able to successfully utilize extensive, repeated cyclical purification coupled with pyrogen-monitoring using *Limulus* amoebocyte lysate activity measurements^43^ to demonstrate low pyrogen production of [^19^F]FDT (**Supplementary Table S1**), it has been argued that depyrogenation methods cannot be fully effective^42,44^.

We therefore developed an expression system free of endotoxin based on the use of *Spodoptera frugiperda* (Sf9) cell culture. An appropriate sequence was designed based on the pFastbac vector system; the coding sequences of *E. coli* OtsA (CAA48913.1) and *E. coli* OtsB (CAA48912.1) were refactored according to *S. frugiperda* MNVP codon usage. Analysis of the resulting gene sequences for *E. coli* and *S. frugiperda* constructs confirmed essentially identical protein gene product primary sequence (**Supplementary Figure S10**). This allowed ready implementation of a protein production workflow **Supplementary Figure S11**) using infectious baculovirus generated through transposon-mediated insertion^45^ (so-called Bac-to-Bac system). Sf9-culture resulted in good yields of proteins OtsA*^Sf^* and OtsB*^Sf^* (∼ 20 mg/L of culture for OtsA*^Sf^*; ∼15 mg/L of culture for OtsB*^Sf^*, **Supplementary Figure S12**) that were essentially endotoxin-free (< 0.1 EU.mL^-1^). This was comparable or superior to those found from *E. coli* expression of OtsA or OtsB and, indeed, of TreT and OtsAB fusion enzymes (both TreT enzymes ∼2-3 mg/L, OtsAB fusion ∼0.1 mg/L of culture). Both OtsA*^Sf^* and OtsB*^Sf^* proved stable up to 37 °C (**Supplementary Figure S13**). Moreover, the kinetic parameters of OtsA*^Sf^* and OtsB*^Sf^* (from Sf9 insect cells) also proved superior to those expressed from *E. coli*^41^ (2-to-3-fold more efficient as judged by k_cat_/*K*_M_, **Table 1**). Finally and importantly, we were able to readily scale enzyme expression in Sf9 up to 10 L in one batch for both, thereby allowing corresponding production of >100 mg of either of the OtsA*^Sf^* and OtsB*^Sf^*enzymes.

### Development of an efficient, scaleable radiosynthesis provides **[^18^F]FDT** for pre-clinial testing

With a putative biocatalytic system in hand, we next sought to optimize its application in the synthesis of [^19^F]FDT as an isotopologue model for **[^18^F]FDT** (**Figure 2**). Reaction parameters were varied (see **Supplementary Methods**), including substrate and enzyme concentrations as well as duration of reactions. First, independently-variable, biocatalyst concentrations further verified the choice of flexible Route C (**Figure 2A**) and allowed optimization of relative [E]_0_ consistent with their kinetic parameters (**Table 1**) to allow maximal flux; this identified of 18.2 µM OtsA*^Sf^* and 6.1 µM OtsB*^Sf^* (**Supplementary Figure S14,15**) as efficient. Second, variation of acceptor substrate [^19^F]FDG concentration in the presence of fixed, maintained amounts of donor substrate UDP-Glc revealed more rapid reaction progress in the presence of equimolar or excess donor, consistent with both a highly efficient glycosylation reaction and also inhibition that could be minimized through control of conditions (**Figure 2B**); variation of donor concentration under these constraints then identified concentrations of ∼130-260 mM as optimal for later reactions.

**Figure 2.**
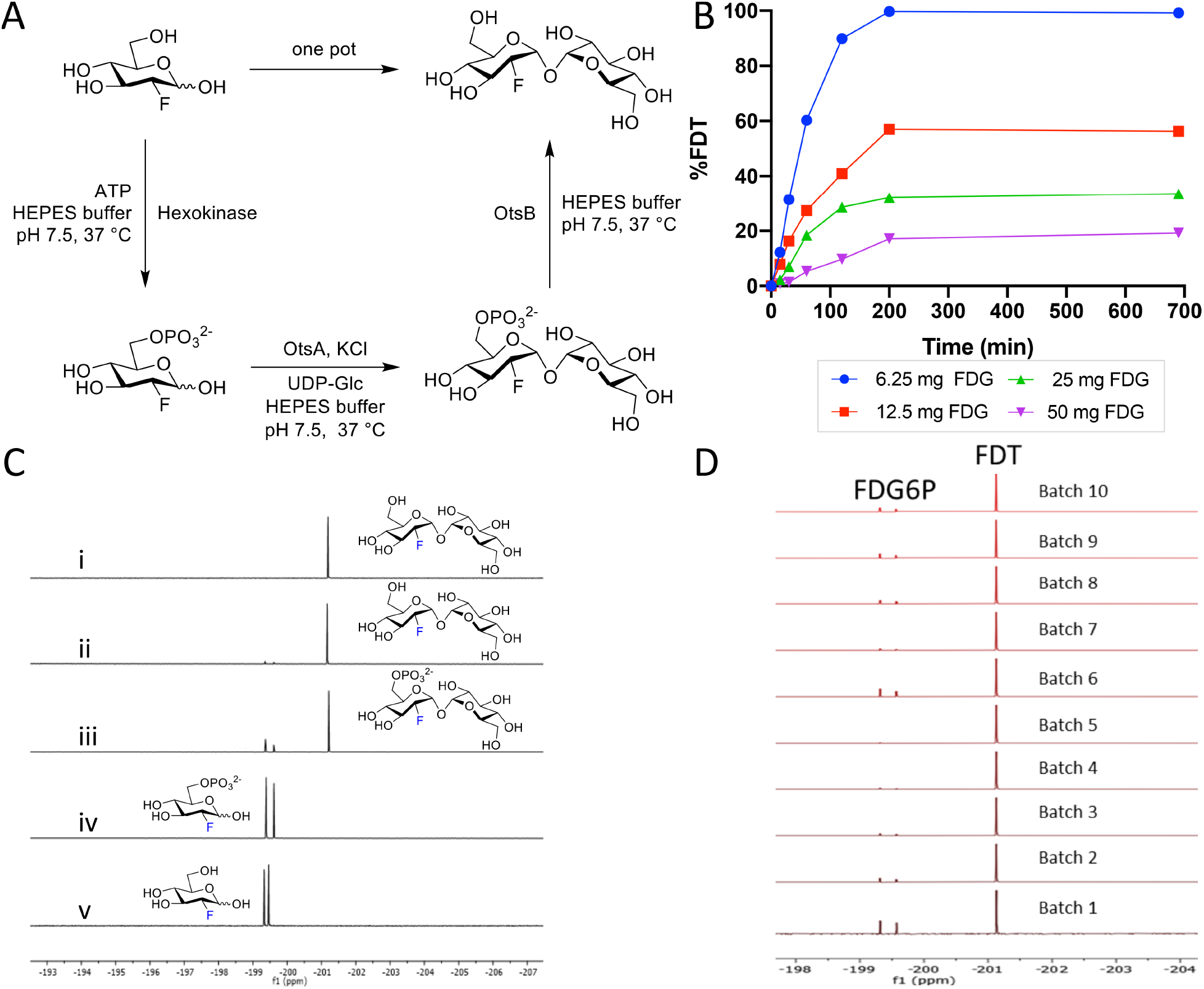
Reaction Optimization, Scale-up and Batch Synthesis in the Development of an Efficient Scaleable One-Pot Synthesis. **A)** Chosen Route C (from **Figure 1**) was tested in a one-pot format [^19^F]FDT using pyrogen-free enzymes OtsA*^Sf^* and OtsB*^Sf^*. **B)** Example reaction optimization with fixed donor sugar Glc-UDP [30 mM] at different acceptor substrate quantities (6.25, 12.5, 25 and 50 mg) of [^19^F]FDG reactions were monitored in real time by calibrated ^19^F NMR to optimize reaction. **C)** Direct reaction monitoring of one-pot, 3-step [^19^F]FDT synthesis from [^19^F]FDG by ^19^F-NMR. NMR spectra: **(i)** ^19^F NMR spectrum of purified [^19^F]FDT; **(ii)** Crude ^19^F NMR spectrum of [^19^F]FDT with small amount of deoxy-fluoro-G6P (**[^19^F]-1**) still present; **(iii)** Crude ^19^F NMR spectrum of intermediates **[^19^F]-1** and **[^19^F]-2**; **(iv)** Crude ^19^F NMR spectrum of **[^19^F]-1** from [^19^F]FDG conversion; and **(v)** reference sample of starting material [^19^F]FDG. **D)** Representative, ^19^F-NMR spectra of crude reaction mixtures containing [^19^F]FDT from repeated batches synthesised by 3-enzymes-3-steps, one-pot syntheses, prior to purification.

With these conditions in hand, we scaled low mg reactions to a scale of 100s of mg (from 100 mg [^19^F]FDG). Reaction progress through each step in a ‘one pot’ operation was monitored by the measurement of the crude ^19^F-NMR spectra from reaction mixtures (**Figure 2C**). This in turn allowed the testing of repeated batches (**Figure 2D**); we reasoned that such a batch scale would prove sufficient for a dose in man (see below for further details) and was therefore a representative test batch. These experiments revealed that at higher scale steps one and three (**Figure 2A**) proved slightly more efficient than step two with the occasional observation of small amounts of unconverted deoxy-fluoro-G6P ([^19^F]-**1**) in some batches (**Figure 2D**) – advantageously, by virtue of its charge, if present [^19^F]-**1** (and similarly [^18^F]-**1**, see below) could be readily removed both during manual and automated syntheses (see below), further confirming the advantage of Route C. In this way we could readily, consistently and repeated access [^19^F]FDT at 100 mg scales through one-pot biocatalytic syntheses; allowing the ready synthesis of grams of [^19^F]FDT that when characterized (including via ^1^H-, ^13^C-, ^19^F-, ^23^Na-NMR, ^1^H qNMR, microanalysis and LC detected by multiple methods – see **Supplementary Figures S16-29**) complied fully with stringent measures of purity (> 98% organic, with NaCl as the only co-eluent (∼10-12%)) for subsequent *in vivo* toxicity testing (see below and **Supplementary Methods**).

Next, with ‘cold’ syntheses in hand, we applied essentially analogous methods for the one-pot syntheses of **[^18^F]FDT**: first manually and then in an automated fashion that might prove suitable for GMP synthesis in Phase 1 studies. Using commercially available aqueous solutions of [^18^F]FDG with activity of 5-15 mCi we observed full conversion to **[^18^F]FDT** in 60 min. Purification was effected simply by adding ethanol to precipitate protein, filtration (5 µm filter) and then purification of the resulting filtrate using ion-exchange (‘Luna’ NH_2_, see **Supplementary Methods**). Product **[^18^F]FDT** was eluted with ethanol in a non-decay-corrected radiochemical yield (RCY) over three steps of 34 ± 14 % (n = 5) and with a radiochemical purity >99% (**Supplementary Figure S30**) and an overall reaction time of 60 minutes.

To test possible, ‘real world’ application, these radiosyntheses were repeated from different batches of starting [^18^F]FDG and also used directly in *in vivo* studies (see below). Depending on starting material concentration, activity and condition, overall production times for **[^18^F]FDT** of 30-120 min were tested. For example, reaction times could be reliably reduced from 60 min to 30 min by increasing concentration of enzymes, (OtsA*^Sf^*, OtsB*^Sf^* and hexokinase to 9, 15 and 45 μM, respectively) to ensure >99% conversion to provide comparable quantities and the same high purities of **[^18^F]FDT**. No detectable, residual protein from the reaction was observed in the final product, as tested by Western blot (< 1 μg/reaction, **Supplementary Figure S31**).

Next, after this successful manual standardization and variation, a fully automated synthesis was performed in a GE Tracerlab FX-N module (**Supplementary Figure S32)**. Even higher, overall decay-corrected radiochemical yield (**Supplementary Table S2**) was observed with higher amounts of starting [^18^F]FDG; under typical conditions (100 mCi [^18^F]FDG), 41 ± 4 % (n=2) non-decay corrected yield of **[^18^F]FDT** was obtained in 50 min. The identity of product **[^18^F]FDT** was confirmed via co-injection of an authentic standard using LC-MS; radiochemical purity was >98%, determined by HPLC (**Supplementary Figures S33, S34**). Specific activity, assessed using an adapted enzymatic assay^46^ of decayed samples (see **Methods**), revealed that, as expected, activity was essentially dependent on [^18^F]FDG source – useful activities could be routinely achieved (average 0.21 ± 0.08 µmol **[^18^F]FDT** at specific activity of 69 ± 26 mCi/mg (23.6 ± 8.8 Ci/mmol) at the end of synthesis and formulation, see **Supplementary Methods**).

### [^19^F]FDT and **[^18^F]FDT** are metabolically stable

*In vitro* [^19^F]FDT in human plasma showed no degradation (**Supplementary Figure S35**). For evaluations *in vivo,* solutions of [^19^F]FDT in saline (100 µL of 50 µM and 500 µM) were injected into mice, and plasma subsequently analyzed (see **Supplementary Methods**; both LC-MS and ^19^F-NMR spectroscopy allowed detection of [^19^F]FDT and any putative degradation (**Supplementary Figure S36**). Together, these confirmed stability of [^19^F]FDT both in plasma *in vitro* and metabolically *in vivo*.

Finally, this stability was confirmed for **[^18^F]FDT** through direct assessment of radiochemical purity maintained >95% in both blood and urine of rabbits (**Supplementary Figure S3G**) (see also above and below). As noted above, this contrasted strongly with the significant metabolism of both 6-[^18^F]FDT and 4-*epi*-[^18^F]FDT.

### Radiotracers label mycolate lipids in Mtb-infected tissues

Although we have previously shown that FITC-Tre (**Figure 1b**) can be incorporated into the cell surface of *Mtb* in cultured macrophages^21^, its use for non-invasive *in vivo* imaging is not possible and indeed no current, validated probe exists for this purpose. To determine such utility, we tested if **[^18^F]FDT** incorporation was detectable within complex, tubercular lesions within the practical imaging timeframes allowed by the half-life [^18^F] (t_1/2_ ∼110 min).

First, mouse lung tissue was mixed *ex vivo* with *Mtb*-infected lung homogenate (10^6^ *Mtb*/mL) and **[^18^F]FDT** added and incubated at 37 °C for 60 min. Upon extraction (chloroform) radio-TLC (**Supplementary Figure S3F**) revealed formation of corresponding, radio-labeled trehalose monomycolates and dimycolates (based on R_f_ and comparison to authentic standards of TMM and TDM), consistent with the successful formation of [^18^F]TMM and [^18^F]TDM, respectively. Furthermore, consistent with a common, conserved mechanism, when the caseum from a large tubercular cavity in diseased New Zealand White rabbits that had been probed with 4-*epi*-[^18^F]FDT (8 mCi, see **Supplementary Methods**) was excised from the lung, the corresponding lipids extracted with chloroform from PBS homogenate also correlated with mycolate standards, when observed by radio-TLC (**Supplementary Figure S3F**). Together these data suggested that FDT-based probes could not only act as probes that label mycobacterial lipids, in a manner consistent with the *Mtb*-selective mechanism outlined in **Figure 1A**, but could do so within practical imaging timeframes.

### [^18^F]FDT is a specific, consistent Mtb-radiotracer in vivo with uptake that correlates with bacterial burden in a non-human primate TB model

A useful PET probe for diagnosing tuberculosis should have minimal signal in the absence of *Mtb* infection, particularly in the lung where the majority of infections occur. Importantly, in a non-human primate (NHP) marmoset TB model^47^ **[^18^F]FDT** displayed low background signal in the lung of a naïve marmoset compared to ∼45-day *Mtb*-infected marmoset (**Figure 3A, 3B, 3C**). To confirm that this enhanced signal was specific to uptake of tracer, rather than, for example, non-specific uptake (proportional to tissue density) or normal distribution, ‘pre-blocking’ experiments were conducted with both trehalose and ‘cold’ [^19^F]FDT (administered 1 h and 5 minutes) prior to the use of **[^18^F]FDT** radiotracer dose. Consistent with specificity and mode-of-action, these reduced average uptake of **[^18^F]FDT** into lesions by 40% (**Figure 3D, 3E**, ∼40% reduction **F**). Since non-specific uptake resulting from transit or non-bound normal accumulation of the radiotracer should not be displaced by competition with blocker / ‘cold’ compound, a reduction in PET signal is consistent with specific binding.

**Figure 3.**
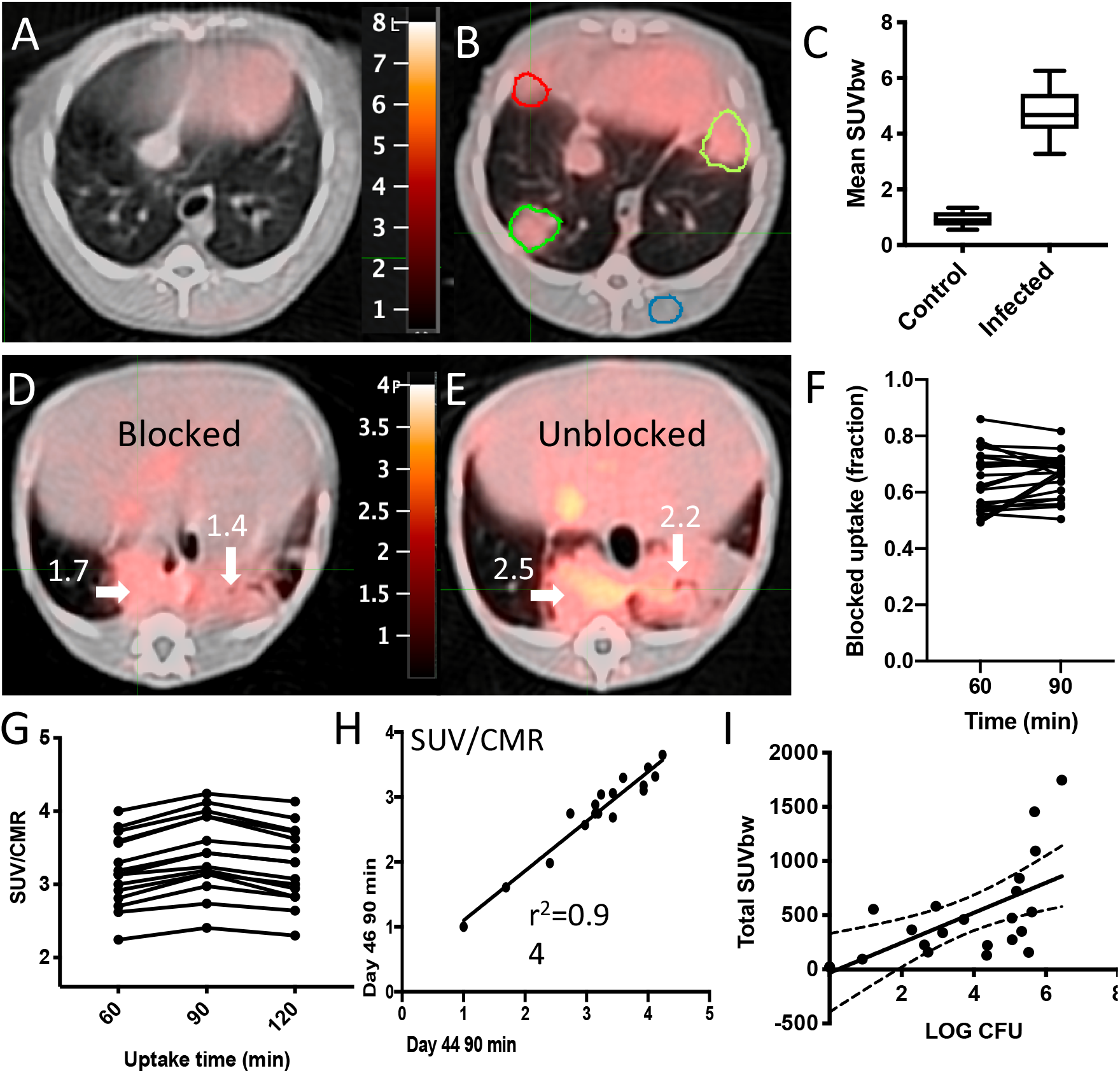
Pulmonary [^18^F]FDT PET uptake into Tuberculous Lung Lesions is Reproducible within 90 min with Correlated Signal in Lesions with Higher Bacterial Loads. **A) [^18^F]FDT** PET/CT scan of a naïve marmoset lung (administered 1 mCi) **[^18^F]FDT** and imaged 90 min post-injection (transverse view). **B) [^18^F]FDT** PET/CT scan of a representative, Mtb-infected marmoset lung showing lesions outlined in green and red (transverse view). The iliocostalis muscle was used for normalization of PET uptake (region outlined in blue). The target dose was 2.2 mCi/kg (∼1 mCi). **C)** The **[^18^F]FDT** PET signal from Mtb-infected marmoset lungs is significantly higher than the signal from uninfected (control) marmoset lungs. The mean Standard uptake values (SUVbw) are represented as boxplots where the central bars represent the median. At least 4 lung regions of interest (ROI) were measured from 3 infected and 2 uninfected marmosets (p<0.0001). **D)** and **E)** Transverse images of **[^18^F]FDT** PET uptake scans for unblocked (**D**) and blocked (**E**) uptake in the same infected marmoset. **[^18^F]FDT** uptake was blocked by administering excess blocker, split into two administrations 1 h and 5 min prior to radiotracer administration. **F)** Reduction of **[^18^F]FDT** uptake in marmosets administered cold [^19^F]FDT blocker prior to being administered **[^18^F]FDT** (1.2 mCi) compared to when administered **[^18^F]FDT** alone. Animals (n = 22) were randomized as to the order of blocked scan; two days later the same animals were imaged with the treatments reversed. Uptake was measured in images collected 60 min and 90 min after probe administration. In both cases the animal receiving [^19^F]FDT blocking dose showed significantly lower accumulation of **[^18^F]FDT** into tubercular lesions. **G)** Comparison of **[^18^F]FDT** (1.0 mCi) lesion uptake (SUV/CMR) at 60, 90, and 120 mins post injection (lesions from two infected, antibiotic-treated marmosets) demonstrates that signal does not increase after 90 minutes. **H) [^18^F]FDT** uptake (SUV/CMR) is reproducible (r^2^ = 0.94), as illustrated by comparison of individual lesion SUV/CMR values from marmosets with imaging repeated two days apart (44 dpi and 46 dpi). **I) [^18^F]FDT** uptake into tubercular lesion tends to increase with higher mycobacterial loads (p = 0.001, ρ = 0.64). The total SUV of each lesion was compared with mycobacterial colony forming units measured (log CFU) in the lesions from two marmosets.

Optimal imaging time was determined to be 90 minutes post-dose (**Figure 3G**); there was no additional improvement in signal-to-noise at 120 minutes scan. Reproducibility of **[^18^F]FDT** uptake was assessed by measuring lesion uptake (SUV) in animals scanned 48 hours apart (44 and 46 days-post-infection (dpi)), sufficient time to allow full tracer clearance but short enough to minimize possible infection progression (**Figure 3H**); consistent **[18F]FDT** uptake (SUVs) was observed confirming reproducible uptake.

Finally, to determine whether lesion uptake of **[^18^F]FDT** correlated with disease, we determined bacterial burden at sites that were imaged by **[^18^F]FDT**. Post-necropsy excision of lesions (n = 21) revealed that **[^18^F]FDT** uptake into individual lesions (SUV per lesion) was significantly correlated (p = 0.001, π = 0.64) with the number of culturable Mtb bacteria (CFU) from each lesion (**Figure 3I**).

### [^18^F]FDT uptake and radiotracer signal differs from that of [^18^F]FDG in Mtb-infected models

We reasoned that this designed, mode-of-action of **[^18^F]FDT** that correlated well with mycobacterial burden might complement the differing mode-of-action shown by [^18^F]FDG uptake and hence reveal additional, valuable physiological information when used in tandem. In particular [^18^F]FDG in TB appears to label immune cells^8,9^ that are highly metabolically-active.

To the test the potential for different labeling patterns, we directly compared **[^18^F]FDT** and [^18^F]FDG scans collected sequentially in *Mtb*-infected marmosets (n = 3), with sufficient intervening time to clear (**Figure 4**). These revealed clearly differential labeling of lesions and pooling in tissue regions with different characteristics. Strikingly, no quantitative correlation was observed for the different labeling patterns either via identification of regions-of-interest or use of total glycolytic activity. Notably little-to-no overlap was seen for regions of most intense radioactive signal.

**Figure 4.**
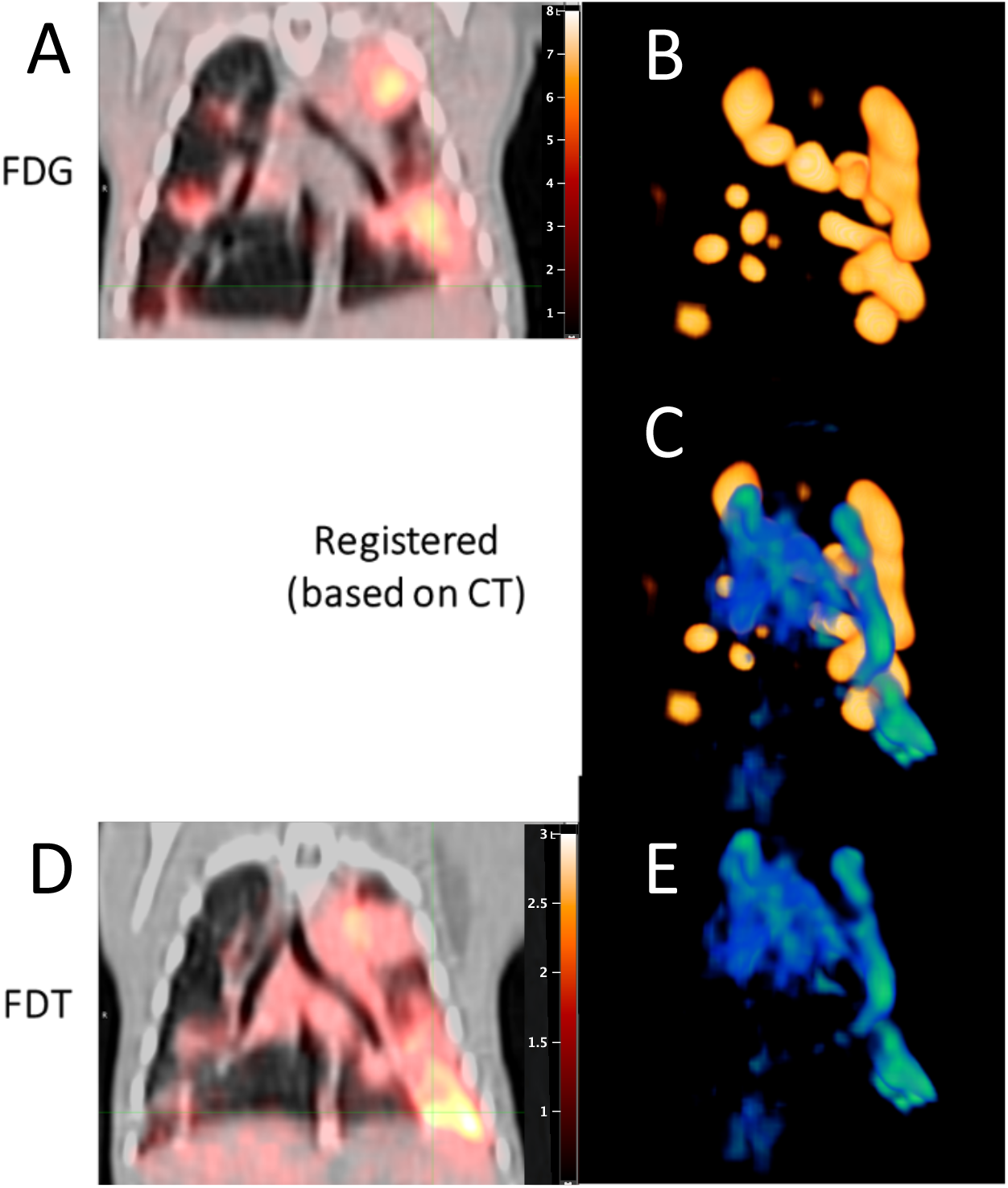
Differential Uptake and Labeling of Tubercular Lesions by [^18^F]FDT and [^18^F]FDG in *Mtb*-Infected Marmosets. Serial [^18^F]FDG (0.8 mCi) (**A** and **B**) and **[^18^F]FDT** (1.1 mCi) (**D** and **E**) scans of the same infected marmoset were collected at 90 minutes post injection and co-registered based on the PET/CT shown in (**A**) and (**D**), respectively. In (**C**) the three-dimensional maximum intensity projection (MIP) of the FDG signal in the lung (colored shades of yellow-to-gold (**B**)) was overlaid with the three-dimensional MIP of the **[^18^F]FDT** signal (colored shades of green-to-blue (**E**), green showing most intense labeling) to indicate the differences in the regions of intense labeling. Together these show clear differences in regions of intense radioactive signal indicating that the two probes are pooling in tissue regions with different characteristics.

### [^18^F]FDT allows direct monitoring of treatment that correlates with reduction in bacterial load

One of the most significant challenges facing current TB diagnosis and treatment programmes is a source of reliable information on the disease burden in patients ^48^. This is exacerbated by the often-prolonged treatment regimens that may continue over months and even years. Compliance with continual treatment is vital and direct feedback on progress of treatment would prove invaluable not only to the clinician but also patient ^49,50^. Moreover, direct measurement of the *in vivo* effectiveness of new therapeutics would provide a powerful new tool in their clinical development ^51-53^.

We therefore tested whether **[^18^F]FDT** would report effectively on response to a representative *Mtb* therapy. Marmosets were treated for 4 weeks (five days per week) with first-line HRZE (combined isoniazid (H), rifampicin (R), pyrazinamide (Z) and ethambutol (E)) therapy and serial images were taken using both **[^18^F]FDT** (**Figure 5E and F**) along with [^18^F]FDG as a comparator (**Figure 5A and B**). Notably, [^18^F]FDG showed mixed, refractory and inconsistent uptake as judged by both SUVs and total lesion glycolysis (TLG) (**Figure 5C and D**), despite the fact that effective treatment was confirmed by low bacterial burden after treatment (post-necropsy logCFUs of 2.6 and 2.8, respectively). By striking contrast, **[^18^F]FDT** scans showed a significant reduction (mean 33 ± 13% SD, p <0.01) in SUV signal and a lower total **[^18^F]FDT** uptake (**Figures 6G and H**) during treatment consistent with the determined drop in disease burden. Together, these data suggest that **[^18^F]FDT** could provide a more accurate probe of disease treatment.

**Figure 5.**
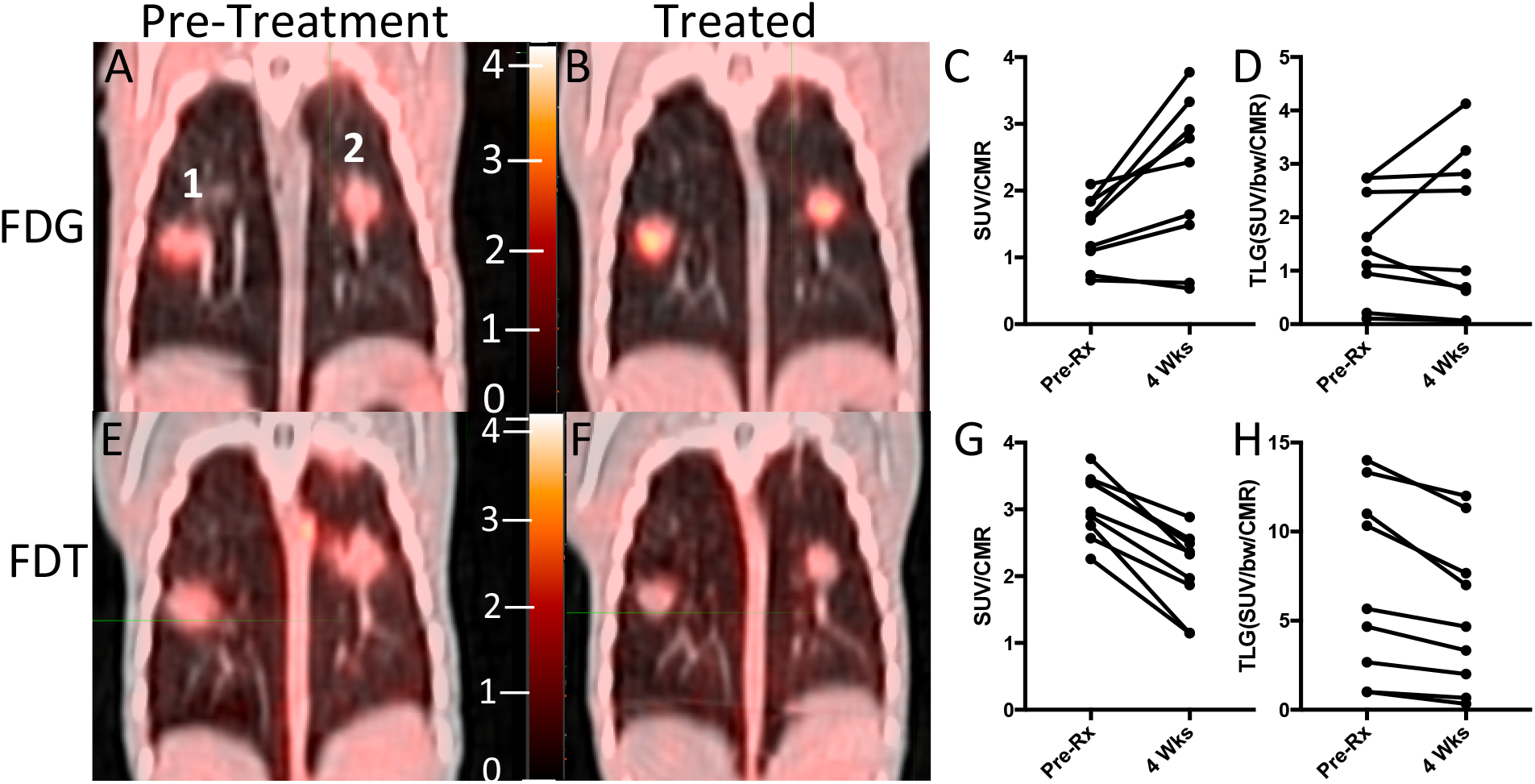
[^18^F]FDT Scans of an *Mtb*-infected Marmoset Correlate with 4-week HRZE Treatment. **A)** Pretreatment scan 60 min post-injection with [^18^F]FDG PET (0.7 mCi) shows two lesions (indicated as 1 and 2). **B)** Marmoset imaged with [^18^F]FDG after 4 weeks of treatment showing the same 2 lesions. **C)** [^18^F]FDG SUV before and after treatment of the lung lesions was variable and occasionally increased. **D)** Total probe uptake as determined by total lesion glycolysis (TLG) of the lesions showed a lesion-dependent pattern suggestive of a highly refractory response to [^18^F]FDG uptake, despite treatment. **E and F)** By contrast, similarly-timed (pretreatment and 4 weeks of treatment) **[^18^F]FDT** scans 90 minutes post injection (E 0.7 mCi; F 1 mCi) showed reduced SUV and total **[^18^F]FDT** uptake in the lesions after 4 weeks of treatment. Low bacterial load was indeed observed in the lesions: lesion 1 = 2.6 logCFU, lesion 2 = 2.8 logCFU. **G)** After HRZE combination drug therapy, the **[^18^F]FDT** SUV/CMR of the lung lesions was significantly decreased (p< 0.001) and **H)** the total **[^18^F]FDT** uptake in the lesions, as determined by TLG, was also reduced (p<0.005). One representative image pair is shown of three similar treatment experiments.

**Figure 6.**
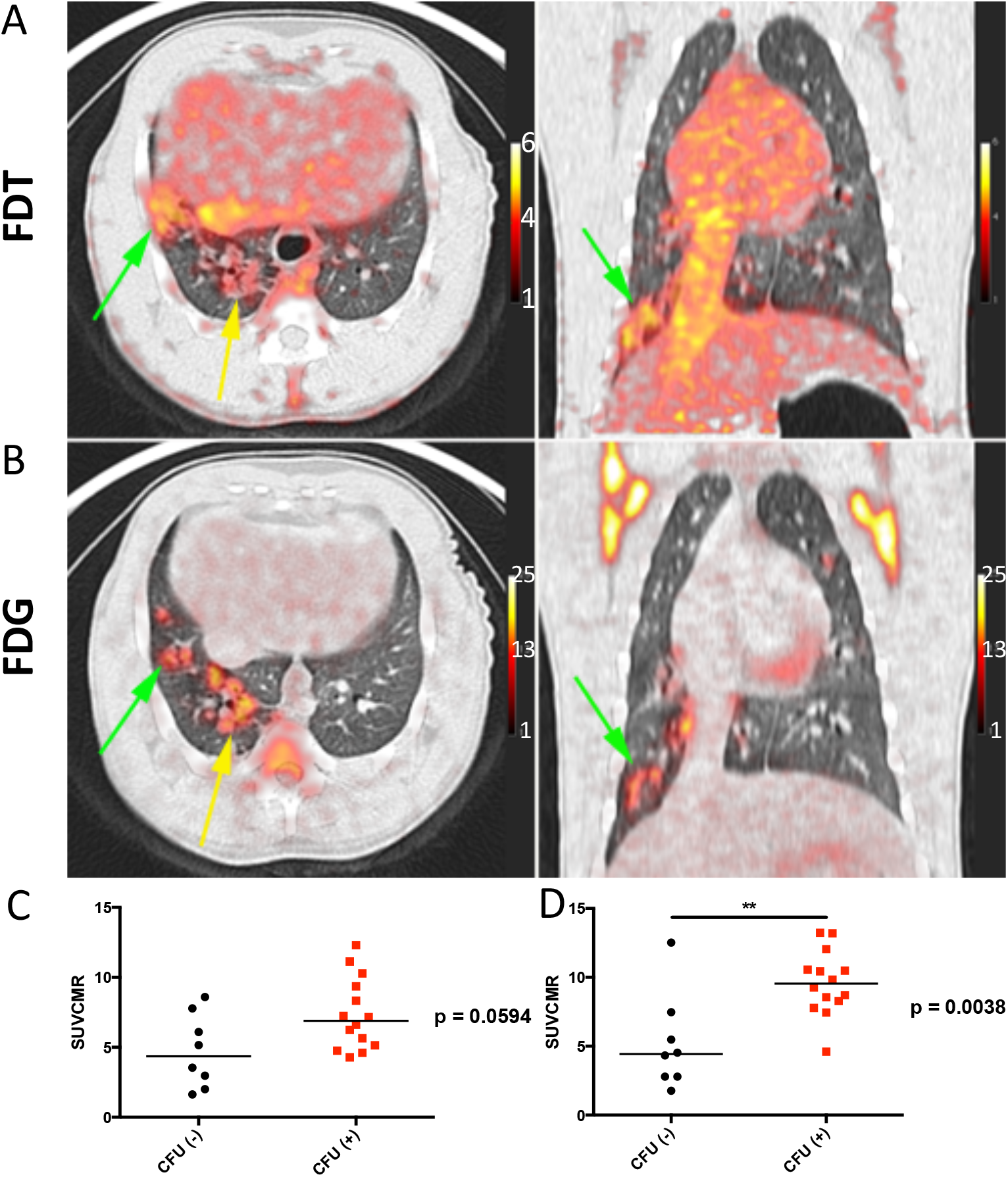
[^18^F]FDT is also an Effective and Specific Radiotracer in an ‘Old World’ NHP TB Model. **A)** Axial and coronal views of the lung of a representative Mtb-infected cynomolgus macaque showing labeling of the tubercular lesion clusters labeling (green and yellow arrows) with **[^18^F]FDT** (5 mCi) scanned 60 minutes post-injection. Calibration scale in SUV. **B**) Axial and coronal views of the same lung lesion clusters labeled with [^18^F]FDG (5 mCi) 60 minutes post-injection. Calibration scale in SUV. **C**) The animal was necropsied and the individual lesions were plated. *Ante mortum* probe uptake of the lesions was compared to their culture status. In images captured at 60 minutes there was a trend toward lesions with culturable bacteria having higher probe uptake. **D**) By 120 minutes, the uptake of probe was significantly higher among lesions with culturable mycobacteria than among sterile lesions. Data from one of three representative animals are shown.

### [^18^F]FDT is also an Effective Mtb-Radiotracer in an ‘Old World’ Non-Human Primate TB Model

Whilst, NHPs represent the most realistic pre-clinical TB models, species-specific differences are known.^54^ To confirm the results (see above) found in the ‘New World’ (marmoset) NHP models, we also tested the **[^18^F]FDT** in ‘Old World’ cynomolgus macaques (n = 3) infected with *Mtb* (strain Erdman) (**Figure 6**). TB in cynomolgus macaques is characterized by heterogeneous granulomas with a spectrum of *Mtb* burden, from lesions with many culturable bacilli to sterile lesions without live *Mtb*^55^. As in marmosets, **[^18^F]FDT** proved to be just as effective in specifically visualizing lesions (**Figure 6A**) in a manner that again is distinct from and complements [^18^F]FDG (**Figure 6B**). The uptake of **[^18^F]FDT** into cynomolgus macaques correlated well with bacterial disease burden, as lesions with culturable *Mtb* bacilli exhibited higher uptake of **[^18^F]FDT** (**Figure 6 C, D**).

### [^19^F]FDT is non-toxic and **[^18^F]FDT** shows rapid clearance resulting in low background dose exposure

No signs of toxicity or adverse effects were observed in any of the imaging studies described above. To confirm this apparent compatibility as well as determine the broader physiological characteristics of this novel radiotracer, both the biodistribution and toxicity of FDT were assessed in healthy animals.

First, dynamic PET imaging was performed in a healthy rhesus macaque for approximately 115 min after bolus intravenous administration of 151 MBq of **[^18^F]FDT** (**Figure 7A**). This indicated rapid distribution of **[^18^F]FDT** from the blood stream through the tissues and prominent accumulation of radioactivity in kidney and urinary bladder, indicating renal excretion with minimal accumulation in other organs. Organ radiation absorbed doses were calculated by extrapolating the rhesus macaque data to humans, with organs receiving the highest doses being the urinary bladder wall (0.143 mSv/mBq; 2 hr voiding interval), kidneys (0.119 mSv/MBq), and adrenals (0.022 mSv/mBq) (**Figure 7B**). The effective dose was 0.0154 mSv/MBq. Next, additionally, we performed biodistribution studies in two *Mtb*-infected marmosets injected with 74 MBq/kg **[^18^F]FDT**. Small samples of tissue were collected at necropsy 120 min post-injection and the radioactivity was measured in a calibrated gamma counter. In both animals, there was little tissue uptake of the radiotracer in organs other than in kidney (**Figure 7C**). Finally, single (acute) and multiple (chronic) intravenous (iv) dose toxicity studies of [^19^F]FDT, enabled by ready synthesis (see above), were conducted in both rats and beagle dogs. The animals were given either daily iv injections of [^19^F]FDT at 100 × the expected human dose for seven consecutive days or a single iv injection at either 100 × or 1000 × the expected human dose once. Thereafter animals were observed daily for mortality and morbidity, clinical observations, body weight, food consumption, clinical pathology (hematology, serum chemistry and coagulation parameters). All animals survived until scheduled euthanasia (day 9 or day 21). There were no adverse findings in any the parameters evaluated in the study.

**Figure 7.**
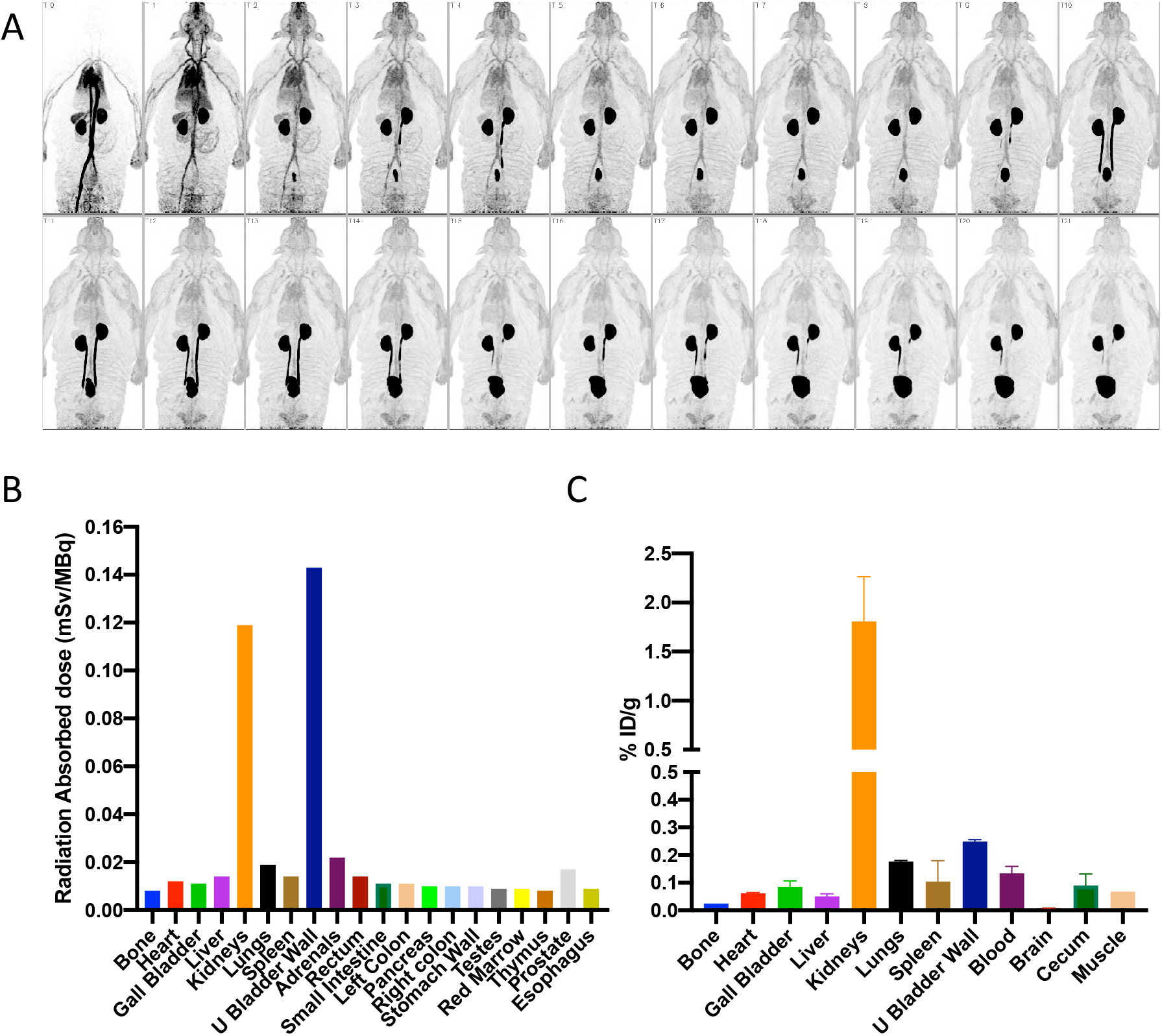
Dynamic Biodistribution of [^18^F]FDT in Naïve Rhesus Macaques and Infected Marmosets to Estimate Organ Exposure. **A)** Coronal maximum intensity projections of **[^18^F]FDT** PET activity in a representative naïve rhesus macaque collected over approximately 115 min in scan frames of increasing duration in a dynamic PET scan (from top left) after administration of 151 MBq of **[^18^F]FDT**. **B)** Organ radiation absorbed doses (mSv/MBq) calculated by extrapolating the rhesus macaque data to humans; organs receiving the highest doses were urinary bladder wall, kidneys, and adrenals. **C)** Similar to the rhesus experiments, the quantification of the radioactivity in marmoset tissues (using gamma counting of **[^18^F]FDT** radioactivity in excised tissues of euthanised marmosets) indicates the kidneys were the solid organs with the highest proportion of the injected dose (%ID/g) when marmosets were necropsied 120 min after being administered **[^18^F]FDT**.

## Discussion

Our results reveal that readily-generated **[^18^F]FDT** appears to complement FDG as an effective tracer of TB, with better selectivity and correlation to mycobacterial burden in lesions. We suggest that this selectivity is likely a result of its mechanism-based mode-of-action (**Figure 1A**) that unlike FDG co-opts specific TB-associated enzymes not present in host, allowing selective incorporation into mycobacterial lipids. Nonetheless, FDG is a known, effective probe of TB. Together these observations suggest that FDG may report on inflammation proximal to but not located directly in a bacterial lesion, whereas **[^18^F]FDT** appears to report directly on site of *Mtb* enzymatic action (and therefore on bacterial viability). The lack of direct co-location (see **Figure 4**) suggests that some tubercular lesions may therefore be seemingly less metabolically active than inflammatory hotspots. In this context, it may be that these differences are speculatively indicative of the physiology of host response, including the highly active balance of pro-and anti-inflammatory mediators,^56^ perhaps differentially driven by different cell subpopulations with different metabolic activities and locations within a lesion.

In the context of clinical evaluation of therapeutic response to TB treatment it is noteworthy that FDG avidity is maintained in a large fraction of cured patients even at the end of therapy.^57^ This suggests that inflammation persists even after bacterial viability is lost and so limits the use of FDG-PET in determining the appropriate duration of treatment. Moreover, bacterial viability in sputum is lost months before patients can be safely discontinued without experiencing disease relapse; this suggests clear limits on the use of sputum sampling. Our proposed use of **[^18^F]FDT**-PET now offers the important potential to have real-time monitoring of bacterial viability in the absence of culturable bacteria and may be an important tool in creating novel treatment regimens of shorter duration.

Our results also suggest that use of **[^18^F]FDT** can now avoid potentially refractory sources of signal from non-TB-associated inflammatory responses that potentiate metabolism in a lesion-like region. This has long been a consideration in patients with other diseases associated with inflammation of the lung, for example cancer^11-13,58^, but has been further highlighted by confusion generated from FDG-PET in putative COVID-19 patients with lung pathology.^14^ Moreover, other granulomatous disorders, where histopathological distinction from TB proves difficult, such as Crohn’s disease ^59^, might also now be more accurately diagnosed.

Together therefore our results suggest that disease-selectivity demonstrated by **[^18^F]FDT** will allow effective monitoring of disease and its treatment. This is likely to be invaluable for clinical evaluation of not only effectiveness in the development of new therapies but also in longitudinal monitoring and hence compliance.

The scaleable, pyrogen-free synthesis methods that we describe here use highly selective biocatalysis under aqueous conditions that are therefore readily implemented by the non-expert. This biocatalytic approach to utilization of FDG as a ready organic source of ^18^F now complements chemical methods, such as the elegant use of ^18^F-fluorodeoxysorbitol (FDS) that can be directly generated in a one-step reduction and has shown powerful utility in other bacterial species such as *E. coli*;^60^ the extension here to a ready, one-pot multi-step method now reveals the potential to consider even more complex sugar-based probes. This now enables potential, distributable radiochemical synthesis of **[^18^F]FDT** to be conducted anywhere there is access to FDG; scales of up to grams have now been demonstrated. Furthermore, full, pre-clinical assessments (see above) reveal no adverse effects. This, as well as the high specific activities and good radiochemical efficiencies that we disclose here now suggest **[^18^F]FDT** as a new, viable radiotracer for TB, suitable for Phase 1 trials.

## Materials and Methods

### Materials

Non-radioactive [^19^F]FDG was purchased from CarboSynth. For radioactive synthesis, [^18^F]FDG was purchased from Cardinal Health Ltd. Normal saline was obtained from Quality Biological (Gaithersburg, MD, USA). All other chemicals and solvents were received from Sigma-Aldrich (St. Louis, MO, USA) and used without further purification. Column and Sep-Pak cartridges used in this synthesis were obtained from Agilent Technologies (Santa Clara, CA, USA) and Waters (Milford, MA, USA), respectively. Sep-Paks were conditioned prior to use with 5 mL absolute ethanol. Analytical HPLC analyses for radiochemical work were performed on an Agilent 1200 Series instrument equipped with multi-wavelength detectors. Mass spectra (MS) of decayed [^18^F]-FDT solutions were recorded on a 6130 Quadrupole LC/MS, Agilent Technologies instrument equipped with a diode array detector. LC-MS analysis of [^18^F]-FDT was performed on Agilent 1260 HPLC system coupled to an Advion expression LCMS mass spectrometer with an ESI source. The LC inlet was Agilent 1200 series chromatographic system equipped with 1260 quaternary pump, 1260 Infinity autosampler, 1290 thermostatted column compartment and radiation detector. Hexokinase enzyme was purchased from Megazyme. Detailed procedures for expression and purification of other enzymes (OtsA also known as TPS, OtsB also known as TPP, OtsAB fusion and TreT variants) both in *E. coli* and Sf9 insect cells (the latter denoted by use of *^Sf^*) are given in the Supplementary Methods and below.

### Enzyme Kinetics

Kinetic parameters of TreT-catalysed synthesis of trehalose were assayed continuously by coupling the formation of UDP to the reactions of pyuruvate kinase and lactate dehydrogenase. The decrease in absorbance of NADH at 340 nm (ε340= 6220 M^-1^cm^-1^) was measured at 37 °C using a spectrophotometer equipped with a thermostat. The rate of UDP formation is proportional to the rate of NADH oxidation where one molecule of NADH is oxidized for each molecule of UDP formed. As the reaction involves two substrates, therefore, the kinetic parameters determined based on pseudo substrate concentration. TreT enzymes were incubated in the presence of fixed acceptor concentration and variable donor concentration, followed by fixed donor concentration and variable acceptor concentration. For continuous assay, each reaction was measured as triplicates by setting up the reactions in 96 well plates. Reactions were initiated by adding TreT enzyme and incubated at 37 °C (see supplementary methods for further details).

OtsA kinetic parameters were also determined by monitoring the formation of T6P by NADH-linked continuous assay. The rate of T6P formation is proportional to the rate of NADH oxidation where one molecule of NADH is oxidized for each molecule of T6P formed. OtsB-catalyzed trehalose formation was monitored by Malachite-green assay protocol as described previously ^61^.

For the NADH-linked based assays, the absorbance values obtained from the linear range of concentrations were plotted against time to determine the rate of the reaction (velocity). Plot of variable substrate concentrations against Rate, processed in Graphpad Prism to obtain Michaelis-Menton plots. From these plots kinetic parameters of each enzyme were calculated. All results shown are based on triplicate analysis of the samples.

### Enzyme production scale-up in Sf9 cells expression

In order to obtain a reasonable expression level of OtsA and OtsB proteins, the Bac-to-Bac® Baculovirus Expression System was selected. The original coding sequence of *E. coli* OtsA (*otsA*, CAA48913.1) and OtsB (*otsB*, CAA48912.1) was optimized according to *S. frugiperda* MNPV codon usage. The DNA fragment was then cloned into the *EcoRV* restriction enzyme site of the pUC57 vector. Then the plasmids pUC57-OtsA and pUC57-OtsB and pFastBac Dual vector were digested with *BamHI* and *NotI*, respectively, and ligated to create the recombinant plasmids of OtsA (pFastBac-OtsA) and OtsB (pFastBac-OtsB). The bacmid DNAs of OtsA (Bacmid-OtsA) and OtsB (Bacmid-OtsB) were generated by transformation of the recombinant plasmids pFastBac-OtsA and pFastBac-OtsB into the DH10Bac^TM^ *E. coli*, respectively. Correct insertions of the *otsA* and *otsB* genes into bacmid DNA were confirmed by PCR analysis using pUC/M13 standard primers: 5-CCCAGTCACGACGTTGTAAAACG-3 (forward) and 5-AGCGGATAACAATTTCACACAGG-3 (reverse).

Based on the bacmid DNAs, the recombinant baculovirus of OtsA and OtsB were made by transfection of Sf9 cells. Culture and preparation of Sf9 cells were performed according to the protocol provided by manufacturer. A Sf9 cell stock with a viability >95% and a density of 1.5 × 10^6^ – 2.5 × 10^6^ cells/mL was prepared before proceeding to transfection.

### Experimental procedures to generate P1 baculovirus for OtsA and OtsB

The P1 baculovirus stocks of OtsA and OtsB were generated using Sf9 cells in 6-well culture plates. Briefly, 2 ml of Sf9 cells were added to each well of two 6-well plates with the cell number of 8 × 10^5^ in each well. Cells were allowed to attach for 15 minutes at room temperature. Transfectin II was used as the transfection reagent and the transfection experiments were performed according to the protocol provided by manufacturer. Different amount of bacmid DNAs of OtsA and OtsB were used. After transfection, the Sf9 cells were further cultured at 27 °C for 3 days, then the P1 baculovirus (in supernatant) was harvested by centrifugation. The P1 baculovirus are found in the culture medium and the Sf9 cell pellets were used to check protein expression. Fetal bovine serum (FBS) was then added to each P1 baculovirus solution to reach a final concentration of 2%. The P1 baculovirus stock solution were then stored at 4 °C, protected from light.

Amplification of P1 baculovirus of OtsA and OtsB: In order to make baculovirus stock for large-scale protein expression and to increase virus titer, P2 and P3 baculovrius stocks of OtsA and OtsB were prepared from their P1 baculovirus. Briefly, a total of 150 mL of suspended Sf9 cells with a cell density of 2.0 × 10^6^ cells/mL was infected by P1 baculovirus of OtsA or OtsB, respectively. The 150 mL infection culture composition was: 1% P1 baculovirus of OtsA or OtsB, 1% FBS, 1% ethanol, Sf9 cells with a density of 2.0 × 10^6^ cells/mL. After infection at 27 °C with shaking (150 rpm) for 72 hours, the P2 baculovirus (in culture medium) was harvested by centrifugation at 1100 rpm for 10 minutes at room temperature. FBS was added to a final concentration of 2%, then the solution was filtered by syringe filter (0.45 μm) and stored at 4 °C, protected from light. Similar infection procedures were performed to prepare P3 baculovirus, except using P2 baculovirus stock solution for cell infection.

### Expression and Purification of C-terminally tagged, recombinant OtsA

Sf9 cells were grown in serum-free medium to a density of 2 × 10^6^ cells/mL and a cell viability of greater than 95%. The cell culture (1 L) was transfected with the viral stock, 2.8 x 10^9^ pfu/mL, at a multiplicity of infection (MOI) of unity. At 72 h post-infection, the cells were harvested by centrifugation. After freeze/thaw cycle, the pellet was resuspended in lysis buffer (100 mM HEPES, 200 mM NaCl, 10 mM imidazole, 10 mM MgCl_2_, 1 mM βME). DNase 3 mg and 1x Complete® protease inhibitor cocktail was also added into the protein pellet and lysed using a homogenizer. Cell debris was removed by centrifugation and the cell free extract passed through a 0.45 μm membrane filter. The filtrate was applied to a Ni–NTA column equilibrated in lysis buffer. After removal of the cell debris by centrifugation, the protein of interest was purified with a linear gradient of 10–300 mM imidazole in 100 mM HEPES, 200 mM NaCl, 10 mM MgCl_2_, 1 mM βME, pH 7.5, using Ni-NTA. Protein samples were exchanged into 100 mM Hepes, 150 mM NaCl, 10 mM MgCl_2_, pH 7.5, using a Hiload^TM^26/60 Superdex^TM^ 200 desalting column. The sample was passed through an Acrodisc Mustang E membrane (Pall #MSTG25E3) to reduce endotoxin to 1.9 EU/mL. After the addition of 10% glycerol, the aliquots containing the recombinant enzyme were flash frozen, and stored at −80 °C.

### Expression and purification of recombinant OtsB

Insect cells were grown in serum-free medium to a density of 2 × 10^6^ cells/mL and a cell viability of greater than 95%. The cell culture (1 L) was transfected with the viral stock, 2.0×10^9^ pfu/mL, at a multiplicity of infection (MOI) of unity. At 72 h post-infection, the cells were harvested by centrifugation. After freeze/thaw cycle, the pellet was resuspended in lysis buffer (50 mM Tris-HCl, 200 mM NaCl, 5 mM MgCl_2_, 30 mM imidazole, 1 mM βME). DNase 3 mg, 1x Complete® protease inhibitor cocktail was also added in the pellet and lysed using a homogenizer. Cell debris was removed by centrifugation and the cell free extract passed through a 0.45 µm membrane filter. The filtrate was applied to a Ni–NTA column equilibrated in lysis buffer. After removal of the cell debris by centrifugation, the protein of interest was purified with a linear gradient of 10–300 mM imidazole in 50 mM Tris-HCl, 200 mM NaCl, 5 mM MgCl_2_, 1 mM βME, pH 7.5, using Ni-NTA. The fractions with potential protein were concentrated using Vivaspin (5 MWCO) and then desalted on Hiload^TM^ 26/60 Superdex^TM^ 200 with 50 mM Tris-HCl, 5 mM MgCl_2_, 100 mM NaCl, pH 7.5. The sample was passed through an Acrodisc Mustang E membrane (Pall #MSTG25E3) to reduce endotoxin to 1.9 EU/mL. After the addition of 10% glycerol, the aliquots containing the recombinant enzyme were flash frozen, and stored at −80 °C. SDS–PAGE analysis revealed the presence of a protein with the correct molecular weight.

### FDT Synthesis, purification and characterization

In a 50 mL sterile falcon tube were added; ^19^F-2-FDG (100 mg, 367 mM), ATP (320 mg, 368 mM) and MgCl_2_ (2 mg). The contents were dissolved in HEPES (100 mM HEPES) buffer solution containing 100 mM NaCl (100 mM, 1 ml, pH 7.5). The pH was then adjusted to 8.0 with NaOH (5 M). Hexokinase (0.6 mL) was added to the reaction mixture to initiate the reaction. The reaction mixture was incubated at 30 °C, 50 rpm for 1-2 hours. Phosphorylation of FDG to 2-FDG6P was monitored by TLC and ^19^F NMR spectroscopy. Full conversion to 2-FDG6P was observed within 2 hours for each batch synthesized.

Within the same pot, UDP-glucose (200 mg, 110 mM), KCl (100 mM) and OtsA enzyme (26 μM) was added. The total reaction volume was then adjusted to 3 mL. The reaction mixture was incubated to similar conditions as above for 24 h to allow appropriate conversion to FDT6P. The conversion of FDG6P to FDT6P was monitored by ^19^F-NMR spectroscopy. The average maximum conversion after 24 h was between 65-70 %. After 24 h, in the same reaction mixture OtsB enzyme (6.23 μM) was added. The pH was adjusted to 8.0 with NaOH (5 M) and reaction mixture was incubated at 37 °C, 50 rpm. Reaction progress was monitored by TLC and ^19^F NMR. The reaction was usually complete within 2 hours.

The enzymes from crude reaction mixture were precipitated by either heating the mixture at 95 °C for 10 min or adding 2 mL EtOH and isolated by centrifugation (Viva spin, 10,000 MWCO) and/or filtration. The resulting filtrate was lyophilized and the crude mixture was further precipitated with MeOH:MeCN 90:10 solution and then purified by flash column chromatography. For purification normal phase silica was used as stationary phase and a mixture of water:isopropanol:ethyl acetate (4:48:48 v:v:v) was used as mobile phase. FDT was separated from residual FDG6P and residual FDG. The fractions containing FDT were pooled, concentrated and then lyophilised to dryness. The lyophilization step was repeated for a further 2-3 times to get rid of any residual water. The freeze-dried compound was then characterized by proton (^1^H NMR), carbon (^13^C NMR), fluorine (^19^F NMR) and ESI-MS.

### Manual Radiosynthesis and characterization of [^18^F]-FDT

For radiosynthesis using 3 enzymes, to a 1.5 or 2 mL microcentrifuge tube containing a solution of commercial [^18^F]-FDG in 0.9 % NaCl, reactants were added sequentially to achieve final concentration of UDP-glucose 30 mM, MgCl_2_ 10 mM, KCl 100 mM, ATP 10 mM, Hexokinase 640 U, OtsA 0.7 mg and OtsB 0.4 mg and enough HEPES buffer (100 mM HEPES, 150 mM NaCl and 10 mM MgCl_2_, pH 7.5) to achieve the desired volume of 1 mL. After gentle mixing, the tube was capped and incubated for 60 min at 37 °C in a thermoshaker inside the hot cell. Reaction progress was monitored by radio-HPLC during this period. After 40 min, reaction was almost 80 % completed. After 60 min, the enzymes in the reaction mixtures were precipitated by adding ethanol solution, filtered through 5 µm filter and then purified by Luna-NH_2_ Sep-Pak cartridge method (details in the supplementary methods).

Radiochemical yield of the purified product was calculated by assaying the activity (mCi) of the vial, reported as non-decay corrected yield and is based on starting activity of [^18^F]-FDG. The radiochemical purity of the product was analysed by radio-HPLC using an Agilent 1200 series HPLC coupled with radio detector for monitoring the radioactivity. Radiochemical purity was determined by calculating the area under the curves in the radiochromatogram.

For radiosynthesis using 1 enzyme i.e. TreT similar procedure as described above was adopted. TreT being thermophile allowed the incubation of the reaction at higher temperature, therefore we carried out the reactions both at 37 °C and 60 °C.

### Chemoenzymatic synthesis of [^18^F]-FDT for the animal models

For the biodistribution and blocking studies, [^18^F]-FDG (20-30 mCi in 0.8 −1 mL) was added to the reaction mixture containing 100 µL 1 M HEPES buffer, pH 7.6, 20 µL 1 M MgCl_2_, 20 µL 1 M ATP, 60 µL 1 M UDP-glucose, ∼50 µL OtsA (1 mg), ∼50 µL OtsB (1 mg), 20 µL hexokinase (5 mg). The reaction mixture was incubated at 37 °C for 30 min. After 30 min, the mixture was diluted with absolute ethanol (4 mL) and passed through a 5 µm syringe filter. The eluent was passed slowly through an amine Sep-Pak SPE cartridge at a flow rate of 1-2 drops per second. The eluent was then concentrated in vacuo. The resulting solution was filter-sterilized into a sterile vial for delivery. Identity of the compound was confirmed by LC-MS.

### Automated Synthesis of [^18^F]-FDT

[^18^F]-FDG (1480-3700 MBq; 40-100 mCi in 0.8 −1.8 mL) was transferred under vacuum to the Reactor 1 containing 100 µL 1 M HEPES buffer, pH 7.6, 20 µL 1 M MgCl_2_, 20 µL 1 M ATP, 60 µL 1 M UDP-glucose, ∼50 µL OtsA (1 mg), ∼50 µL OtsB (1 mg), 20 µL hexokinase (5 mg). The reaction mixture was incubated for 30 min at 45 °C and absolute ethanol was added (3 - 6 mL). The reaction mixture was passed through the filter (5 µm) and a stack of three NH_2_-cartridges and collected in Reactor 2. An additional 1 mL 75% ethanol in water was added to rinse the Reactor 1 and transferred into the Reactor 2. The combined solution was concentrated under nitrogen at 60 °C for 10 min. 2 mL saline was added to the Reactor 2. The final [^18^F]-FDT solution was transferred to the product vial through a sterile filter (0.22 µm,). The quality of the product was determined by analytical HPLC (Condition: 4.6 x 250 mm, 2.7 µm AdvanceBio Glycan Mapping Column; solvent A = 50 mM ammonium formate, pH 4.5, solvent B = acetonitrile; Flow rate = 0.5 mL/min;gradient 0-15 min 68-62% B; 15-20 min 62-68% B). Identity of the compound was confirmed by LC-MS (**Supplementary Figure S32**)

### Specific Activity Analysis of [^18^F]FDT

The radioactivity of the final products, [^18^F]-FDT, was obtained using the radiation detector. To quantify the mass of the decayed [^18^F]-FDT in the form of [^18^O]trehalose, a modified version of a trehalose quantification enzymatic assay^46^ was used. A trehalose standard calibration curve (linear fit, R^2^ = 0.9984) was generated, following the enzymatic assay using known concentrations of trehalose. The decayed masses of [^18^F]FDT from the syntheses were calculated based on this calibration curve. Then, the specific activity of the final product was calculated, following the definition, radioactivity at the end of the synthesis/unit mass of compound.

### Chemical Synthesis of 4-OTf Trehalose Ac 7

In a 10 mL tube, 50 mg of 4-OH trehalose Ac_7_ (0.078 mmol, 1 eq) and 24 mg of 2,6-di-t-butyl-4-methylpyridine (0.018 mmol, 1.5 eq) were dissolved in 3 mL of anhydrous dichloromethane. The mixture was cooled to 0 °C and 17 µL of triflic anhydride (0.012 mmol, 1.3 eq) was added slowly. After stirring at 0 °C for 1 h, the mixture was warmed to room temperature for 3 d. The reaction mixture was evaporated to dryness at low temperature and purified by silica column chromatography (9:1 CH_2_Cl_2_:EtOAc and then 7:3 of the same mixture, elution completed 15 min after loading) to give 25.9 mg of 4-OTf Tre Ac_7_ as a white solid in 43% yield. TLC (1:1 CH_2_Cl_2_:EtOAc) Rf=0.58 for the product and Rf=0.40 for starting material 4-OH Tre Ac_7_. TLC: Rf=0.58 (1:1 CH_2_Cl_2_:EtOAc, starting material 4-OH Tre Ac7 Rf=0.40);^1^H NMR (400 MHz CDCl_3_) δ ppm 2.04, 2.07, 2.08, 2.08, 2.11, 2.13 (7x 3H, s, CH_3_ OAc), 4.02 (1 H, dd, J6b’,6a’ = 12.0 Hz, J6b’,5’ 2.4 Hz, H-6b’ CH_2_OAc), 4.06 (1H, ddd, J5’,4’ = 10.4 Hz, J5’,6a’ 5.6 Hz, J5’,6b’ = 2.4 Hz, H-5’), 4.22 (3 H, m, H-5, H-6b, H-6a), 4.99 (1H, dd, J4,5 10.0Hz, J4,3 9.4Hz, H-4) 5.03 (1 H, dd, J2,3 = 10.4 Hz, J2,1 4.0 Hz, H-2), 5.06 (1H, dd, J2’,3’ = 10.4 Hz, J2’,1’ 4.0 Hz, H-2’), 5.07 (1 H, dd, J4’,5’ = 10.4 Hz, J4’,3’ 9.6 Hz, H-4’), 5.27 (1H, d, J1,2 4.0 Hz, H-1), 5.29 (1H, d, J1’,2’ 4.0 Hz, H-1’), 5.48 (1H, dd, J3’,2’ = 10.4 Hz, J3’,4’ 9.6 Hz, H-3’), 5.70 (1H, dd, J3,2 = 10.4 Hz, J3,2 9.4 Hz, H-3); ^13^C NMR (101 MHz CDCl_3_) δ ppm 20.4, 20.5, 20.6, 20.6, 20.6, (7x CH_3_ acetates), 61.0 (C-6), 61.6 (C-6’), 67.2 (C-5), 68.3, 68.3 (C-5’), 68.4, 68.4 (C-3, C-4’), 69.8 (C-2 and C-2’), 70.0 (C-3’), 78.7 (C-4 OTf), 92.3 (C-1), 92.8 (C-1’), 169.3, 169.5, 169.8, 170.1 (7x C=O acetates); ^19^F NMR (376.5MHz, CD_3_OD):-74.4, s, CF3 OTf; HRMS ESI^+^ [C_27_H_35_O_20_F_3_S+Na]^+^ requires 791.1287, found 791.1246.

### Chemical Synthesis of 6-OTf Trehalose Ac 7

In a 50 mL flask, 160.6 mg of 6-OH trehalose Ac_7_ (0.252 mmol, 1 eq) and 77 mg of 2,6-di-t-butyl-4-methylpyridine (0.378 mmol, 1.5 eq) were dissolved in 10 mL of anhydrous dichloromethane. The mixture was cooled to 0°C and 55 µL of triflic anhydride (0.328 mmol, 1.3 eq) was added slowly. The mixture was stirred while monitoring by TLC, 1 h at 0°C followed by 24 h at r.t. The reaction mixture was evaporated to dryness at low temperature and purified by silica column chromatography (9:1 CH_2_Cl_2_:EtOAc and then 7:3 of the same mixture, elution completed 15 min after loading) to give 93.3 mg of 6-OTf Tre Ac_7_ as a white solid in 48% yield. TLC (1:1 CH_2_Cl_2_:EtOAc) Rf=0.58 for the product and Rf=0.40 for starting material 6-OH Tre Ac_7_. ^1^H NMR (400 MHz CDCl_3_-d) δ ppm 2.03, 2.03, 2.04, 2.05, 2.08, 2.09, 2.09 (7x 3H, s, CH_3_ OAc), 3.99 (1 H, dd, J6b’,6a’ = 12.0 Hz, J6b’,5’ 2.4 Hz, H-6b’ CH_2_OAc), 4.05 (1H, ddd, J5’,4’ = 10 Hz, J5’,6a’ 5.6 Hz, J5’,6b’ = 2.4 Hz, H-5’), 4.22 (1H, m, H-5) 4.25 (1 H, m, H-6a’ CH_2_OAc), 4.41 (1 H, dd, J6b,6a = 11.2 Hz, J6b,5 2.4 Hz, H-6b CH_2_OTf), 4.52 (1 H, dd, J6a,6b = 11.2 Hz, J6a,5 6 Hz, H-6a CH_2_OTf), 5.00-5.10 (4H, m, H-2, H-2’, H-4, H-4’), 5.28 (1H, d, J1’,2’ 3.6 Hz, H-1’), 5.33 (1H, d, J1,2 3.6 Hz, H-1), 5.48 (1H, dd, J3,4 = 10.0 Hz, J3,2 9.6 Hz, H-3), 5.50 (1H, dd, J3’,4’ = 10.0 Hz, J3’,2’ 9.6 Hz, H-3’); ^13^C NMR (101 MHz CDCl_3_-d) δ ppm 20.6, 20.7, 20.7 (7x CH_3_ acetates), 61.7 (C-6’ OAc), 67.8 (C-5), 68.3 (C-5’), 68.3 (C-4), 68.4 (C-4’), 69.4 (C-2, C2’), 69.5 (C-3’), 70.0 (C-3), 73.3 (C-6 OTf), 92.4 (C-1), 93.1 (C-1’), 169.3, 169.4, 169.5, 169.6, 169.9, 170.0, 170.6 (7x C=O acetates); ^19^F NMR (376.5MHz, CD_3_OD):-74.4, s, CF_3_ OTf; HRMS (ESI^+^) calcd. for C_27_H_35_F_3_O_20_SNa^+^ (M+Na^+^): 791.1287, found 791.1259

### Animal and Ethics Assurance

This study was carried out in accordance with the recommendations in the Guide for the Care and Use of Laboratory Animals of the National Institutes of Health. Biodistribution studies with naive rhesus macaques were approved by the Institutional Animal Care and Use Committee (IACUC) of the NIH Clinical Centre (Bethesda, MD). The IACUC of the NIAID, NIH approved the experiments described herein with rabbits and marmosets under protocols LCID-3 and LCID-9 respectively (Permit issued to NIH Intramural Research Program as A-4149-01), and all efforts were made to provide intellectual and physical enrichment and minimize suffering. Once infected, female rabbits or marmosets of both sexes were housed individually or paired in biocontainment cages in a biological level 3 animal facility approved for the containment of *M. tuberculosis*. All NIH studies were performed in accordance with the regulations of the Division of Radiation Safety, at the National Institutes of Health (Bethesda, MD, USA). The University of Pittsburgh cynomolgus macaque studies were approved by its IACUC and Division of Radiation Safety and animals were pair housed in an approved ABSL3 facility (Regional Biocontainment Facility, Pittsburgh PA).

### Detection of serum [^19^F]FDT

FDT standards were prepared in human plasma (Sigma Aldrich), whereby 20 μL plasma samples were spiked with various known concentrations of [^19^F]FDT i.e. 10 μM, 20 μM, 40 μM, 120 μM, 160 μM, and 300 μM in 1.5 ml Eppendorf tube and diluted with up to 50% acetonitrile solution. Samples were vortexed for 1 min and centrifuged at 13000 rpm for 10 min and supernatant was analysed directly by LC-MS using LC method 2.

For *in vivo* study, six white mice, CD1, male, weighing between 22-26 grams, were used. Two concentrations of [^19^F]FDT i.e. 500 μM and 50 μM (final concentration in the blood) were injected into the mice, through the tail vein injection. Prior to injection, each mouse was pre-bled (100 μL) and control samples were obtained. Five min post injection, blood samples were drawn up (775 μL) in syringe containing 75 μL sodium citrate as anticoagulant, hence total blood volume was 700 μL.

Samples were spun at 13000 rpm for 10 min and the supernatant (plasma) was used for further analysis. 200 μL of plasma sample was transferred into fresh 1.5 mL Eppendorf tube and diluted with equal amount of acetonitrile to precipitate out blood plasma protein and other macromolecules. Samples were further spun at 13000 rpm for 5 min. Supernatant was transferred into a mass spectrometry vial and analyzed using LC-MS analysis method 2 & 3 as shown in Supplementary Table S9.

### PET/CT instrumentation, probe administration, and image acquisition

Rabbits and marmosets at NIAID were anaesthetized and maintained during imaging as previously described^62,63^. Briefly, syringes of FDG or FDT were measured in a dose calibrator immediately before and after injection, targeting = injected doses of 2 mCi/kg. During uptake and distribution of the probes, a CT scan from the base of the skull spanning the lungs and the upper abdominal cavity was acquired as described for each species on a helical eight-slice Neurological Ceretom CT scanner (NeuroLogicia, Danvers, MA) operated as part of a hybrid pre-clinical PET/CT system utilizing a common bed. The animal bed was then retracted into the microPET gantry (MicroPET Focus 220, Siemens Preclinical Solutions, Knoxville, TN) and sixty minutes ± 5 min post FDG injection, a series of 2 or 3, 10 minute emission scans with 75 mm thick windows with a 25 mm overlap were acquired caudal to cranial. The FDT scans were acquired beginning at 60, 90, and 120 minutes ± 5 min post injection with the same duration, window and overlap. For the blocking studies, injections of 150 µg of [^19^F]-FDT synthesized as described above were administered 60 min and 5 min prior to the FDT tracer and the scans were acquired as before. Two animals were used in the blocking studies with one animal receiving the blocking agent while the other was administered saline only prior to the FDT tracer. Two days later, the administrations were reversed so that the second animal received the blocking agent. The emission data for all scans were processed and corrected as described previously^62^. For the blocking studies, injections of 150 µg of [^19^F]-FDT synthesized as described above were administered 60 min and 5 min prior to the FDT tracer injection and the scans were acquired as before. Two animals were used in the blocking studies with one animal receiving the blocking agent while the other was administered saline only prior to the FDT tracer. Two days later the administrations were reversed so that the second animal received the blocking agent. The emission data for all scans were processed and corrected as described previously^62^. Cynomolgus macaques at the University of Pittsburgh were imaged using a Siemens microPET Focus 220 PET scanner and a Neurological Ceretom CT scanner as previously described ^52^. Scans were viewed using OsiriX (Pixmeo, Geneva, Switzerland) to identify and analyze individual granulomas as done previously ^64^.

### Uptake / Time course analysis PET/CT data analysis

For FDG glycolytic activity measurements, PET/CT images were loaded into MIM fusion software (MIM Software Inc, Cleveland, OH USA) to create lung contours using the CT 3D region growing application with upper and lower voxel threshold settings of 2 and -1024 HU respectively with hole filling and smoothing applied. Dense lesion centers were subsequently identified for inclusion in the lung region manually and the program calculated the FDG signal parameters. In addition, each lesion within the lung was marked with a 3-D region of interest (ROI) and the SUV statistics for the ROIs were captured into excel sheets for analysis as previously described.^62^

### Biodistribution in Rhesus and Marmosets

An adult male rhesus macaque (12.4 kg) was administered 151 MBq of [^18^F]FDT via saphenous vein catheter after being anesthetized with ketamine and then maintained with 2-4% isoflurane/97% oxygen. Dynamic whole-body PET images were obtained on a Siemens mCT PET/CT scanner. The images were reconstructed using an iterative time-of-flight algorithm with a reconstructed transaxial resolution of about 4.5 mm. The PET acquisition was divided into 22 time frames of increasing durations, 2 x 15 sec, 4 x 30 sec, 8 x 60 sec, and 8 x 120 sec. Each frame in the sequence was gathered from 4 bed positions to obtain whole body dynamic data over the scan duration (see Fig 7A). Volumes of interest (VOIs) were drawn over major organs, non-decay corrected time-activity curves were generated and integrated, and then used to generate organ residence times that were scaled to human body and organ weights. Organ radiation absorbed doses and the effective dose were then calculated using OLINDA 2.1 (Vanderbilt University, Nashville TN) ^65,66^.

Two marmosets were administered 2 mCi/kg FDT (in ∼300 µL PBS) via tail vein catheter while anesthetized with 3% isoflurane/97% oxygen and PET/CT scans were collected as described above and a final 200 μL whole blood sample was drawn via femoral vein puncture and placed immediately into a 1.5 mL tube that was coated with potassium ethylenediamine tetraacetic acid. After euthanasia, the liver, right kidney, skeletal muscle (right quadriceps), brain, and bone (femur) and other organ samples were harvested and weighed. Each of the tissue samples were transferred to gamma counter tubes and counted on a Wallac Wizard 1480 Gamma Counter (Perkin Elmer, Waltham, MA). Count data were converted to percent injected dose per gram using a dose calibration curve.

### Metabolism analysis and Detection of labeled mono and dimycolates

Blood (100 µL) and urine (100-300 µL) were collected from animals and diluted 2:1 v/v with acetonitrile to sterilize them. These samples were then vortexed, centrifuged and the supernatant analyzed by either HPLC or TLC. TLC plates were visualized via exposure to a phosphor screen for 16 h prior to imaging using a phosphor screen and Typhoon FLA7000 (GE Healthcare, Pittsburgh PA). Silica gel plates used to assay the metabolism samples and lesion extracts were 150 Å Silica Gel HL 250µ m 10 x 20 cm channeled plates with a pre-adsorbent zone purchased from Analtech/iChromatography (Newark, DE). Plates were read using a BioScan AR-2000 reader.

### Toxicity studies

For toxicity studies, 2 g of [^19^F]-FDT was synthesized and tested for quality control as described above and in Supplementary Figure S37 and shipped to a contract research organization. Toxicity studies were conducted by SRI International, Biosciences Division, Menlo Park, CA, USA in rats and dogs.

For studies in rats, male and female Sprague Dawley rats (15/sex/group) were given a single iv administration of [^19^F]-FDT at 1.32 mg/kg (100 times of human dose; Group 3) or 13.2 mg/kg (1000 times of human dose; Group 4) on day 7 or 1.32 mg/kg/day for 7 consecutive days (total dose 9.24 mg/kg; Group 2. A control group (15/sex; Group 1) was given 7 days of vehicle (50 mM HEPES buffer, pH 7.5 with 7.5 mM MgCl_2_) at an equivalent dose volume of 1 mL/kg on days 1-7. Animals were euthanised on day 9 (main groups) or day 21 (recovery groups).

**Experimental Design for the toxicity Studies in Rats**
| **Group** | **Treatment** | **Dose Level**<br>**(mg/kg/day)** | **No.**<br>**Doses** | **Total**<br>**Dose**<br>**(mg/kg)** | **Main**<br>**groups**<br>**animal** | **Recovery**<br>**Group**<br>**animals** |
| --- | --- | --- | --- | --- | --- | --- |
| 1 | Vehicle | 0 | 7 | 0 | 10M/10F | 5M/10F |
| 2 | 100x human dose | 1.32 | 7 | 9.24 | 10M/10F | 5M/10F |
| 3 | 100x human dose | 1.32 | 1 | 1.32 | 10M/10F | 5M/10F |
| 4 | 1000x human dose | 1.32 | 1 | 1.32 | 10M/10F | 5M/10F |

To assess the toxicity associated with [^19^F]-FDT in dogs, male and female Beagle dogs (5/sex) were given a daily iv injection of [^19^F]-FDT at 0.4 mg/kg/day (100 times of human dose; Group 2) for 7 consecutive days, or a single iv administration of [^19^F]-FDT at 0.4 or 4 mg/kg (100 and 1000 times of human dose in Groups 3 and 4, respectively) on Day 7. A control group (5/sex; Group 1), was given 7 days of vehicle (HEPES buffer, pH 7.5, 50 mM with MgCl_2_ 7.5 mM), at an equivalent dose volume of 0.25 ml/kg on Days 1-7. Animals were sacrificed on Day 9 (3/sex/ group; main groups) or Day 21 (2/sex/group; recovery groups). The single dose administration for Groups 3-4 were initiated on Day 7 so that clinical pathology and necropsy occurred on the same calendar day for all Groups (1-4), thus allowing for sharing of vehicle control data for analysis of clinical pathology and necropsy results between the two dose regimens.

**Experimental Design for the toxicity Studies in Dogs**
| **Group** | **Treatment** | **Dose Level**<br>**(mg/kg/day)** | **No.**<br>**Doses** | **Total**<br>**Dose**<br>**(mg/kg)** | **Main**<br>**groups**<br>**animal** | **Recovery**<br>**Group**<br>**animals** |
| --- | --- | --- | --- | --- | --- | --- |
| 1 | Vehicle | 0 | 7 | 0 | 3M/3F | 2M/2F |
| 2 | 100x human dose | 0.4 | 7 | 2.8 | 3M/3F | 2M/2F |
| 3 | 100x human dose | 0.4 | 1 | 0.4 | 3M/3F | 2M/2F |
| 4 | 1000x human dose | 4 | 1 | 4 | 3M/3F | 2M/2F |

## Acknowledgments

We thank Becky Sloan for technical assistance with animal procedures and the Comparative Medicine Branch (NIAID) for animal handling. We thank the NIH PET Department for providing excellent technical support. We thank James Collier from the Nasmyth group and Shubhashish Mukhopadhyay from the Burgess-Brown group for their assistance in larger-scale enzyme expressions in insect cells. We also thank Oxford Chemistry Department NMR and Mass Spectrometry team for their continued support in experimentation and data analysis. The Chemistry theme at the Rosalind Franklin Institute is supported by the EPSRC (V011359/1 (P)).

## Author contributions

D.M.W., M.L.S., D.M.S., E.D., P.H., L.E.V., J.L.F., C.A.S., J.A.T., L.J.F. and C.E.B. designed and implemented the various animal studies; N.K., Y.M.A., G.A.M., F.B., R.S., F.D’H., D.O.K., B.G.D., and N.S.M. designed and performed the radiochemistry; N.K., F.D’H., S.S-L., and B.G.D designed and implemented the various cold chemistry experiments; N.K. and N.Y. established the insect cell expression system via virus bacmid DNA construction, virus amplification and initial expression and purification; N.K., S.S.L., Y.G. and R.R. expressed enzymes and performed kinetic analyses; K.M.B. performed initial syntheses and proof-of-principle experiments in prototype probes; W.D., M.K.P., F.G. and A.G.W. analyzed PET/CT scans; N.K., L.E.V., B.G.D., and C.E.B. drafted the manuscript; all authors approved the final version of the manuscript.

## Supplementary Information

Supplementary Materials and Methods Supplementary Figures (1-37)

Supplementary Tables (1-9)

## References

1. WorldHealthOrganization. Global Tuberculosis Report 2021. (Geneva (Switzerland), 2021).

2 Maitra, A. et al. Early diagnosis and effective treatment regimens are the keys to tackle antimicrobial resistance in tuberculosis (TB): A report from Euroscicon’s international TB Summit 2016. Virulence 8, 1005–1024 (2017). https://doi.org:10.1080/21505594.2016.1256536

3 Bhoil, A. et al. Challenges using PET-CT for international multicentre coordinated research projects in developing countries. Nuclear Medicine Communications 40, 93–95 (2019). https://doi.org:10.1097/MNM.0000000000000958

4 Gallach, M. et al. Addressing Global Inequities in Positron Emission Tomography-Computed Tomography (PET-CT) for Cancer Management: A Statistical Model to Guide Strategic Planning. Med Sci Monit 26, e926544–e926544 (2020). https://doi.org:10.12659/MSM.926544

5. TheInternationalAtomicEnergyAuthority. IAEA Medical imAGIng and Nuclear mEdicine (IMAGINE) <https://humanhealth.iaea.org/HHW/DBStatistics/IMAGINE.html. > (

6 Gambhir, S. S. Molecular imaging of cancer with positron emission tomography. Nat Rev Cancer 2, 683–693 (2002). https://doi.org:10.1038/nrc882

7 Langer, A. A systematic review of PET and PET/CT in oncology: a way to personalize cancer treatment in a cost-effective manner? BMC Health Serv Res 10, 283 (2010). https://doi.org:10.1186/1472-6963-10-283

8 Kim, I. J. et al. Double-phase 18F-FDG PET-CT for determination of pulmonary tuberculoma activity. Eur J Nucl Med Mol Imaging 35, 808–814 (2008). https://doi.org:10.1007/s00259-007-0585-0

9 Davis, S. L. et al. Noninvasive pulmonary [18F]-2-fluoro-deoxy-D-glucose positron emission tomography correlates with bactericidal activity of tuberculosis drug treatment. Antimicrobial Agents and Chemotherapy 53, 4879–4884 (2009). https://doi.org:10.1128/aac.00789-09

10 Yu, W. Y. et al. Updates on (18)F-FDG-PET/CT as a clinical tool for tuberculosis evaluation and therapeutic monitoring. Quant Imaging Med Surg 9, 1132–1146 (2019). https://doi.org:10.21037/qims.2019.05.24

11 Glaudemans, A. W. & Signore, A. FDG-PET/CT in infections: the imaging method of choice? Eur J Nucl Med Mol Imaging 37, 1986–1991 (2010). https://doi.org:10.1007/s00259-010-1587-x

12 Sathekge, M. et al. FDG uptake in lymph-nodes of HIV+ and tuberculosis patients: implications for cancer staging. Q J Nucl Med Mol Imaging 54, 698–703 (2010).

13 Corstens, F. H. & van der Meer, J. W. Nuclear medicine’s role in infection and inflammation. The Lancet 354, 765–770 (1999). https://doi.org:10.1016/s0140-6736(99)06070-5

14 Treglia, G. The role of 18F-FDG PET for COVID-19 infection: myth versus reality. Clinical and Translational Imaging 8, 125–126 (2020). https://doi.org:10.1007/s40336-020-00367-z

15 Weinstein, E. A. et al. Noninvasive determination of 2-[18F]-fluoroisonicotinic acid hydrazide pharmacokinetics by positron emission tomography in Mycobacterium tuberculosis-infected mice. Antimicrob Agents Chemother 56, 6284–6290 (2012). https://doi.org:10.1128/aac.01644-12

16 Xu, B. et al. Can multimodality imaging using 18F-FDG/18F-FLT PET/CT benefit the diagnosis and management of patients with pulmonary lesions? Eur J Nucl Med Mol Imaging 38, 285–292 (2011). https://doi.org:10.1007/s00259-010-1625-8

17 Elbein, A. D., Pan, Y. T., Pastuszak, I. & Carroll, D. New insights on trehalose: a multifunctional molecule. Glycobiology 13, 17r–27r (2003). https://doi.org:10.1093/glycob/cwg047

18 Woodruff, P. J. et al. Trehalose is required for growth of Mycobacterium smegmatis. Journal of Biological Chemistry 279, 28835–28843 (2004). https://doi.org:10.1074/jbc.M313103200

19 Bloch, H. Virulence of mycobacteria. Bibliotheca Tuberculosea 9, 49–61 (1955).

20 Kilburn, J. O., Takayama, K. & Armstrong, E. L. Synthesis of trehalose dimycolate (cord factor) by a cell-free system of Mycobacteriumsmegmatis. Biochemical and Biophysical Research Communications 108, 132–139 (1982). https://doi.org:10.1016/0006-291X(82)91841-1

21 Backus, K. M. et al. Uptake of unnatural trehalose analogs as a reporter for Mycobacterium tuberculosis. Nature Chemical Biology 7, 228–235 (2011). https://doi.org:10.1038/nchembio.539

22 Hodges, H. L., Brown, R. A., Crooks, J. A., Weibel, D. B. & Kiessling, L. L. Imaging mycobacterial growth and division with a fluorogenic probe. Proceedings of the National Academy of Sciences 115, 5271 (2018). https://doi.org:10.1073/pnas.1720996115

23 Holmes, N. J. et al. A FRET-Based Fluorogenic Trehalose Dimycolate Analogue for Probing Mycomembrane-Remodeling Enzymes of Mycobacteria. ACS Omega 4, 4348–4359 (2019). https://doi.org:10.1021/acsomega.9b00130

24 Kamariza, M. et al. Rapid detection of Mycobacterium tuberculosis in sputum with a solvatochromic trehalose probe. Science Translational Medicine 10 (2018). https://doi.org:10.1126/scitranslmed.aam6310

25 Dutta, A. K. et al. Trehalose Conjugation Enhances Toxicity of Photosensitizers against Mycobacteria. ACS Central Science 5, 644–650 (2019). https://doi.org:10.1021/acscentsci.8b00962

26. Initial aspects and concepts of this work were first disclosed at the International Carbohydrate Symposium, Tokyo, 2010.

27. The development of an optimized, pyrogen-free method was first disclosed in the following thesis.

28 Khan, R. Development of tuberculosis imaging probes based on sugars and proteins, University of Oxford, (2020).

29 Ishihara, R. et al. Molecular cloning, sequencing and expression of cDNA encoding human trehalase1The sequence data reported in this paper have been submitted to the DDBJ, EMBL and GenBank Sequence Databases with Accession Number AB000824.1. Gene 202, 69-74 (1997). https://doi.org:10.1016/S0378-1119(97)00455-1

30 Withers, S. G., Rupitz, K. & Street, I. P. 2-Deoxy-2-fluoro-D-glycosyl fluorides. A new class of specific mechanism-based glycosidase inhibitors. Journal of Biological Chemistry 263, 7929–7932 (1988).

31 Petroni, D., Menichetti, L. & Poli, M. Historical and radiopharmaceutical relevance of [18F]FDG. Journal of Radioanalytical and Nuclear Chemistry 323, 1017–1031 (2020). https://doi.org:10.1007/s10967-020-07013-y

32 Tewson, T. J., Welch, M. J. & Raichle, M. E. [18F]-labeled 3-deoxy-3-fluoro-D-glucose: synthesis and preliminary biodistribution data. Journal of Nuclear Medicine 19, 1339–1345 (1978).

33 Jiao, H. et al. Trehalase Gene as a Molecular Signature of Dietary Diversification in Mammals. Molecular Biology and Evolution 36, 2171–2183 (2019). https://doi.org:10.1093/molbev/msz127

34 Nigudkar, S. S. & Demchenko, A. V. Stereocontrolled 1,2-cis glycosylation as the driving force of progress in synthetic carbohydrate chemistry. Chemical Science 6, 2687–2704 (2015). https://doi.org:10.1039/C5SC00280J

35 Qu, Q., Lee, S.-J. & Boos, W. TreT, a Novel Trehalose Glycosyltransferring Synthase of the Hyperthermophilic Archaeon Thermococcus litoralis. Journal of Biological Chemistry 279, 47890–47897 (2004). https://doi.org:10.1074/jbc.M404955200

36 Kouril, T., Zaparty, M., Marrero, J., Brinkmann, H. & Siebers, B. A novel trehalose synthesizing pathway in the hyperthermophilic Crenarchaeon Thermoproteus tenax: the unidirectional TreT pathway. Archives of Microbiology 190, 355 (2008). https://doi.org:10.1007/s00203-008-0377-3

37 Urbanek, B. L. et al. Chemoenzymatic synthesis of trehalose analogues: rapid access to chemical probes for investigating mycobacteria. Chembiochem 15, 2066–2070 (2014). https://doi.org:10.1002/cbic.201402288

38 Ryu, S. I., Park, C. S., Cha, J., Woo, E. J. & Lee, S. B. A novel trehalose-synthesizing glycosyltransferase from Pyrococcus horikoshii: molecular cloning and characterization. Biochem Biophys Res Commun 329, 429–436 (2005). https://doi.org:10.1016/j.bbrc.2005.01.149

39 Giaever, H. M., Styrvold, O. B., Kaasen, I. & Strom, A. R. Biochemical and genetic characterization of osmoregulatory trehalose synthesis in Escherichia coli. Journal of Bacteriology 170, 2841–2849 (1988). https://doi.org:10.1128/jb.170.6.2841-2849.1988

40 Gosselin, S., Alhussaini, M., Streiff, M. B., Takabayashi, K. & Palcic, M. M. A continuous spectrophotometric assay for glycosyltransferases. Analytical Biochemistry 220, 92–97 (1994). https://doi.org:10.1006/abio.1994.1303

41 Errey, J. C. et al. Mechanistic insight into enzymatic glycosyl transfer with retention of configuration through analysis of glycomimetic inhibitors. Angew Chem Int Ed Engl 49, 1234–1237 (2010). https://doi.org:10.1002/anie.200905096

42 Wakelin, S. J. et al. “Dirty little secrets”—Endotoxin contamination of recombinant proteins. Immunology Letters 106, 1–7 (2006). https://doi.org:10.1016/j.imlet.2006.04.007

43 Iwanaga, S. Biochemical principle of Limulus test for detecting bacterial endotoxins. Proc Jpn Acad Ser B Phys Biol Sci 83, 110–119 (2007). https://doi.org:10.2183/pjab.83.110

44 Schwarz, H., Schmittner, M., Duschl, A. & Horejs-Hoeck, J. Residual Endotoxin Contaminations in Recombinant Proteins Are Sufficient to Activate Human CD1c+ Dendritic Cells. PloS one 9, e113840 (2014). https://doi.org:10.1371/journal.pone.0113840

45 Luckow, V. A., Lee, S. C., Barry, G. F. & Olins, P. O. Efficient generation of infectious recombinant baculoviruses by site-specific transposon-mediated insertion of foreign genes into a baculovirus genome propagated in Escherichia coli. Journal of Virology 67, 4566 (1993).

46 Carillo, P. et al. A fluorometric assay for trehalose in the picomole range. Plant Methods 9, 21 (2013). https://doi.org:10.1186/1746-4811-9-21

47 Via, L. E. et al. Differential Virulence and Disease Progression following Mycobacterium tuberculosis Complex Infection of the Common Marmoset (Callithrix jacchus). Infect Immun 81, 2909–2919 (2013).

48 Dheda, K., Barry, C. E., 3rd & Maartens, G. Tuberculosis. The Lancet 387, 1211–1226 (2016). https://doi.org:10.1016/S0140-6736(15)00151-8

49 Vijay, S. et al. Risk factors associated with default among new smear positive TB patients treated under DOTS in India. PloS one 5, e10043–e10043 (2010). https://doi.org:10.1371/journal.pone.0010043

50 Gleeson, L. et al. Recurrent TB in Ireland is predominantly due to relapsed infection and is frequently associated with poor compliance with therapy. European Respiratory Journal 48, PA2668 (2016). https://doi.org:10.1183/13993003.congress-2016.PA2668

51 Matthews, P. M., Rabiner, E. A., Passchier, J. & Gunn, R. N. Positron emission tomography molecular imaging for drug development. British Journal of Clinical Pharmacology 73, 175–186 (2012). https://doi.org:10.1111/j.1365-2125.2011.04085.x

52 Coleman, M. T. et al. PET/CT imaging reveals a therapeutic response to oxazolidinones in macaques and humans with tuberculosis. Science Translational Medicine 6, 265ra167 (2014). https://doi.org:10.1126/scitranslmed.3009500

53 Ankrah, A. O. et al. PET/CT imaging of Mycobacterium tuberculosis infection. Clinical and Translational Imaging 4, 131–144 (2016). https://doi.org:10.1007/s40336-016-0164-0

54 Peña, J. C. & Ho, W.-Z. Monkey models of tuberculosis: lessons learned. Infect Immun 83, 852–862 (2015). https://doi.org:10.1128/IAI.02850-14

55 Lin, P. L. et al. Sterilization of granulomas is common in active and latent tuberculosis despite within-host variability in bacterial killing. Nature Medicine 20, 75–79 (2014). https://doi.org:10.1038/nm.3412

56 Sasindran, S. & Torrelles, J. Mycobacterium Tuberculosis Infection and Inflammation: what is Beneficial for the Host and for the Bacterium? Frontiers in Microbiology 2, 2 (2011).

57 Malherbe, S. T. et al. Persisting positron emission tomography lesion activity and Mycobacterium tuberculosis mRNA after tuberculosis cure. Nature Medicine 22, 1094–1100 (2016). https://doi.org:10.1038/nm.4177

58 Sánchez-Montalvá, A. et al. Usefulness of FDG PET/CT in the management of tuberculosis. PloS one 14, e0221516 (2019). https://doi.org:10.1371/journal.pone.0221516

59 Limsrivilai, J. et al. Validation of models using basic parameters to differentiate intestinal tuberculosis from Crohn’s disease: A multicenter study from Asia. PloS one 15, e0242879 (2020). https://doi.org:10.1371/journal.pone.0242879

60 Weinstein, E. A. et al. Imaging Enterobacteriaceae infection in vivo with 18F-fluorodeoxysorbitol positron emission tomography. Science Translational Medicine 6, 259ra146-259ra146 (2014). https://doi.org:10.1126/scitranslmed.3009815

61 Carter, S. G. & Karl, D. W. Inorganic phosphate assay with malachite green: An improvement and evaluation. Journal of Biochemical and Biophysical Methods 7, 7–13 (1982). https://doi.org:10.1016/0165-022X(82)90031-8

62 Via, L. E. et al. Infection dynamics and response to chemotherapy in a rabbit model of tuberculosis using [18F] 2-fluoro-deoxy-D-glucose positron emission tomography and computed tomography. Antimicrobial Agents and Chemotherapy 56, 4391–4402 (2012).

63 Via, L. E. et al. Differential virulence and disease progression following Mycobacterium tuberculosis complex infection of the common marmoset (Callithrix jacchus). Infect Immun 81, 2909–2919 (2013). https://doi.org:10.1128/iai.00632-13

64 White, A. G. et al. Analysis of 18FDG PET/CT Imaging as a Tool for Studying Mycobacterium tuberculosis Infection and Treatment in Non-human Primates. J Vis Exp, 56375 (2017). https://doi.org:10.3791/56375

65 Brugarolas, P. et al. Development of a PET radioligand for potassium channels to image CNS demyelination. Scientific Reports 8, 607 (2018). https://doi.org:10.1038/s41598-017-18747-3

66 Riemann, B., Schäfers, K., Schober, O. & Schäfers, M. Small animal PET in preclinical studies: opportunities and challenges. Q J Nucl Med Mol Imaging 52, 215 (2008).

