## Supplementary Information for "Distributable, Metabolic PET Reporting of Tuberculosis"

### SUPPORTING INFORMATION

### Supplementary Materials and Methods

Unless otherwise noted, all chemicals and solvents were purchased from Sigma-Aldrich (Milwaukee, WI, USA), Fisher Scientific (Hanover Park, IL, USA), Sigma-Aldrich UK, Alfa Aesar, Fisher UK, Carbosynth or Acros and used without further purification. Columns and Sep-Pak cartridges used in this synthesis were obtained from Agilent Technologies (Santa Clara, CA, USA) and Waters (Milford, MA, USA), Phenomenex UK respectively. Sep-Pak was conditioned with 5 mL ethanol. Analytical HPLC analyses for radiochemical work were performed on an Agilent 1200 Series instrument equipped with multi-wavelength detectors. Mass spectra (MS) of decayed [$^{18}$F]FDT solution were recorded on a 6130 Quadrupole LC/MS, Agilent Technologies instrument equipped with a diode array detector. LC-MS analysis of fluorine-18 labeled trehalose was performed on Agilent 1260 HPLC system coupled to an Advion expression LCMS mass spectrometer with an ESI source. The LC inlet was Agilent 1200 series chromatographic system equipped with 1260 quaternary pump, 1260 Infinity autosampler, 1290 thermostatted column compartment, and radiation detector. Column flow was split (1:4) between the mass spectrometer and the radiation detector. Instrument control and data processing performed using Advion's Mass Express and Quant Express Software.

#### General Biology Experimental Procedures

All biological equipment and prepared biological solutions, such as media, plastic and glassware, used in the handling of microbial cultures were autoclaved at 121 °C. All biological work was performed in a BASSAIRE Class II Laminar Flow Cabinet. Bacterial cultures were incubated in Innova® 42 incubator Shaker Series. Centrifugation was performed in Avanti Jxn-30 Centrifuge for cell harvesting and post sonicated cell cultures, and Megafuge Centrifuge was used for protein concentration and DNA samples. Micro centrifugation was performed on Eppendorf Centrifuge 5424 R. Gel electrophoretic analysis was performed on the following equipment: Invitrogen XCell SureLockTM and BioRad Power Pac Basic (200 V used, 50 min.) with using MOPS/MES as buffer.

**Preparation of Lysogeny Broth Agar Plates**

Lysogeny broth (LB, 6.25 g) and agar (2.5 g) were both added to water (250 mL) and the mixture was autoclaved. After cooled to ~45 ºC, a stock solution of appropriate antibiotic was added and the agar was poured into plates and allowed to settle. Stored at 4 ºC.

**Transformation**

Plasmid stock (1 µL, 10 ng) and cell stock (20 µL) mixed in 14 mL polyene tube and incubated on ice for 30 min, the mixture was then heat shocked to 42 °C for 45 s then incubated on ice for 2 min. Super optimal broth with catabolic repression (SOC) medium (200 µL) was added and the mixture was incubated at 37 °C for 1h. The mixture was plated on agar plates with the appropriate antibiotic (2 plates, one with 100 µL one with 25 µL). Plates were then incubated overnight (37 °C).

**DNA Preparation**

A single colony was picked and placed into 5 mL of lysogeny broth (LB) with the appropriate antibiotic. The culture was shaken at 37 °C overnight. Cells were pelleted by centrifugation (4700 rpm, 20 min, 4 °C) then DNA preparation carried out according to the instructions provided with the QIAGEN QIAPrep spin miniprep kit. DNA concentration was determined by absorption on a Nanodrop and Sanger sequencing performed by Source Bioscience. Sequence analysis was performed by using source bio edit.

**SDS-PAGE**

Sodium dodecyl sulfate – polyacrylamide gel electrophoresis (SDS-PAGE) was carried out to check the purity of all proteins. These were carried out using 4 – 12% bis-Tris polacrylamide gels with either MOPS or MES buffer. To the protein sample (8 μL) was added 2 × SDS loading buffer (3 μL 80 mM Tris-HCl buffer pH 6.8, 8% v/v glycerol, 40 μM bromophenol blue, 80 mM SDS, 2% β-mercapethanol). The solution was then heated to 95 ºC for 10 min. The sample was then applied to a well in the gel. The gel was then run at 200 V for 1 h and then visualised by the addition of Coomassie InstantBlue dye (Gentaur) and gentle stirring for 2 h at room temperature.

**Western Blots**

SDS-PAGE was done as before, but using a Pre-Stained protein ladder and the gel was not stained with InstantBlue dye. Instead, the gel was electroblotted onto a membrane for 7 min using an iBlot machine. The membrane was then washed with 50 mM Tris-HCl, 150 mM NaCl and 0.05% Tween 20, pH 7.6 (TBS-T + 5% BSA w/v , 45 mL) for 1 h. The membrane was further washed with TBS-T + 5% BSA w/v, 45 mL and monoclonal anti-polyHistidine-alkaline phosphatase antibody produced in mouse (Sigma, 16 μL). The membrane was then rocked for 4 h at room temperature. After this time, the solution was removed and the membrane washed with TBST buffer (3 × 20 mL). BCIP/NBT alkaline phosphate substrate (Sigma Aldrich, 4 mL) was added and the membrane was gently rocked until protein bands could be seen (~15 min). The membrane was then washed with water.

**Protein Mass Spectrometry**

Various machines used for protein LC-MS. In all cases solvent A was water and solvent B was acetonitrile with both containing 0.1% formic acid.

Set-up 1: Waters LCT Classic coupled to a Shimadzu 20 Series HPLC using a Thermo Proswift (250 x 4.6 mm x 5 μm) column. Electrospray source parameters were as follows: capillary voltage 3000 V; cone voltage 25 V. Spectra were calibrated to at least 17 matched peaks of the multiply charged ion series of equine myoglobin run under equivalent conditions. The gradient program is shown below; a flow rate of 0.4 ml/min was used throughout (see also Supplementary Table S3).

Set-up 2: Waters Xevo G2-QS QToF MS coupled to a Water Acquity UPLC using a Thermo Proswift (250 x 4.6 mm x 5 μm) column. Electrospray source parameters were as follows: capillary voltage 3000 V; cone voltage 20 V. Spectra were calibrated through use of an internal lock-spray. The gradient program is shown below; a flow rate of 0.3 ml/min was used throughout (see also Supplementary Table S4).

Nitrogen was used as the nebulizer and desolvation gas at a total flow of 600 l $hr^{-1}$. Spectra were calibrated using a calibration curve constructed from a minimum of 16 matched peaks from the multiply charged ion series of equine myoglobin obtained at a cone voltage of 35 V. Data was processed using MassLynx software (v. 4.1 from Waters) according to the manufacturer's instructions.

**Circular Dichroism (CD) Spectroscopy**

For CD measurement, protein was buffer exchanged into phosphate buffer (10 mM potassium phosphate, pH 7.5). Measurements were made at room temperature using a Chirascan CD-spectrometer (Applied Photophysics) using 0.1 cm quartz cuvette. Background readings were taken with using 0.2 mL of buffer only, followed by protein samples measured at concentrations of 0.2 mg/mL. Scans were run using 0.5 s per time point and 0.5 nm increments, scanning from 185 to 280 nm. Data reported is the mean average of 3 results.

**Liquid Chromatography-mass spectrometry for Small Molecule Analysis**

Liquid chromatography–mass spectrometry (LC–MS) for quantification of trehalose and glucose was performed on the LC-MS Quattro instrument coupled with agilent technologies HPLC system, using a TOSOH TSKgel Amide-80 column (4.6 mm x 250 mm, 5µM, 80 Å). Solvent A ($CH_3CN$) and ($H_2O$) with a volume ratio of 70:30 were used as the isocratic mobile phase at a flow rate of 1.0 mL / min. The MS file was programme as 20 min for each sample. The electrospray source was operated with a capillary voltage of 3.0 kV and a cone voltage of 20 V. Nitrogen was used as the nebulizer and desolvation gas at a total flow of 500 $Lh^{-1}$. The data was acquired by single ion recording (SIR) of m/z 341 for trehalose and m/z 179 for glucose in negative mode and processed using MassLynx software (v. 4.1 from Waters) according to the manufacturer's instructions. See Supplementary Table S9 for the LC-MS methods employed in the detection of FDT.

**High Performance Liquid Chromatography (HPLC) for Enzyme Reaction Analysis**

All of the enzyme reactions were analysed by HPLC for UDP-glucose consumption and UDP release. For the analysis, few different stationary and mobile phases were used to optimise the separation between UDP and UDP-glucose (summarised in Supplementary Table S5). Method 3 was selected as a method of choice because it gave the best separation between the two phosphate sugars.

**Expression, Purification and Characterisation of OtsA *E. coli***

pET-22b(+) C-Terminal-His vector containing gene for OtsA was transformed into BL21 (DE3) cells using standard transformation protocol. The transformed cells were used to inoculate an overnight culture in 100 mL of freshly prepared LB medium containing 100 μL of Ampicillin stock solution. The culture was incubated overnight at 37 ˚C. The overnight culture was used to isolate plasmid DNA using the standard QIAGEN miniprep kit protocol. The gene construct was confirmed by sequencing.

In a typical expression experiment performed on a 800 mL LB medium scale, transformed cells were grown overnight at 37 ˚C in 100 mL of freshly prepared LB containing 100 μL of Ampicillin stock solution (100 mg/mL). 4 mL of overnight culture was used to inoculate 800 mL of freshly prepared LB media containing 800 μL of Amp stock solution. The culture was incubated at 30 ˚C with shaking. OD600 was measured from time to time. Once the OD ~ 0.45, protein expression was induced by adding 4 mL of 1 M IPTG solution to a final concentration of 0.5 M. The culture was left to grow overnight at 30 ˚C for 18 hours. The culture was harvested by centrifugation (8000 rpm, 10 minutes, 4 ˚C). Cell pellet was suspended in binding buffer (50 mM Hepes, 200 mM NaCl, 10 mM imidazole, pH 7.5) and one tablet of Complete protease inhibitor cocktail (Roche), DNAase and lysozyme was added. The mixture was stirred vigorously at 4 ˚C for 2 hours. The cells were lysed via sonication. The suspension was then centrifuged (14000 rpm, 30 min) to remove cell debris. The supernatant was then filtered using a 0.45 μm syringe filter. The filtered supernatant was applied to a pre-equilibrated (using binding Buffer) GE Life Sciences 5mL HisTrap Ni-NTA column. Purification was done at 4 ˚C. The column was washed by 15 column volumes of binding buffer, followed by a linear gradient to 100 % Elution Buffer (50 mM Hepes, 200 mM NaCl, 500 mM Imidazole, pH 7.5) over 16 column volumes. Eluted protein fraction were analysed using SDS-PAGE analysis.

**Expression, Purification and Characterisation of His-MBP-OtsB from *E. coli***

In a routine restriction digestion reaction for vectors, 2 µL of each restriction enzyme (NdeI & XhoI), 3 µL of NEB buffer 4 (10x), 1 µg of vector (pDB-His-MBP) were taken in an eppendorf tube. dd H2O was added to reaction mixture to make reaction volume 30 µL. The reaction mixture was incubated at 37 ˚C for 4 h. For insert, similar reaction was set-up with 3 µL of OtsB gene in pET-29b(+) vector. Digested vector and insert was loaded onto 1% Agarose gel. The gel was run at 110 V for 40 minutes.

Restriction-digested products were extracted from agarose gel using the standard QIAquick gel extraction kit protocol. Under UV transilluminator, gel fragments containing restriction-digested products were excised using a clean, sharp scalpel. Gel were transferred to eppendorf tubes, weight of gel was taken. 3 volumes of QG buffer were added to 1 volume of gel. The sample was incubated at 50 ˚C for 10 minutes with occasional vortexing. 1 gel volume of isopropanol was added and was mixed by inverting the eppendorf tube 4-6 times. The sample was decanted into a spin column and was centrifuged for 1 minute. The flow through was discarded. 500 µL of QG buffer was added to the column and centrifuged for 1 minute. The supernatant was discarded and 750 µL of buffer PE was added to the column. The spin column was centrifuged for 1 minute. The supernatant was discarded, to remove the residual wash buffer; column was centrifuged further for 1 minute. The spin column was placed in a new Eppendorf tube and 50 µL of dd H2O was placed at the centre of column. The column was left to stand for 4-5 minutes, after which it was centrifuged for 1 minute. The DNA was stored at −20 $^{o}$C for further manipulation. Concentration of products was calculated using Nanodrop.

For ligation, 50 ng of vector was combined with a 3-fold molar excess of PCR product. The volume was adjusted to 20 µL with dd H2O. 2 µL of 10 x T4 ligation buffer was added and

mixed thoroughly. 1 µL of Quick T4 DNA Ligase was added and mixed thoroughly. Reaction mixture was centrifuged briefly and was incubated at 25 C for 5 minutes. The ligated product was stored at - 20 °C. Transformation of the ligated product was done in XL10 Gold competent cells through standard transformation protocol
using 2 µL of ligated reaction mixture. Transformed cells were plated on LB-Agar/Kan plates and incubated at 37 ˚C overnight.
Single colonies of transformed cell were used to inoculate overnight culture. Plasmid DNA was isolated from the overnight culture using standard miniprep protocol. The successful sub-cloning was verified by sequencing.
pDB-His-MBP vector containing gene for OtsB was transformed into BL21 (DE3) cells using standard transformation protocol. A single transformed cell was used to inoculate a 100 mL starter culture of freshly prepared LB medium containing 100 µL of Kanamycin stock solution (50 mg/mL). The culture was incubated at 30 ˚C for 16 hours. 8 mL of starter culture was used to inoculate 800 mL of LB medium containing 800 µL of Kanamycin stock solution. The culture was incubated at 37 ˚C with shaking. OD600 was measured from time to time. Once OD600 reaches close to 0.6. Expression was induced by adding 800 µL of 1 M IPTG solution. The culture was incubated at 30 ˚C with shaking for 16 hours.
The culture was harvested by centrifugation (8000 rpm, 10 minutes, 4 °C). Cell pellet was suspended in binding buffer (50 mM Tris, 200 mM NaCl, 20 mM Imidazole, pH 7.5) containing Complete protease inhibitor cocktail (Roche), DNAase and lysozyme. The mixture was stirred vigorously at 4 ˚C for 2 hours. The cells were lysed via sonication using program P9 consisting of 0.5 sec on, 0.5 sec off pulses for 30 sec followed by 1 minute of cooling at 40% amplitude. The sonication was done for 5 cycles. The suspension was centrifuged (20000 rpm, 40 min) to remove cell debris. The supernatant was then filtered using a 0.45 µm syringe filter. The filtered supernatant was applied to a pre-equilibrated (Binding Buffer) GE

Life Sciences 5ml HisTrap HP column. Purification was done at 4 ˚C. The column was washed by 10 column volumes of binding buffer, followed by a linear gradient to 100 % Elution Buffer (50 mM Tris, 200 mM NaCl, 500 mM imidazole, pH 7.5) over 15 column volumes. Purification was done at a flow rate of 5 mL/min. Eluted protein fraction were analysed using SDS-PAGE analysis. Buffer exchange was done using dialysis tubing with MWCO 10,000 Da. Fractions containing the ca. 72 KDa protein were pooled and dialyzed thrice against 4 liters of 50 mM Tris, pH 7.5 and stored at 4 ˚C. Protein were concentrated using Vivaspin with MWCO 30,000 Da to a final concentration of 12.55 mg/mL. The protein was further stored at -20 $^{o}$C and characterised by SDS-PAGE.

**Expression, Purification and Characterisation of TreT *P. horikoshii* Enzyme**

The DNA plasmid from group plasmid bank was verified by sequencing from source bio-science. The plasmid encoded a C-terminal 6 x Histidine tag (His-tag) on the protein. The plasmid was transformed into *E.coli* (BL21 DE3) cells and plated on LB agar containing 50 mg/mL kanamycin. One of the resulting grown colony was inoculated into 10 mL LB media containing 10 µL of kanamycin (50 mg/mL) solution and left incubating overnight (37 °C, 200 rpm).

To express the TreT protein, 10 mL grown culture was inoculated into 1 L of LB media containing 1 mL of kanamycin solution (50 mg/mL) and left incubating (37 °C, 200 rpm). $OD_{600}$ was measured every one hour and once the inoculated culture reached the mid-log phase, it was induced with 1 mL of IPTG solution (final concentration = 1 mM) and grown in shaking incubator overnight (37 °C, 200 rpm).

Cells were then harvested by centrifugation (4700 rpm, 20 min, 4 °C) and the pellet was resuspended in 30 mL lysis buffer (300 mM NaCl, 50 mM $NaH_2PO_4$, 20 mM imidazole). One protein inhibitor tablet, 25 mg lysozyme and 5 mg of DNase was also added into the lysate and lysis reaction was carried out on rotating platform (60 min, 4 °C). Lysed cells were sonicated (5 x 30 s, 30 s pulse on, 60 s pulse off, 75% amplitude). Sonicated cells were centrifuged (21000 x g, 20 min, 4 °C) to clarify the lysate. Supernatant was then filtered (0.45 μm and 0.2 μm filter). Next supernatant was loaded on a 50 mL FPLC loop and purified by Ni-NTA chromatography using 5 mL His-trap HP column and eluted with 15 column volumes of elution buffer (300 mM NaCl, 50 mM $NaH_2PO_4$, 250 mM imidazole). Protein samples were concentrated using Vivaspin at 10 kDa MWCO (Sartorius). Elution buffer was exchanged for 300 mM NaCl, 50 mM HEPES, pH 7.5. Protein samples were then characterised by SDS-PAGE to verify protein purity and LC-MS to confirm the mass of the protein.

**Expression, Purification and Characterisation of TreT *T. tenax* Enzyme**

2 x 1 L cultures of TreT *T. tenax* protein were expressed in BL21 (DE3) cells. Expression was carried out by inoculating 10 mL cultured BL21 (DE3) transformed with plasmid DNA into 1 L of LB media. 1 mL of Kanamycin sulphate solution (50 mg / mL) was used as the resistance antibiotic. Both the flasks were shaken at 200 rpm and 37 °C for 3 hours when the $OD_{600}$ value reached to 0.6. At this stage protein induction was carried out by using 1.0 mL of IPTG solution (final concentration = 1 mM), post induction both flasks were then kept overnight in the shaking incubator. Cells were then harvested by centrifugation (4700 rpm, 20 min, 4 °C), and the pellet was resuspended in 30 mL lysis buffer (300 mM NaCl, 50 mM $NaH_2PO_4$, 20 mM imidazole). One protein inhibitor tablet, 25 mg lysozyme and 5 mg of DNase was also added into the lysate and lysis reaction was carried out on rotating platform (60 min, 4 °C). Lysed cells were sonicated (5 x 30 s, 30 s pulse on, 60 s pulse off, 75% amplitude). Sonicated cells were centrifuged (21000 x g, 20 min, 4 °C) to clarify the lysate.

Supernatant was then filtered (0.45 µm and 0.2 µm filter). Next supernatant was loaded on a 50 mL FPLC loop and purified by Ni-NTA chromatography using 5 mL His-trap HP column and eluted with 15 column volumes of elution buffer (300 mM NaCl, 50 mM $NaH_2PO_4$, 250 mM imidazole).

FPLC fractions were then analysed by SDS-PAGE. Fraction containing potential protein were concentrated using Vivaspin at 10 kDa MWCO (Sartorius) and centrifuging at 8000 x g. Protein concentration was determined by nanodrop and characterisation was done with mass spectrometry and SDS-PAGE.

**Plasmid Design and Trial Expression of OtsAB Fusion Enzyme**

N-terminus His-tagged OtsAB fusion protein plasmid DNA was transformed into BL21 (DE3) cells. The grown colonies were inoculated into the LB media (10 mL) containing 10 µL of ampicillin (100 mg/mL) solution incubated overnight in the shaking incubator (37 °C, 200 rpm). Small scale expression trial was carried out by pipetting 250 µL of grown culture in 20 mL of LB media in 6 different 50 mL falcon tube. 20 µL of ampicillin solution was also added. Tube 1 & 2 were used as control (0.4 $OD_{600}$ and 0.6 $OD_{600}$); tube 3 & 4 were induced with 0.5 mM and 1 mM IPTG solution when $OD_{600}$ value was 0.4 and 0.6 respectively. A similar process was repeated for tube 5 & 6. 1 mL of sample was taken from each tube at 0 min, before adding IPTG solution, 4 h, 16 h, 24 h and 48 h post IPTG induction.

Cells were then harvested from liquid culture by centrifugation (10,000 x g, 10 min, 4 °C). The cell solution was decanted and the pellet was allowed to drain completely. 100 µL of bugbuster reagent was added into each of the sample and the cell pellet was completely re-suspended by vortexing slowly. Cells were then incubated at a shaking platform for 20 min at room temperature. Insoluble cell debris was removed by centrifugation (16000 x g, 20 min, 4 °C). Supernatant was transferred into a fresh tube and analysed by SDS-PAGE.

**Large Scale Expression of OtsAB Fusion Protein**

N-terminus His-tagged OtsAB fusion protein plasmid DNA (2 µL) was transformed into BL21 (DE3) cells (20 µL). The grown colonies were inoculated into the LB media (10 mL) containing 10 µL of ampicillin solution (100 mg/mL) incubated overnight in the shaking incubator (37 °C, 200 rpm). Large scale expression was carried out by inoculating 10 mL of grown culture in 1000 mL of LB media in 2 different 2L conical flasks. 1000 µL of ampicillin solution (100 mg/mL) was also added. Both flasks were then incubated in the shaking incubator (37 °C, 200 rpm) until the $OD_{600}$ reached 0.4. Cells were then incubated at 15 °C for further 30 min and then induced by adding IPTG solution (1 mL, 1 mM) and incubated for further 24 h at 15 °C.

After 24 h cells were harvested by centrifugation (9000 rpm, 20 min, 4 °C) and lysed with lysis buffer (50 mM Tris, 50 mM imidazole and 150 mM NaCl, pH 7.5) for 30 min at room temperature. Lysis was carried out by adding lysozyme (25 mg), DNase (5 mg) and 1 protein inhibitor tablet. Post lysis cells were sonicated (60% amplitude, 6 cycles of 30 s pulse on, 60 s off). Sonicated cells were then centrifuged to get rid of cell debris (20000 rpm, 20 min, 4 °C) and filtered through 0.45 µm and 22 µm filter and loaded onto FPLC loop for purification using His-trap HP 5 mL column as stationary phase and eluted with 15 column volumes of elution buffer (50 mM Tris. 150 mM NaCl, 500 mM imidazole).

FPLC fractions were then visualised by SDS-PAGE method and fraction containing potential protein were concentrated through Vivaspin (5 mL, 30 kDa MWCO) by centrifugation (5000 x g, 15 min, 4 °C). Protein samples were then characterised by LC-MS.

**Enzymatic Reaction of OtsAB fusion Protein**

To test the protein activity enzymatic reaction was carried out. 20 µL Glucose-6-phosphate (20 mM); 20 µL of UDP-glucose (20 mM), 1.25 µL of manganese chloride (2.5 mM) and 20 µL of enzyme (0.23 µM) were added into 1.5 mL Eppendrof tube. The total reaction volume was adjusted to 100 µL by adding Tris/NaCl buffer (50 mM Tris; 150 mM NaCl, pH 7.5). Reaction was started by adding the enzyme solution and stopped by placing on ice and quenched by adding 20 µL acetonitrile solution. Reaction was carried out on the thermoshaker (37 °C, 700 rpm) and continued for 24 h.

**Detection of Trehalose by LC-MS for TreT and OtsAB Fusion enzyme**

In order to detect the trehalose LC-MS method was used as described in the general experimental condition. Standard solutions for glucose and trehalose were analysed along with the samples. For TreT reaction UDP-glucose was used in excess and 8 mM glucose was used as a substrate. Enzyme concentration, metal ion concentration and buffer amounts were kept the same as in the case UDP-Glucose and UDP reaction.

For OtsAB fusion reaction, total reaction volume was 100 µL. In 1.5 mL Eppendorf tube were added: glucose-6-phosphate 2 µL (10 mM), $MnCl_2$ 1.25 µL (2.5 mM) and 0.23 µM enzyme was used. UDP-Glucose concentration was maintained at 50 µL (50 mM). Reaction was made up to 100 µL by adding Tris/HCl buffer solution (50 mM, pH 7.5).

**Reaction Comparison between TreT homologues**

To demonstrate the yield of FDT from TreT enzyme reactions, we tested the reaction for both homologues. Equimolar amount of starting substrate and enzyme concentration was used (see

Supplementary Table S6). All the other parameters and reagents concentration was kept constant so that yield for FDT formation from each enzyme system can be compared.

In summary reaction volume was maintained to 500 μL and HEPES buffer (50 mM HEPES, pH 7.5, 200 mM NaCl and 10 mM $MgCl_2$) was used to make up the required volume. Reactions were initiated by adding the enzyme and then incubated at 37 °C, 700 rpm on the thermoshaker. 150 μL of reaction samples were taken after 30 min, 60 min and 120 min respectively from each reaction mixture. Reactions were stopped by incubating the reaction mixture at 100 °C for 5 min and further diluted to 500 μL with $D_2O$ solution, centrifuged and stored for the analysis.

Sample analysis was carried out using 500 MHz NMR spectrometer (AVB500) and $^{19}$F-NMR signal for the formation of FDT were sought as compared to FDG peaks in all the reaction mixtures.

**Evaluation of Reaction Reversibilty with TreT *P. horikoshii***

To test the reversibility of TreT *P. horikoshii*, we set-up small scale reactions from 1-10 mM of [$^{19}$F]FDT with fixed UDP concentration of 20 mM. In 1.5 mL Eppendorf tube were added the following: UDP 20 mM, [$^{19}$F]FDT 1 mM, 2 mM, 5 mM and 10 mM with a further 1 mM control reaction. Reaction volume was made up to 100 μL with HEPES buffer solution (50 mM HEPES, 200 mM NaCl, 10 mM $MgCl_2$, pH 7.5). Reactions were initiated by adding TreT enzyme and incubated at 37 °C for 2 h. A further reaction with higher amount of UDP substrate (80 mM) and [$^{19}$F] FDT (20 mM) was also set-up as above. Reactions were analysed by [$^{19}$F]-NMR spectroscopy.

**Enzyme production scale-up in Sf9 cells expression**

For higher level expression of OtsA and OtsB proteins, the Bac-to-Bac® Baculovirus Expression System was selected. The original coding sequence of *E. coli* OtsA (CAA48913.1) and OtsB (CAA48912.1) was optimized according to *S. frugiperda* MNPV codon usage. The DNA fragment was then cloned into the *EcoRV* restriction enzyme site of the pUC57 vector. Then the plasmids pUC57-OtsA and pUC57-OtsB and pFastBac Dual vector were digested with *BamHI* and *NotI*, respectively, and ligated to create the recombinant plasmids of OtsA (pFastBac-OtsA) and OtsB (pFastBac-OtsB). The bacmid DNAs of OtsA (Bacmid-OtsA) and OtsB (Bacmid-OtsB) were generated by transformation of the recombinant plasmids pFastBac-OtsA and pFastBac-OtsB into the DH10Bac™ *E. coli*, respectively. Correct insertions of the OtsA and OtsB genes in the bacmid DNA were confirmed by PCR analysis using pUC/M13 standard primers: 5-CCCAGTCACGACGTTGTAAAACG-3 (forward) and 5-AGCGGATAACAATTTCACACAGG-3 (reverse).

Based on the bacmid DNAs, the recombinant baculovirus of OtsA and OtsB were made by transfection of Sf9 cells. Culture and preparation of Sf9 cells were performed according to the protocol provided by manufacturer. A Sf9 cell stock with a viability >95% and a density of $1.5 \times 10^6 – 2.5 \times 10^6$ cells/mL was prepared before proceeding to transfection.

**Experimental procedures for generating P1 baculovirus for OtsA and OtsB**

The P1 baculovirus stocks of OtsA and OtsB were generated using Sf9 cells in 6-well culture plates. Briefly, 2 ml of Sf9 cells were added to each well of two 6-well plates with the cell number of $8 \times 10^5$ in each well. Cells were allowed to attach for 15 minutes at room temperature. Transfectin II was used as the transfection reagent and the transfection experiments were performed according to the protocol provided by manufacturer. Different amounts of bacmid DNAs of OtsA and OtsB were used. After transfection, the Sf9 cells were further cultured at 27 °C for 3 days, then the P1 baculovirus (in supernatant) was harvested by centrifugation. The P1 baculovirus were found in the culture medium and the Sf9 cell pellets were used to check protein expression. Fetal bovine serum (FBS) was then added to each P1 baculovirus solution to reach a final concentration of 2%. The P1 baculovirus stock solutions were then stored at 4 °C, protected from light.

Amplification of P1 baculovirus of OtsA and OtsB: In order to generate baculovirus stock for large scale protein expression and to increase virus titer, P2 and P3 baculovrius stocks of OtsA and OtsB were prepared from their P1 baculovirus. Briefly, a total of 150 mL of suspended Sf9 cells with a cell density of $2.0 \times 10^6$ cells/mL was infected by P1 baculovirus of OtsA or OtsB, respectively. The 150 mL infection culture composition was: 1% P1 baculovirus of OtsA or OtsB, 1% FBS, 1% ethanol, Sf9 cells with a density of $2.0 \times 10^6$ cells/mL. After infection at 27 °C with shaking (150 rpm) for 72 hours, the P2 baculovirus (in culture medium) was harvested by centrifugation at 1100 rpm for 10 minutes at room temperature. FBS was added to a final concentration of 2%, then the solution was filtered by syringe filter (0.45 μm) and stored at 4 °C, protected from light. Similar infection procedures were performed to prepare P3 baculovirus, except using P2 baculovirus stock solution for cell infection.

**Expression and Purification of C-terminally tagged, recombinant OtsA**

Insect cells were grown in serum-free medium to a density of $2 \times 10^6$ cells/mL and a cell viability of greater than 95%. The cell culture (1 L) was transfected with the viral stock, 2.8 x $10^6$ cells/mL. At 72 h post-infection, the cells were harvested by centrifugation. After freeze/thaw cycle, the pellet was resuspended in lysis buffer (100 mM HEPES, 200 mM NaCl, 10 mM imidazole, 10 mM $MgCl_2$, 1 mM βME). DNase 3 mg and 1x Complete®

protease inhibitor cocktail was also added into the protein pellet and lysed using a homogenizer. Cell debris was removed by centrifugation and the cell free extract passed through a 0.45 μm membrane filter. The filtrate was applied to a Ni–NTA column equilibrated in lysis buffer. After removal of the cell debris by centrifugation, the protein of interest was purified with a linear gradient of 10–300 mM imidazole in 100 mM HEPES, 200 mM NaCl, 10 mM $MgCl_2$, 1 mM βME, pH 7.5, using Ni-NTA. Protein samples were exchanged into 100 mM Hepes, 150 mM NaCl, 10 mM $MgCl_2$, pH 7.5, using a Hiload™ 26/60 Superdex™ 200 desalting column. The sample was passed through an Acrodisc Mustang E membrane (Pall #MSTG25E3) to reduce endotoxin to 1.9 EU/mL. After the addition of 10% glycerol, the aliquots containing the recombinant enzyme were flash frozen, and stored at −80 °C.

**Expression and purification of recombinant OtsB**

Insect cells were grown in serum-free medium to a density of $2 \times 10^6$ cells/mL and a cell viability of greater than 95%. The cell culture (1 L) was transfected with the viral stock, $2.0 \times 10^6$ cells/mL. At 72 h post-infection, the cells were harvested by centrifugation. After freeze/thaw cycle, the pellet was resuspended in lysis buffer (50 mM Tris-HCl, 200 mM NaCl, 5 mM $MgCl_2$, 30 mM imidazole, 1 mM βME). DNase 3 mg, 1x Complete® protease inhibitor cocktail was also added in the pellet and lysed using a homogenizer. Cell debris was removed by centrifugation and the cell free extract passed through a 0.45 μm membrane filter. The filtrate was applied to a Ni–NTA column equilibrated in lysis buffer. After removal of the cell debris by centrifugation, the protein of interest was purified with a linear gradient of 10–300mM imidazole in 50 mM Tris-HCl, 200 mM NaCl, 5 mM $MgCl_2$, 1 mM βME, pH 7.5, using Ni-NTA. The fractions with potential protein were concentrated using Vivaspin (5 MWCO) and then desalted on Hiload™ 26/60 Superdex™ 200 with 50 mM Tris-HCl, 5 mM

$MgCl_2$, 100 mM NaCl, pH 7.5. The sample was passed through an Acrodisc Mustang E membrane (Pall #MSTG25E3) to reduce endotoxin to 1.9 EU/mL. After the addition of 10% glycerol, the aliquots containing the recombinant enzyme were flash frozen, and stored at −80 °C. SDS–PAGE analysis revealed the presence of a protein with the correct molecular weight.

**Small-scale synthesis of [$^{19}$F]FDT using 3-Enzyme, One-pot System**

To mimic 'hot' FDT synthesis, we tested 'cold' FDT synthesis from 0.138 mmol – 5.520 mmol scale. Reagent amounts are detailed in Supplementary Table S8. In summary, in a 1.5 mL Eppendorf tube were added the following: FDG (various concentrations, see Supplementary Table S8), ATP 0.72-7 mM, UDP-Glc 20 mM, KCl 100 mM, hexokinase 1 mg, OtsA 1 mg and OtsB 0.1 mg in 50 mM HEPES buffer solution containing 100 mM NaCl and 10 mM $MgCl_2$. The final reaction volume was adjusted to 1 mL in all cases. The reaction was incubated at 37 °C, 400 rpm and monitored by $^{19}$F-NMR and LC-MS. Full conversion was achieved in 45 min. A control sample of FDG with no enzyme added was also analyzed.

**General Chemistry Experimental Procedures**

Proton nuclear magnetic resonance spectra ($^1$H NMR) were recorded on a Bruker AVX 500 (500 MHZ), a Bruker AVB 500 (500 MHz) or a Bruker AVB 400 (400 MHZ) spectrometer as indicated. Carbon nuclear magnetic resonance ($^{13}$C NMR) spectra were recorded on the same spectrometer as described above. $^{19}$F-NMR spectra were recorded on a Bruker AVX500, AVB500, AV600 and AVB400. NMR spectra were fully assigned using COSY and HSQC. All chemical shifts are quoted on the δ scale in ppm using residual solvents as the internal standards. Coupling constants (*J*) are reported in Hz with the following splitting abbreviations: s = singlet, d = doublet, t = triplet, q = quartet, and m = multiplet.

Infrared (IR) Spectra were recorded on a Bruker Tensor 27 Fourier Transform Spectrometer using KBr discs for solids and crystals. Absorption maxima (νmax) are reported in wave number ($cm^{-1}$) and classified as strong (s) or broad (br).

Low-resolution mass spectrometry (LRMS) were recorded using an agilent 6120 Quadrupole spectrometry using electrospray ionisation (ESI) in either positive or negative mode. High resolution mass spectra (HRMS) were recorded on a Thermo Orbitrap Exactive mass spectrometer. The instrument is calibrated with a standard calibration mix from thermo before the run. Liquid Chromatography – Mass spectrometry (LC-MS) analysis was carried out on a Waters Quattro Micro – MS with electrospray ionization operating in negative mode (ESI) interface with Agilent HPLC system and Waters 2777 auto sampler with 4-valve injector module. MS analysis was under the control of micro mass Masslynx 4.1 software.

Thin layer chromatography (TLC) was carried out using aluminium backed sheets coated with 60F254 silica gel (Merck). Visualization of the silica plates was achieved using a UV lamp (λmax = 254 nm), and/or ammonium molybdate 5% in 2M $H_2SO_4$. Column chromatography was carried out using Geduran ® Si 60 silica gel (Merck). Mobile phases are reported in ratio of solvents (e.g. 4:1 petrol/ethyl acetate).

### Radio-HPLC Method

Radio-HPLC analysis was carried out in a Bioscan radio-HPLC system (flow count) coupled with agilent 1200 HPLC system. Two analytical methods were used for the analysis of the depletion of starting material and formation of the product. In method 1, initially isocratic elution of 75:25 acetonitrile/water mobile phase combination was used with TSK gel normal phase HILIC column (phase amide -80, 10 cm x 4.6 mm x 5 μm) as stationary phase. The flow rate was maintained at 0.4 mL/min. This system resulted in the [$^{18}$F]FDG retention time of ~ 5.6 min and [$^{18}$F]FDT of 11.4 min. The mobile phase combination was then slightly adjusted to 70:30 acetonitrile/water and hence retention time of [$^{18}$F]FDG was 4.9 min and [$^{18}$F]FDT was ~ 8.3 min. The total run time was 15 min.

The method 2 allowed the clear separation of phosphate sugars as well as analysis of [$^{18}$F]FDG and [$^{18}$F]FDT. In This method Advanced Glycan Mapping column (4.6 x 250 mm, 2.7 μm) was used as the stationary phase. The mobile phase was the gradient of two solvent system; solvent A= 50 mM ammonium formate, pH 4.5 and solvent B= acetonitrile. Flow rate was maintained at 0.5 mL/min and the total run time was 20 min.

**Radio-TLC Method**

Radio-TLC analysis was carried out in a Bioscan AR-2000 radio-TLC scanner. Silica gel plates 150 Å Silica Gel HL 250 cm 10 x 20 cm were used as the stationary phase and the 75:25 acetonitrile/water combination was used as the mobile phase. Once the strip developed to the 10 cm mark, was scanned by the radio-TLC scanner and analysed.

**Luna-$NH_2$ Purification Method**

[$^{18}$F]FDT was purified by Sep-Pak cartridges. Sep-Pak Aminopropyl ($NH_2$) plus short cartridge (Waters, WAT020535, 360 mg) used as the stationary phase. The cartridge was pre-rinsed with 1 mL EtOH and then twice with 3 mL per wash sterile water for injection. A 10 mL syringe was attached to the cartridge with plunger removed to the 5 μm filter. 1 mL of EtOH was loaded on to the syringe.

The reaction mixture was diluted with 2 mL of EtOH. This mixture was poured into the above syringe with filter. Reaction vial was rinsed with further 1 mL of EtOH and transferred to the syringe. The contents of the syringe were filtered through 5 μm syringe tip filter to remove precipitate. Eluent was loaded onto the $NH_2$ cartridge and eluted without washing at a very low flow rate. Mixture was then concentrated first by evaporation and then by heating at

60 °C with a nitrogen flush. Finally the mixture was diluted with sterile water and filtered through sterile filter and drawn into the syringe.

**OtsA Enzyme kinetic assay**

The OtsA enzyme kinetic assay were measured through a NADH-linked GT-continued spectrophotometric assay. The decrease in absorbance of NADH at 340 nm (ε340 = 6220 $M^{-1}cm^{-1}$) was measured at 25 °C using a spectrophotometer equipped with a thermostat. To determine the kinetic parameters of UDP-Glc, various UDP-Glc cocentrations (0.5 mM, 1 mM, 2 mM, 4 mM and 10 mM) were used at the fixed G6P concentration. The G6P concentrations were fixed at 2 mM, 4 mM, 8 mM, 16 mM and 35 mM. The reaction mixture contained 100 mM HEPES (pH 7.5), 10 mM $MgCl_2$, 300 mM KCl, 1 mM potassium phosphoenolpyruvate, 300 μM NADH, 17 units of pyruvate kinase, and 24 units of lactate dehydrogenase in a total volume of 200 μL at 25 °C as well as a fixed G6P concentration and variable UDP-Glc concentration. For G6P assay, UDP-Glc concentration was fixed at 10 mM, whereas, G6P concentration varied (1 mM, 2 mM, 4 mM, 8 mM, 16 mM and 32 mM). After incubation for 5 min at 25 °C, reactions were initiated by the addition of OtsA (final concentration 83 nM).Then absorbance of the reaction mixture at 340 nm was recorded continuously.

**TreT *P. horikoshii* Enzyme Kinetic Assay**

TreT *P. horikoshii* enzyme kinetic assay were measured through a NADH-linked GT-continued spectrophotometric assay. The decrease in absorbance of NADH at 340 nm (ε340 = 6220 $M^{-1}cm^{-1}$) was measured at 37 °C using a spectrophotometer equipped with a thermostat. To determine the kinetic parameters of UDP-Glc, various UDP-Glc cocentrations (0.5 mM, 1 mM, 2mM, 4 mM and 10 mM) were used at the fixed Glc concentration. The Glc concentration was fixed at 50 mM. The reaction mixture contained 100 mM HEPES (pH 7.5), 10 mM $MgCl_2$, 300 mM KCl, 1 mM potassium phosphoenolpyruvate, 300 μM NADH, 17 units of pyruvate kinase, and 24 units of lactate dehydrogenase in a total volume of 200 μL at 37 °C as well as a fixed Glc concentration and variable UDP-Glc concentration. For Glc assay, UDP-Glc concentration was fixed at 20 mM, whereas, Glc concentration varied (0.5 mM, 1 mM, 2 mM, 4 mM, 10 mM). After incubation for 5 min at 25 °C, reactions were initiated by the addition of TreT (final concentration 662 nM).Then absorbance of the reaction mixture at 340 nm was recorded continuously.

**TreT *T. tenax* Enzyme Kinetic Assay**

In order to determine kinetic parameters ($k_{cat}$, $K_m$) of TreT *thermoproteus tenax* enzyme, UDP-Glc concentrations were varied from 0.03125 mM – 7 mM

In 1.5 mL Eppendorf tube were added glucose 50 μl (50 mM), $MgCl_2$ 20 μl (10 mM), TreT enzyme 4 μl, Hepes buffer varied volumes (50 mM, pH 7.5) to make up the total volume to 100 μl. Different volumes of UDP-glucose were added to the reaction mixture so that the final concentration of UDP-glucose in the reaction mixture was 0.01325, 0.06125, 0.125, 0.50, 1.00, 2.00, 3.00, 4.00, 5.00, 6.00 and 7.00 mM. Reactions were started by adding 4 μl enzyme and immediately placing on the thermoshaker at 70 °C. Each reaction was heated for 10 min and reactions were stopped by placing on ice and quenched by adding 20 μl acetonitrile to precipitate the protein. Reactions were then flash-frozen with liquid nitrogen and stored at -20 °C. Prior to HPLC analysis each reaction mixture was centrifuged at 13000 rpm for 20 min. HPLC analysis was carried out by HPLC method 3 as specified in Supplementary Table S5**.** The kinetic parameters of TreT *T.tenax* were also determined based on the variable acceptor substrate concentrations by GT-continued assay as described for TreT *P. horikoshii* under the similar conditions. The amount of UDP-Glc was fixed at 20 mM and Glc concentration varied

(0.5 mM, 1 mM, 2 mM, 4 mM and 10 mM). The reactions were incubated at 37 °C for 5 min and then enzyme (662 nM) added and reaction rates determined.

**OtsAB Enzyme Kinetic Assay**

OtsAB kinetic parameters were determined using HPLC method 3 as specified in Supplementary Table S5. We used variable UDP-Glc concentrations (2 mM, 4 mM, 8 mM, 16 mM and 32 mM) at a fixed G6P concentration of 80 mM. After adding the enzyme (1.23 µM), reactions were incubated for 5 min at 37 °C in a thermoshaker. Reactions were then quenched by adding acetonitrile, centrifuged (13000 rpm, 10 min) and analysed by HPLC. A standard calibration curve of UDP was also generated under the same reaction and analysis conditions. From the resulting UDP calibration curve, unknown concentrations from enzymatic reactions were determined and plotted against the concentrations of G6P substrate to obtain the $K_m$ and $V_{max}$ values.

**OtsB Enzyme Kinetics**

For OtsB$^{sf}$ enzyme kinetics we used $^1$H NMR spectroscopy employing the $^1$H anomeric resonances as primary signals that are well resolved and with low background. Initially, we analyzed standard samples of trehalose-6-phosphate (T6P) and trehalose. For kinetic assay, we incubated several concentrations of T6P (0.5 mM, 1.0 mM, 1.5 mM, 2.0 mM, 2.5 mM, 3.0 mM, 3.5 mM, 4.0 mM and 5.0 mM). Reaction was initiated by adding 10 µL of OtsB and incubated at 37 °C for 10 min. After 10 min, reactions were stopped by boiling at 95 °C for 5 min and then analysed by $^1$H NMR using $D_2O$ as a solvent. A standard calibration curve for trehalose was also generated and, using this calibration curve, concentrations of resulting trehalose from enzymatic reaction were obtained and analyzed through non-linear regression in GraphPad prism to give a Michaelis-Menten curve.

The activity of OtsB *E. coli* was also tested using commercially available Abcam Phosphate assay kit. By determining the amount of inorganic phosphate released from T6P by action of OtsB, enzymatic activity was to calculated. First, 10 µL of 10 mM phosphate standard was added to 900 µL of buffer (50 mM Tris, 5 mM $MgCl_2$, pH 7.5) to generate 100 µM standard phosphate solution. 0,10, 20,30, 40 and 50 µL of 100 µM standard phosphate solution were added to individual wells of a 96-well plate. The volume was adjusted to 200 µL with $dH_2O$ to generate phosphate standards. 30 µL of working reagent was added to all wells and was mixed well. The plate was incubated at 37 °C for 30 minutes. Absorbance was read at 650 nm. A calibration curve was thus generated. As a practical note, in earlier experiments, reaction aliquots were mixed with 30 µL to quench the reaction and then volume was made up to 200 µL; it was found that doing so leads to precipitation. Therefore, in later experiments, reaction aliquots were diluted in the well using the reaction buffer, followed by addition of working reagent. In this way, no precipitate was observed. Next, enzymatic reactions were conducted in a final volume of 200 µL containing the following components: 5 mM $MgCl_2$, 50 mM Tris buffer, pH 7.5, OtsB and T6P. 1.720 µM of enzyme was used in each case. T6P concentration was varied from 0.5 mM to 3 mM with an increment of 0.5 mM. Reaction aliquots were taken from the reaction at time, t = 0.5 min, 1 min, 2 min, 2.5 min, 3 min, 3.5 min, 4 min, 5 min, 6 min, 8 min and 10 minutes. The volume of reaction aliquots was taken in such a way that when diluted with reaction buffer, the maximum possible amount of phosphate remained within the linear range of the assay. 30 µL of working reagent was added after dilution. Samples were incubated at 37 °C for 30 min. Absorbance was read at 650 nm. Using the calibration curve, the rate of formation of Pi was calculated and plotted against time to give rate of reaction at a particular substrate concentration. Data for different substrate concentrations were analyzed using non-linear regression in GraphPad prism to give a Michaelis-Menten curve.

**Chemoenzymatic Reaction Optimisation and Scale-up**

During reaction optimisation we tested the effect of ATP concentrations to the conversion of FDG to FDG6P initially at 1 mg scale. We set-up four different reactions in 1.5 mL Eppendorf tubes as shown in Supplementary Table S7. Reactions were incubated at 37 °C for 1 h at 400 rpm in a thermoshaker and then analysed by $^{19}$F-NMR spectroscopy to observe the conversion.

To test the OtsA and OtsB enzymes needed for a reaction, we tested several concentrations of OtsA (6.65 µM, 13.3 µM, 18.2 µM and 36.6 µM) and OtsB (1.53 µM, 3.06 µM, 6.13 µM, 12.26 µM) in a 50 mg scale while maintaining all the other reaction parameters constant. Reactions were incubated at 37 °C at 400 rpm for 1 h and then analysed by $^{19}$F-NMR spectroscopy.

In order to further optimise the enzymatic reaction conditions using 3 enzyme synthesis method, we tested various substrate concentrations i.e. with fixed enzyme concentrations. In summary in a 50 mL falcon tube were added FDG (various FDG concentrations i.e. 6.25 mg, 12.5 mg, 25 mg and 50 mg), ATP with equivalent or slight excess molar concentration to FDG, hexokinase 5 mg (680 U), UDP-Glc (37 mg with final concentration of 30 mM), OtsA enzyme (2 mg, 18 µM), OtsB (0.4 mg, 6.2 µM) and KCl (100 mM). Reaction volume was made up to 2 mL by HEPES buffer solution (HEPES 50 mM, pH 7.5, 100 mM NaCl, $MgCl_2$ 10 mM). Reactions were incubated at 37 °C and samples were taken after 15 min, 30 min, 60 min, 120 min, 200 min and 690 min, diluted with $D_2O$ and analysed by $^{19}$F-NMR spectroscopy. 50 mg reaction was further optimised with additional UDP-Glc (168 mg, 138 mM) and analysed by $^{19}$F-NMR.

**Chemoenzymatic Synthesis of FDT**

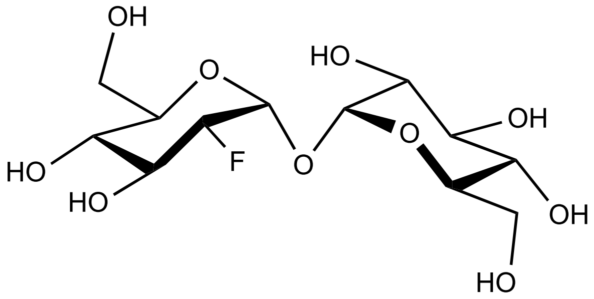

In a 50 mL sterile falcon tube were added; FDG (100 mg, 367 mM), ATP (320 mg, 368 mM) and $MgCl_2$ (5 mg). The contents were dissolved in HEPES buffer solution containing 100 mM NaCl (100 mM, 1 mL, pH 7.5). The pH was then adjusted to 8.0 with NaOH. Hexokinase (0.6 mL) was added to the reaction mixture to initiate the reaction. The reaction mixture was incubated at 30 °C, 50 rpm for 1-2 hours. Phosphorylation of FDG to FDG6P was monitored by TLC and $^{19}$F-NMR spectroscopy. Full conversion to FDG6P was observed within 2 hours for each batch synthesized.

Within the same pot, UDP-Glc (200 mg, 110 mM), KCl (100 mM) and OtsA enzyme (26 µM) was added. The total reaction was then adjusted to 3 mL. The reaction mixture was incubated to similar conditions as above for 24 h to allow appropriate conversion to FDT6P. The conversion of FDG6P to FDT6P was monitored by $^{19}$F-NMR spectroscopy. The average maximum conversion after 24 h was between 65-70 %. After 24 h, in the same reaction mixture OtsB enzyme (6.23 µM) was added. The pH was adjusted to 8.0 and reaction mixture was incubated at 37 °C, 50 rpm. Reaction progress was monitored by TLC and $^{19}$F-NMR. The reaction was usually complete within 2 hours.

**Purification and Characterisation of FDT**

The enzymes from crude reaction mixture were precipitated by either heating the mixture at 95 °C for 10 min or adding 2 mL EtOH and isolated by centrifugation (Vivaspin, 10,000 MWCO) and/or filtration. The resulting filtrate was lyophilized and the crude mixture was further precipitated with MeOH/MeCN 90:10 solution and then purified by flash column chromatography. For purification normal phase silica was used as stationary phase and mixture of Ethyl acetate: Iso-propanol and water (48%/48%/4%) were used as mobile phase. FDT was separated from residual FDG6P and residual FDG. The fractions containing FDT were pooled, concentrated and then lyophilised to dryness. The Lyophilisation step was repeated for a further 2-3 times to get rid of any residual water. The freeze dried compound was then characterised by proton ($^{1}H$ NMR), carbon ($^{13}C$ NMR), fluorine ($^{19}F$ NMR) and ESI-MS. $[\alpha]_D^{25}$ + 128.9 (c 1.00, $H_2O$). $^{1}H$ NMR (500 MHz, $D_2O$) δ 5.33 (d, $J$ = 3.9 Hz, 1H, H-1), 5.11 (d, $J$ = 3.7 Hz, 1H, H-1′), 4.40 (ddd, $J$ = 49.0, 9.6, 3.9 Hz, 1H, H-2), 4.02 (dt, $J$ = 13.2, 9.4 Hz, 1H, H-3), 3.79 – 3.62 (m, 7H, H-5, H-6a, H-6b, H-3′, H-5′, H-6a′, H-6b′), 3.56 (dd, $J$ = 9.9, 3.7 Hz, 1H, H-2′), 3.41(t, $J$ = 9.7 Hz, 1H, H-4) , 3.35 (t, $J$ = 9.4 Hz, 1H, H-4′) ppm. $^{13}C$ NMR (125 MHz, $D_2O$) δ 94.0 (C-1′), 91.2 (d, $J_{F,C-1}$ = 21.8 Hz, C-1), 89.5 (d, $J_{F,C-2}$ = 186.6 Hz, C-2), 72.6 (C-3′), 72.2 (C-5 and C-5′), 71.4 (d, $J_{F,C-3}$ = 15.8Hz, C-3) 70.9 (C-2′), 69.6(C-4′), 69.1 (d, $J$ = 8.0 Hz, C-4), 60.5 (C-6 or C-6′), 60.3 (C-6 or C-6′) ppm. $^{19}F$ NMR (470 MHz, $D_2O$) δ – 201.18 ppm. IR (neat) ν 3264 (OH), 1647, 1409, 1150, 1103, 990, 948, 841, 802, 705 $cm^{-1}$. ESI-MS: ESI-MS calculated for $C_{12}H_{21}F_1O_{10}Na_1$ [M + Na] 367.1011, found 367.1011.

**Stability Testing of FDT Solution in HEPES Buffer**

In order to establish the storage conditions of the final formulation of an injection, small scale stability testing of the final formulation was carried out. In summary, FDT sample was dissolved in HEPES Buffer solution (50 mM, pH 7.5) containing 7.5 mM $MgCl_2$. One portion of the dissolved solution was stored at room temperature (20 – 25 °C) and the other portion was stored between 2-8 °C. Samples were taken at day 0, 3, 5, 7 and 10 and analysed by $^{19}F$

NMR spectroscopy in triplicate. The means of the obtained integrals are then plotted against the day of testing. In addition, spectras were observed for any additional peaks as well.

#### Stability Testing of FDT Solid

In order to ensure that the compound will be stable throughout the testing period, on-going stability testing of the FDT sample from solid was also investigated over nine months' period. Material for testing was stored at - 20 °C and periodic testing was carried out by dissolving ~ 8 mg of sample in $D_2O$ and then subsequently analysed by $^1$H NMR spectroscopy.

#### Determination of Purity by $^1$H NMR

Purity of the final compound was determined by $^1$H NMR spectroscopy by determining relative concentration of the substance as compared to impurities. Known quantity of FDT was dissolved in $D_2O$ and quantified by $^1$H NMR spectroscopy. Maleic acid was used as an internal standard.

#### $^{23}$Na-NMR Quantification

In order to determine the amount of salt (NaCl) in the FDT compound we used $^{23}$Na-NMR in AVX500 instrument. We generated the calibration curve of NaCl solution of known concentration and then run the FDT samples using the same parameters as NaCl standards. The amount of Na was calculated from the standard calibration curve as an unknown and then from molar ratio of Na:FDT the amount of salt then determined.

**[$^{18}$F]FDT Synthesis using TreT *P. horikoshii***

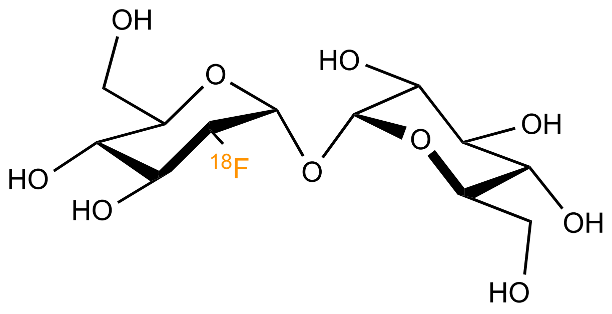

In a 2 mL Eppendorf tube were added UDP-Glc (30 µL of a 1M aq solution), TreT enzyme (0.7 mg). Reaction was started by adding [$^{18}$F]FDG (500 µL, 14.1 mCi/mL ). Reaction volume was made up to 1 mL by adding HEPES buffer solution (100 mM HEPES, 10 mM $MgCl_2$, pH 7.5). The reaction was incubated at 37 °C in a thermoshaker and monitored by radio-HPLC. Once [$^{18}$F]FDG fully converted to [$^{18}$F]FDT, the reaction mixture was purified on a Luna-$NH_2$ matrix as described in the Methods section.

**[$^{18}$F]FDT Synthesis using 3 Enzymes in one pot**

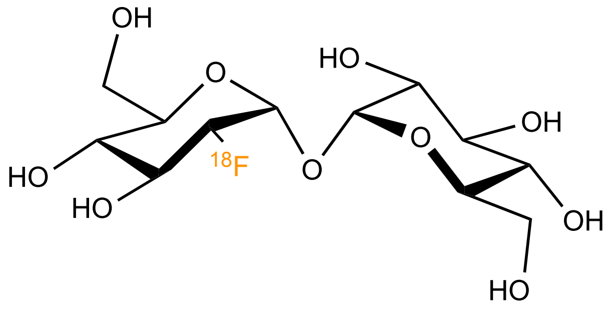

In a 2 mL Eppendorf tube were added UDP-Glc (30 µL of a 1M aq solution), ATP (10 mM), hexokinase (5 mg, 640 U), OtsA (0.7 mg, 12.3 µM), OtsB (0.4 mg, 12.5 µM). [$^{18}$F]FDG (500 µL, 15.6 mCi) was added to the reaction mixture to start the reaction. Final volume of reaction was adjusted to 1 mL by adding HEPES buffer solution and incubated at 37 °C in a thermoshaker. Reaction progress was monitored by radio-HPLC and stopped after 60 min when fully converted to the product and purified by Luna-$NH_2$ cartrdige method.

**[$^{18}$F]FDT Synthesis using 2 Enzymes in one pot**

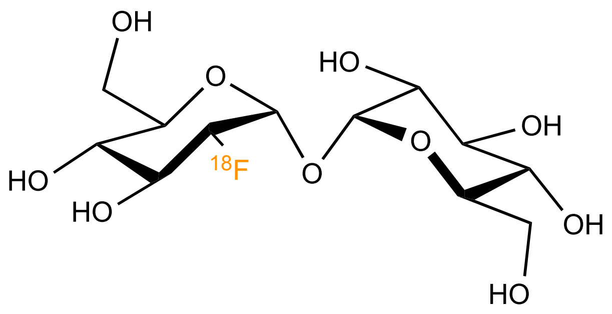

In a 2 mL Eppendorf tube were added UDP-Glc (30 µL of a 1M aq solution), ATP (10 mM), OtsAB fusion enzyme (0.1 mg. 1.3 µM), hexokinase ( 2 mg, 256 U). The reaction was started by the addition of [$^{18}$F]FDG (250 µL, 3.5 mCi). The reaction volume was adjusted to 1 mL by adding HEPES buffer solution (HEPES 100 mM, $MgCl_2$ 10 mM, NaCl 100 mM, pH 7.5). The reaction was started by incubating at 37 °C and monitored by radio-HPLC. The reaction did not work under the conditions described above even after 90 min of incubation.

In an another attempt, higher enzyme concentrations were added into the reaction mixture and incubated for 2 h at 37 $^{o}$C but there was no conversion at all.

**Automation of [$^{18}$F]FDT Synthesis**

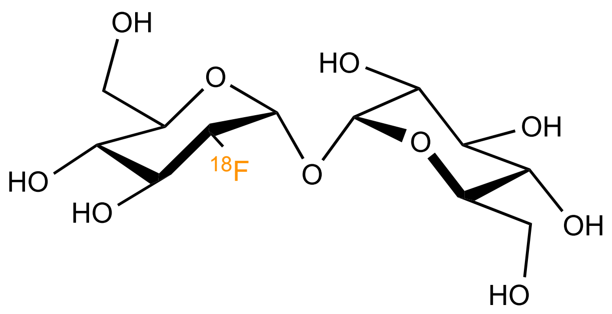

[$^{18}$F]FDG (1480- 3700 MBq; 40-100 mCi in 0.8 -1.8 mL) was transferred under vacuum to the Reactor 1 containing 100 µL 1 M HEPES buffer, pH 7.6, 20 µL 1 M $MgCl_2$, 20 µL 1 M

ATP, 60 µL 1 M UDP-Glc, ~50 µL OtsA (1 mg), ~50 µL OtsB (1 mg), 20 µL hexokinase (5 mg). The reaction mixture was incubated for 30 min at 45 °C and ethanol was added (3 - 6 mL). The reaction mixture was passed through the filter (5 µm) and a stack of three $NH_2$-cartridges and collected in Reactor 2. An additional 1 mL 75% ethanol in water was added to rinse the Reactor 1 and transferred into the Reactor 2. The combined solution was concentrated under nitrogen at 60 °C for 10 min. 2 mL saline was added to the Reactor 2. The final [$^{18}$F] FDT solution was transferred to the product vial through a sterile filter (0.22 µm). The quality of the product was determined by analytical radio-HPLC (Condition: 4.6 x 250 mm, 2.7 µm AdvanceBio Glycan Mapping Column; solvent A = 50 mM ammonium formate, pH 4.5, solvent B = acetonitrile; Flow rate = 0.5 mL/min;gradient 0-15 min 68-62% B; 15-20 min 62-68% B). Identity of the compound was confirmed by LC-MS.

**Chemoenzymatic synthesis of [$^{18}$F]FDT for Animal Studies**

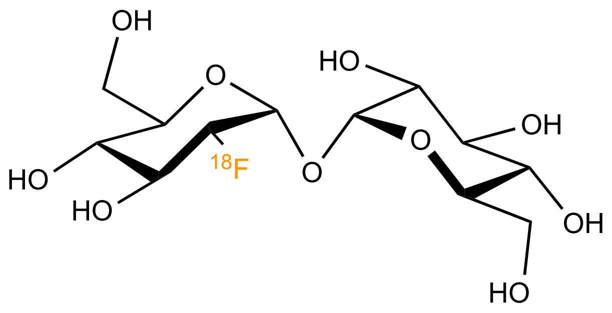

For the animal studies, [$^{18}$F]FDG (20-30 mCi in 0.8 -1 mL) was injected to the reaction mixture containing 100 µL 1 M HEPES buffer, pH 7.6, 20 µL 1 M $MgCl_2$, 20 µL 1 M ATP, 60 µL 1 M UDP-Glc, ~50 µL OtsA (1 mg), ~50 µL OtsB (1 mg), 20 µL hexokinase (5 mg). The reaction mixture was incubated at 37 °C for 30 min. After 30 min, the mixture was diluted with absolute ethanol (4 mL) and passed through a 5 µm syringe filter. The eluent was passed slowly through an amine Sep-Pak SPE cartridge at a flow rate of 1-2 drops per second.

The eluent was then concentrated in vacuo. The resulting solution was filter-sterilized into a sterile vial for delivery. Identity of the compound was confirmed by LC-MS.

**Specific Activity Analysis of [$^{18}$F]FDT**

The radioactivity of the final products, [$^{18}$F]FDT, was obtained using a radiation detector. To quantify the mass of the decayed [$^{18}$F]FDT in the form of [$^{18}$O] trehalose, a modified version of the trehalose quantification enzymatic assay was performed*(46)* . A trehalose standard calibration curve (linear fit, $R^2$ = 0.9984) was generated using known concentrations of trehalose. The 'cold equivalent' masses of [$^{18}$F]FDT from syntheses were calculated based on this calibration curve. Then, the specific activity of the final product was calculated, following the definition, radioactivity at the end of the synthesis / unit mass of compound.

**Chemical synthesis of [$^{19}$F]-3-FDG**

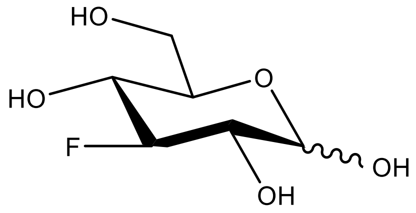

In a 100 mL flask was dissolved 0.50 g of diacetone allofuranose (1.92 mmol, 1 eq) and 0.50g of *N*,*N*-diaminopyridine (4.1 mmol, 2.1 eq) in 20 mL of dry dichloromethane. The mixture was cooled down to -20°C using an acetone/dry ice bath then 0.5 mL of diethylaminosulfur trifluoride

(DAST, 3.8 mmol, 2.0 eq) was added slowly, then the solution was allowed to return slowly to room temperature while stirring overnight. TLC (95:5 CH2Cl2:MeOH) showed a single compound ($R_f$ 0.77). The mixture was cooled to 0 °C and 5mL of methanol was added slowly. The mixture was stirred 1h and evaporated to dryness to give a yellow oil that was purified by silica column chromatography (49:1 $CH_2Cl_2$:EtOAc then 24:1 $CH_2Cl_2$:EtOAc) to give 416 mg of a colourless oil, that was used in the next step.

In a 50mL flask was dissolved 416mg of diacetone 3-FDG (1.58 mmol) in 20mL of water and 4 mL of ethanol. Ca. 6 mL of activated DOWEX 50W-H8 was added and the mixture was stirred for 92 hours. The mixture was filtered, and the liquid was evaporated and dried under high vacuum to give 240.9 mg of crude compound. This crude compound was dissolved in a mixture 7:2:1 ethyl acetate:ethanol:water and filtered through a short plug of silica, then the silica

was washed with 200mL of the same mixture. The solution was evaporated and dried under high vacuum to give 128 mg of a colorless oil in 44% overall yield.

Analysis: $^{1}$H NMR (400 MHz, $D_2O$): 5.14 (1H, app t, 4.0 Hz, H-1 α), 4.56 (1H, d, 8.0 Hz, H-1 β), 4.49 (1H, dt, $J_{H\text{-}F}$ 52 Hz, 8 Hz H-3) 4.31(1H, dt, $J_{H\text{-}F}$ 52 Hz, 8 Hz, H-3), 3.80-3.40 (10H, m, H-2, H-4, H-5,H-6, H-6'); $^{13}$C NMR (100.6 MHz, $D_2O$): 96.8 (d, $J_{C\text{-}F}$ 181 Hz C-3), 95.3 (d, $J_{C\text{-}F}$ 179 Hz C-3'), 95.5(d, $J_{C\text{-}F}$ 12 Hz C-1 beta), 92.5 (d, $J_{C\text{-}F}$ 10 Hz C-1 α), 74.9 (d, $J_{C\text{-}F}$ 9 Hz ), 72.9 (d, $J_{C\text{-}F}$ 16 Hz), 71.1(d, $J_{C\text{-}F}$ 7 Hz), 70.3 (d, $J_{C\text{-}F}$ 17 Hz), 68.3 (C-5), 68.1 (C-5), 60.6(C-6), 60.5(C-6); $^{19}$F NMR(376.5 MHz, $D_2O$): β -195.2, dt, $J_{H\text{-}3\text{-}F}$ 52.7 Hz, $J_{H\text{-}4\text{-}F}$ =$J_{H\text{-}2\text{-}F}$ =15.0 Hz; α -200.1, dtd, $J_{H\text{-}3\text{-}F}$ 52.7 Hz, $J_{H\text{-}4\text{-}F}$, $J_{H\text{-}2\text{-}F}$ 15.0 Hz, 4.0 Hz; MS ($ESI^-$) $[M+Cl]^-$ found 217.05 and 219.05, require 217.0279 and 219.0250.

**1,2:5,6-di-*O*-isopropylidene-3-*O*-triflyl-α-D-allofuranose**

260 mg of 1,2:5,6-di-*O*-isopropylidene D-allofuranose (1 mmol, 1eq) and 308 mg of 2,6-di-tertbutyl-4-methylpyridine (1.5 mmol, 1.5 eq) were dissolved in 10 mL of dichloromethane. 220 µL of trifluoromethanesulfonic anyhydride (1.3 mmol, 1.3 eq) were added slowly at room temperature and the mixture was stirred overnight at room temperature. TLC showed

completion of the reaction, and the mixture was evaporated to dryness under vacuum at less than +30°C. The

residue was purified by silica column chromatography (95:5 $CH_2Cl_2$:EtOAc) to give 309.7 mg of as a colorless oil (0.788 mmol, 79%).

Analysis: TLC $R_f$=0.85 (80:20 $CH_2Cl_2$:EtOAc); MS (ESI+) found 415.05, [M+Na+] requires 415.0645; $^1$H NMR (400 MHz $CDCl_3$) δ ppm 5.84 (1H, d, $J_{1,2}$=4.0Hz, H-1), 4.91 (1H,

app t, 6.2 Hz, H-3), 4.78 ( 1H, app t, 4.0 Hz), 4.19 (2H, m, H-4 and H-5), 4.12 (1H, dd, $J_{6a\text{-}6b}$=8.8Hz, J6a-5=6.2 Hz, H-6a), 3.92 (1H, dd, $J_{6b,6a}$=8.8Hz, $J_{6b,5}$=4.8 Hz, H-6b), 1.59, 1.45, 1.39,

1.35 (12H, 4s, 4 x $CH_3$ isopropylidene). $^{13}$C NMR (101 MHz $CDCl_3$) δ ppm 114.4, 110.3 ( isopropylidene acetal), 104.1 (C-1), 82.9 (C-3), 77.8, 77.7 (C-2 and C-4 or C-5), 75.1 (C-5 or C-

4), 66.3 (C-6), 26.9, 26.5, 26.2, 24.8 (4 x $CH_3$ isopropylidene), $CF_3$ OTf weak. $^{19}$F NMR (376.5 MHz, $CDCl_3$) δ ppm -74.9, s, $CF_3$ OTf.

**Attempted Chemoenzymatic Synthesis of [$^{18}$F]-3-FDT**

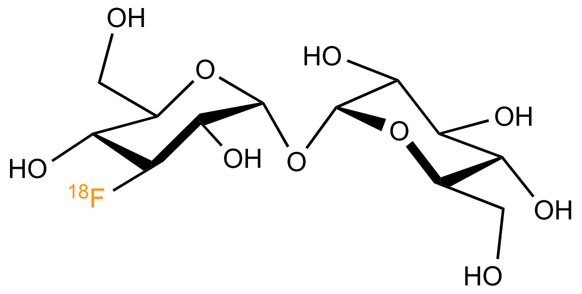

To a 100 μL solution of scintomics dried K[$^{18}$F-]/K222 solution (activity 193.2 MBq at 12:40, was added 200 μL solution of 3-OTf diisopropylidene allofuranose (18.4 mg in acetonitrile). The mixture was heated at 110 ˚C for 10 min, after which it was cooled down to room

temperature. 200 μL of 5 N HCl solution was added into the reaction mixture and the solution was heated at 110 °C for further 10 min, cooled to room temperature and monitored by TLC for the formation of [$^{18}$F]-3-FDG. 92 MBq of [$^{18}$F]-3-FDG in 0.5 M phosphate buffer solution (phosphate buffer pH 7.5) from the above reaction mixture was used in an attempted enzymatic one pot synthesis. Following reagents were added : 12 μL OF 1M ATP, 45 μL of 1 M UDP-Glc, 12 μL 1M $MgCl_2$, 100 μL OtsA (23 mg/mL) and 35 μL OtsB. The reaction vial was incubated at 30 °C and monitored by radio-TLC over 4h.

**Chemoenzymatic Synthesis of [$^{19}$F]-3-FDT**

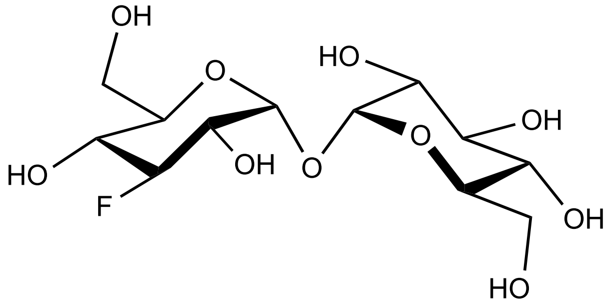

A one-pot reaction mixture containing 2.5 mM 3FGlc, 8.3 mM ATP, 33 mM UDP-glucose, 1.3 mg/mL OtsA and 3.4 mg/mL OtsB was prepared in pH 7.25, 50 mM Hepes buffer. Hexokinase (5.0 mg/mL final concentration) was added to initiate the reaction. Reaction mixture was incubated at 30 °C and monitored by ESI-MS at every 10 minute interval.

Alternatively, aliquots were withdrawn at every 30 minutes. Withdrawn aliquots were frozen on liquid nitrogen and boiled for 1 minute. Proteins were filtered off using a Vivaspin (Sartorius, 10K NMWCO) and each aliquot was analyzed by HPLC (Dionex UltiMate 3000) with the column, Phenomenex Luna $NH_2$ (4.6 × 250 μm). The column was eluted with isocratic 7/3 acetonitrile/water. The maximum concentration of OtsB that could be used was 3.4 mg/mL. Above this concentration, OtsB precipitated while the reaction did not proceed to a meaningful extent at lower than this concentration of OtsB. By ESI/MS, it was found that the reaction did not go completion in 2 hours, after which precipitation formed. Based on peak intensity it was estimated that a mixture 7:1 3FTre:3FGlc was formed. HPLC analysis also revealed poor separation.

**Radiochemical synthesis of [$^{18}$F]-6-FDT**

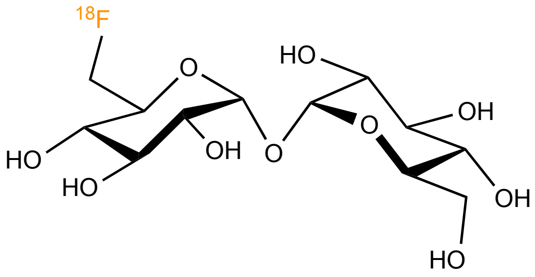

To a 13 x 100 TT, were added 100 µL K222 solution and 60 µL $K_2CO_3$, and 20 µL $^{18}F^-$ (3.20 mCi). The reaction contents were evaporated under stream of argon in a 105 °C heat block. The reaction mixture was then azeotroped 3 times with 200 µL of $CH_3CN$. Triflate precursor, trehalose heptaacetate triflate was dissolved in 300 µL of $CH_3CN$ and added into the reaction mixture and incubated at +55 °C for 15 min. After 15 min, 30 µL of 5 M NaOMe (150 µmol) was added and the reaction was allowed to stand for 5 min. After 37 min, reaction was monitored by analytical HPLC (Luna $NH_2$ 4.6 x 250 mm; isocratic elution of 80% $CH_3CN$ and 20% $H_2O$; flow rate 1 mL/min). 6-[$^{18}$F]-Tre eluted at 8.5 min. The reaction contents were then purified by preparative HPLC (Luna NH2 10 x 150 mm; isocratic elution of 80% $CH_3CN$ and 20% $H_2O$; flow rate 4 mL/min). The desired compound was obtained in 32% yield (± 4%, non-decay corrected, n=2, from preparative HPLC) over 50-55 minutes total reaction time.

**Radiochemical synthesis of 4-epi-[$^{18}$F]-FDT**

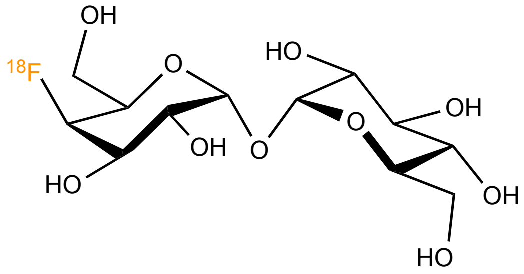

To a 13 x 100 TT, was combined aqueous potassium carbonate (0.1 M, 60 mL, 12 mmol, 2.4 eq), K222 (4.5 mg, 12 mmol, 2.4 eq) as a solution in MeCN (100 mL, 120 mM), and [$^{18}$F]-fluoride (120 mL, 100 mCi). The mixture was azeotroped at 110 °C, washing with MeCN (3 x 200 mL). To this dried mixture was added substrate (5 mg, 6.5 mmol, 1 eq) in MeCN (200 mL, 25 mM), and the reaction was heated at 120 °C for 15 min. the resulting brown solution was diluted with saline (154 mM NaCl, 15 mL) and passed slowly through a $C_{18}$ Sep-Pak. The Sep-Pak was then washed with water (1 x 10 mL), then treated with NaOH (1 M, 200 mL). The NaOH was dispensed onto the Sep-Pak, which was allowed to stand for 5 min at room temperature. The deprotected product was then eluted with water (3 mL), acidified with HCl (1 M, 150 mL) and buffered with saline-sodium citrate (10X, 300 mL). The neutralized material was purified by passage through an Alumina Sep-Pak Plus, followed by a $C_{18}$ SPE). Material was analyzed for radiochemical purity by HPLC and TLC. HPLC was performed on an Agilent 1100 on a Phenomenex Luna –$NH_2$ 4.6 x 250 mm column using an isocratic flow rate of 80:20 MeCN:$H_2O$ at 1 mL/min. TLC analysis was performed using multichannel silica plates with a cellulose preabsorbing zone developed in a mixture of 1:4:4 water:isopropanol:ethyl acetate.

Syntheses were performed in a 21-150 mCi scale yielding 9-66 mCi of epi-4-F-Tre, with an average radiochemical yield (n=10) of 33.7 (+/- 8%, non-decay corrected) and an average synthesis time of 57 min.

### Supplementary Figures

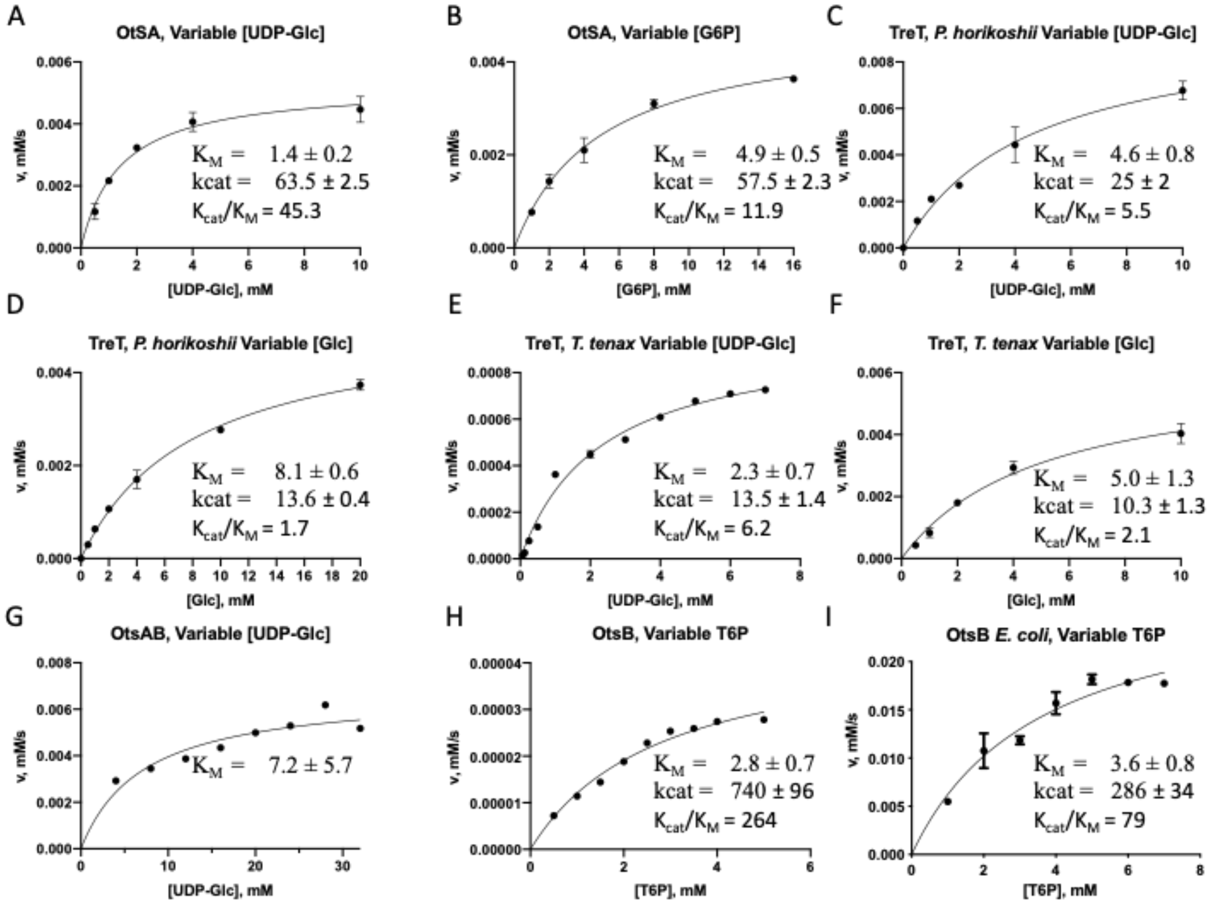

**Supplementary Figure S1. Kinetic parameters donor and acceptor substrates of OtsA, TreT, OtsAB fusion enzyme and OtsB enzyme.**

**A**) Michaelis-Menten plot showing the activity of OtsA as a function of UDP-Glc with a fixed Glc-6P concentration, **B**) Michaelis-Menten plot showing the activity of OtsA as a function of Glc-6P with a fixed UDPG concentrations. **C**) Michaelis-Menten plot showing the activity of TreT *P. horikoshii* as a function of UDP-Glc with a fixed Glc concentration. **D**) Michaelis-Menten plot showing the activity of TreT *P. horikoshi*i as a function of Glc with a fixed UDP-Glc concentrations. **E**) Michaelis-Menten plot showing the activity of TreT *T. tenax* as a function of UDP-Glc with a fixed Glc concentration, **F**). Michaelis-Menten plot showing the activity of TreT *T. tenax* as a function of Glc with a fixed UDP-Glc concentration. **G**). Michaelis-Menten plot showing the activity of OtsAB fusion as a function of UDP-Glc with a fixed G6P concentration. **H**) Michaelis-Menten plot showing the activity of OtsB$^{sf}$ as a function of T6P. **I**) Michaelis-Menten plot showing the activity of OtsB as a function of T6P. Units for kinetic parameters are $s^{-1}$ for $k_{cat}$ and mM for $K_M$.

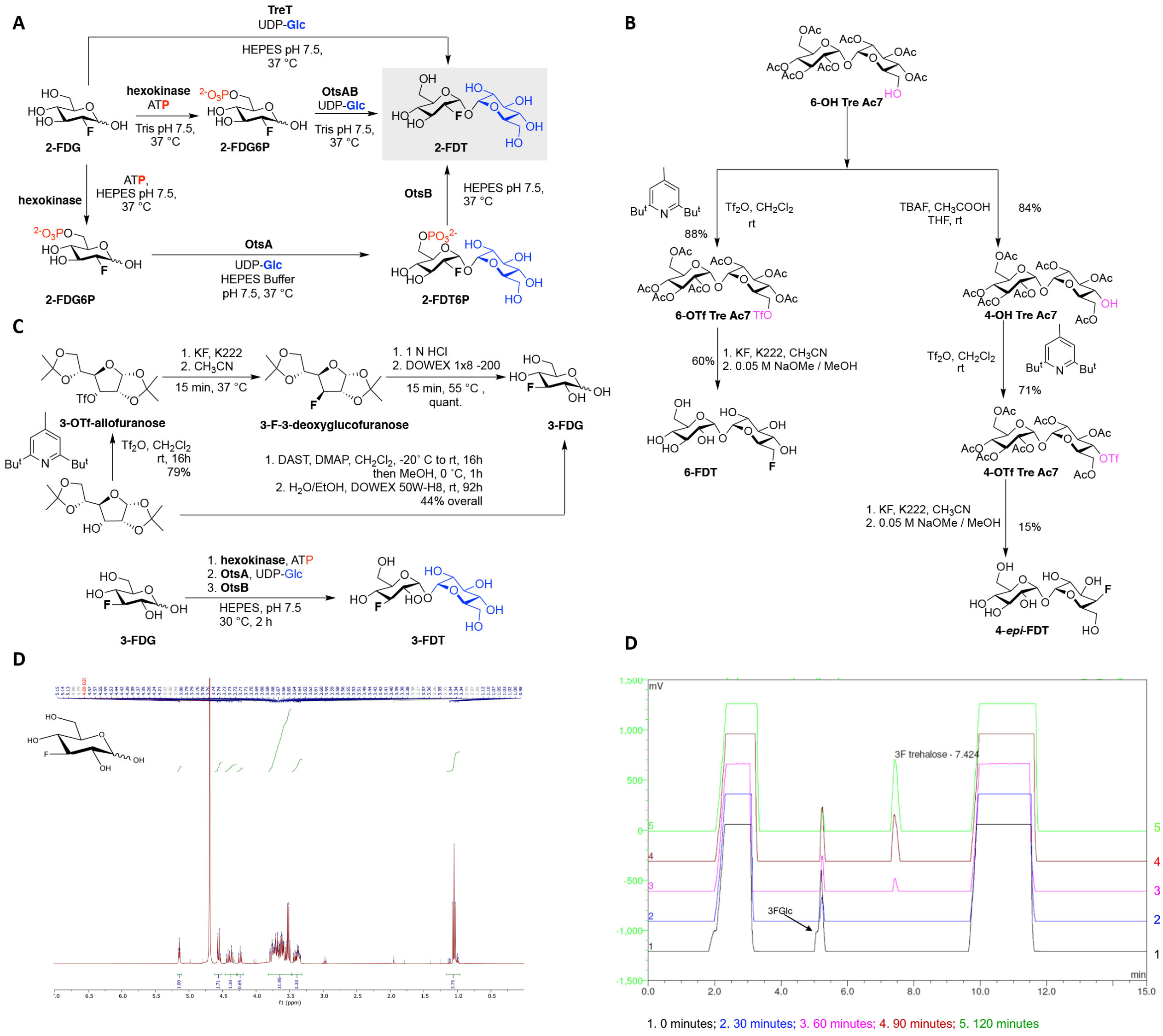

**Supplementary Figure S2. Exploration of 'Cold' Synthetic Routes to FDT variants.**

The 'cold' synthesis of 4 different $^{19}$F-containing FDT variants (termed here 2-FDT – described also as FDT in the main text –, 3-FDT, 4-*epi*-FDT and 6-FDT) was tested and compared by using both chemical and chemoenzymatic approaches. (**A**) For 'cold' 2-FDT synthesis, we tested 3 different chemoenzymatic approaches, see the main text for further details. (**B**) Cold syntheses of both 4-epi-FDT and 6-FDT were tested from the corresponding mono-hydroxy precursors; corresponding *O*-triflate derivatives were synthesised in high yield (88% for 6-OTf-Tre(Ac)$_7$ and 84% for 4-OTf-Tre(Ac)$_7$). Subsequently the substitution of these triflate precursors yielded corresponding fluorinated sugars (6-FDT 60%, 4-epi-FDT

15%). These formed the basis for synthetic routes to corresponding ‘hot’ $^{18}$F variants (see Supplementary Figure S3). (**C**) ‘Cold’ synthesis of 3-FDT was attempted in two stages. In the first stage, precursor 3-fluoro-3-deoxy-D-glucose (3-FDG) was synthesised directly using DAST (to provide a reference) and also from the corresponding triflate precursor (to mimic putative radiosynthesis), as confirmed by $^{1}$H NMR of reaction product (**D**). This 3-FDG was then used in attempted, one-pot chemoenzymatic syntheses of 3-FDT (2.5 mM scale in HEPES buffer solution, 50 mM HEPES buffer, pH 7.25, 30 °C with UDP-Glc(33 mM), hexokinase (5 mg/mL), ATP (8.3 mM), OtsA (1.3 mg/mL) and OtsB (3.4 mg/mL)). Progress of reaction was monitored by HPLC (HPLC chromatograms of this reaction are shown in (**E**)) but did not reveal effective conversion/separation in a manner concomitant with automated radiosynthesis. These were consistent with poor results from attempted synthesis of 3-[$^{18}$F]FDT (see below).

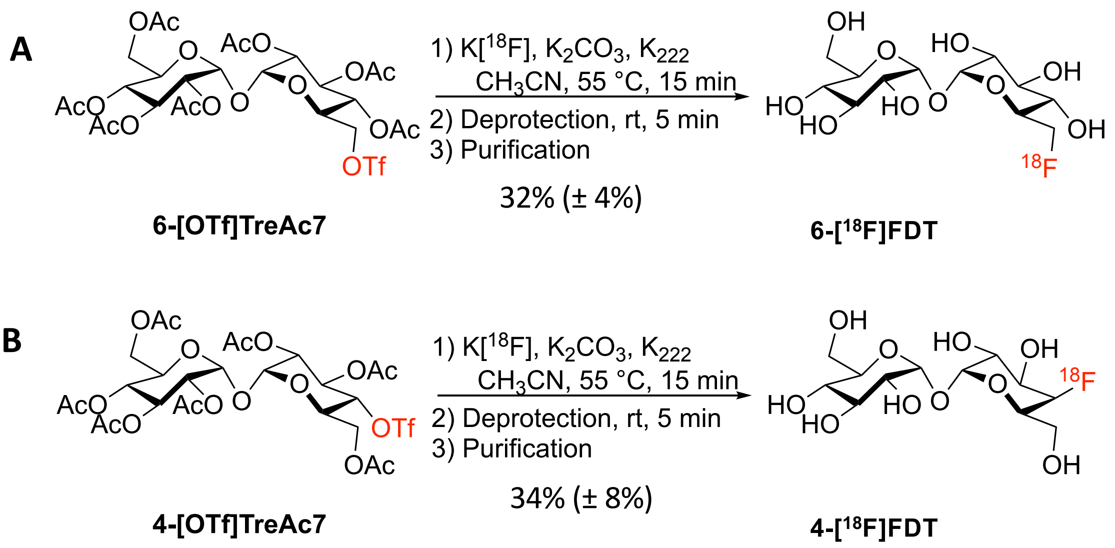

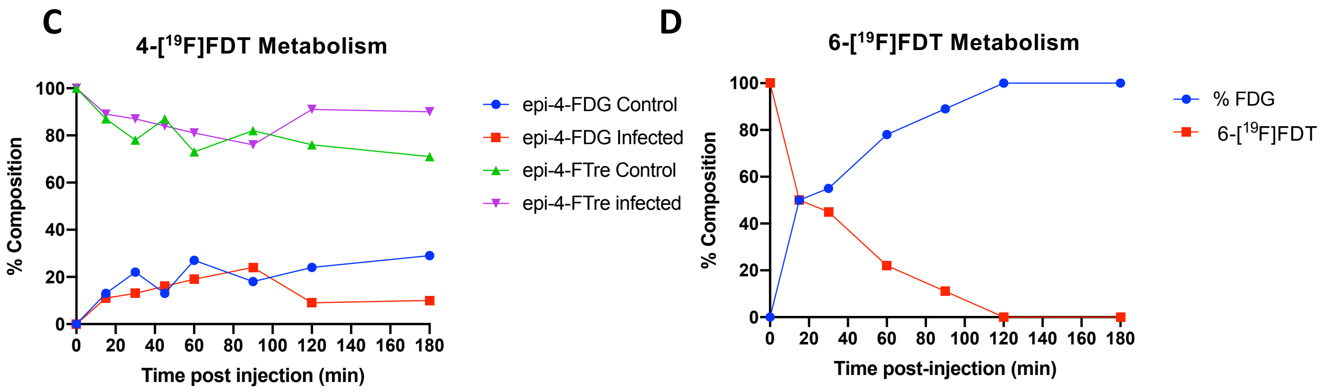

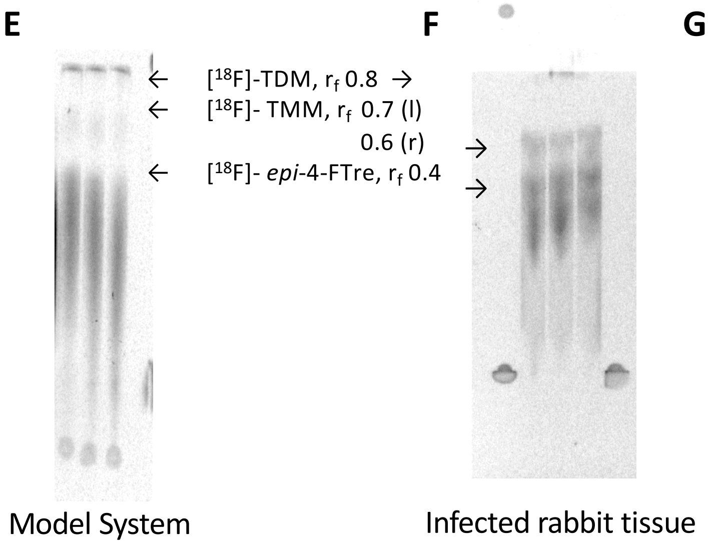

G

| Time (min) | % FDG | % 2-FDT |
|---|---|---|
| 50 | 1.5 | 98.5 |
| 115 | 5.4 | 94.6 |
| urine (120) | 4.2 | 95.8 |

**Supplementary Figure S3. Evaluation of Other [$^{18}$F]FDT Analogs – Synthetic strategies, in vivo Metabolism and Biodistribution/Biostability.**

(**A**) Synthetic route to 6-[$^{18}$F]FDT. **(B)** Synthetic route to 4-*epi*-[$^{18}$F]FDT. **(C)** Concentration-time plot showing that 6-[$^{18}$F]FDT was metabolized rapidly within the typical timeframe of a clinical imaging session with a [$^{18}$F] probe. **(D)** Concentration-time plot showing that 4-*epi*-[$^{18}$F]FDT was metabolized significantly within the typical timeframe of a clinical imaging session with an [$^{18}$F] probe. The all but final timepoints were from blood samples taken from the injected subject. The final timepoint represents the [$^{18}$F]F-labeled constituents in the urine drawn from the bladder at the indicated time. **(E)** [4-*epi*-[$^{18}$F]FDT was incorporated into TMM and TDM in *ex vivo* mouse lung spiked with *M. tuberculosis* after 60 minutes incubation. **(F)** $^{18}$F-trehalose was incorporated in vivo into TMM and TDM. Lipids were extracted from cavity caseum collected from a *M. tuberculosis* infected rabbit 90 minutes after I.V. injection of the probe. The lipid extract was separated by TLC and the radioactivity detected by autoradiography. **(G)** Tabular comparison illustrating that when [$^{18}$F] was incorporated at the 2 position in 2-[$^{18}$F]FDT / [$^{18}$F]FDT was less subject to metabolism than when placed into the 6 or 4 position.

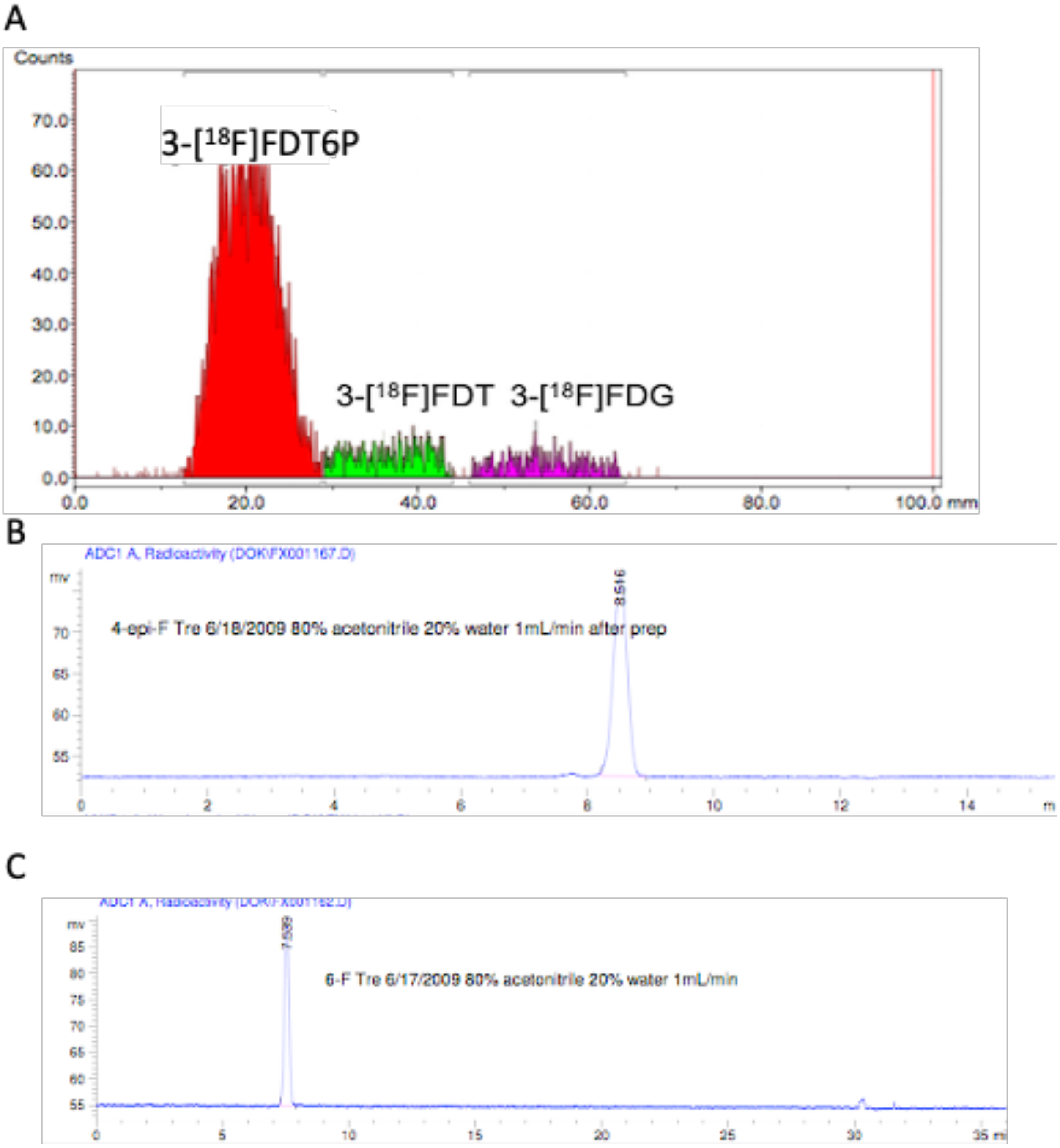

**Supplementary Figure S4. Chromatograms for the Radiosynthesews of [18F]FDT Analogs.**

**A)** Radio-TLC chromatogram of 3-[$^{18}$F]FDT synthesis from 3-[$^{18}$F]FDG. Only 10% conversion to 3-[$^{18}$F]FDT observed over 60 min ($R_f$ = 0.31; region 2), whereas 83% of the intermediate product 3-[$^{18}$F]FDT6P was observed at the start ($R_f$ = 0.0; region 1) and around 6% of the starting material i.e. 3-[$^{18}$F]FDG observed ($R_f$ =0.54; region 3). **B**) Analytical radio

HPLC chromatogram of 4-epi-[$^{18}$F]FDT synthesis, showing full conversion to the desired product with 9% yield. **C**) Analytical radio HPLC chromatogram of 6-[$^{18}$F]FDT with 25 ± 3% overall yield (n=2)

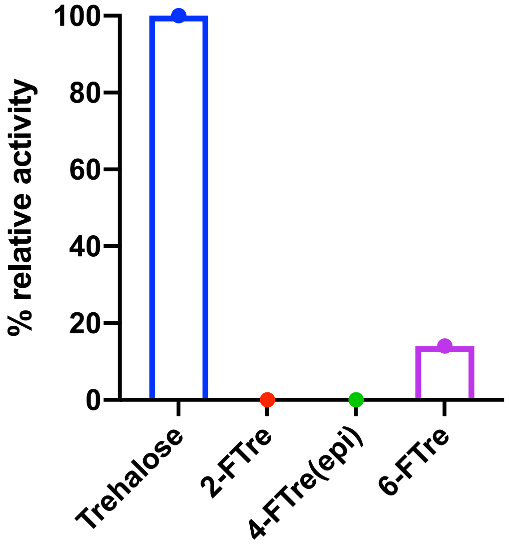

**Supplementary Figure S5. Trehalase Assays.**

Substrates were prepared to a final concentration of 10 mM in 0.05 M citric acid, pH 5.7 including trehalase stock solution (0.3 units/mL). The reaction mixture was incubated at 37 °C for 60 minutes, and then quenched with an equal volume of 0.1 M Tris, pH 7.5. Released glucose was measured with glucose assay kit (see Methods).

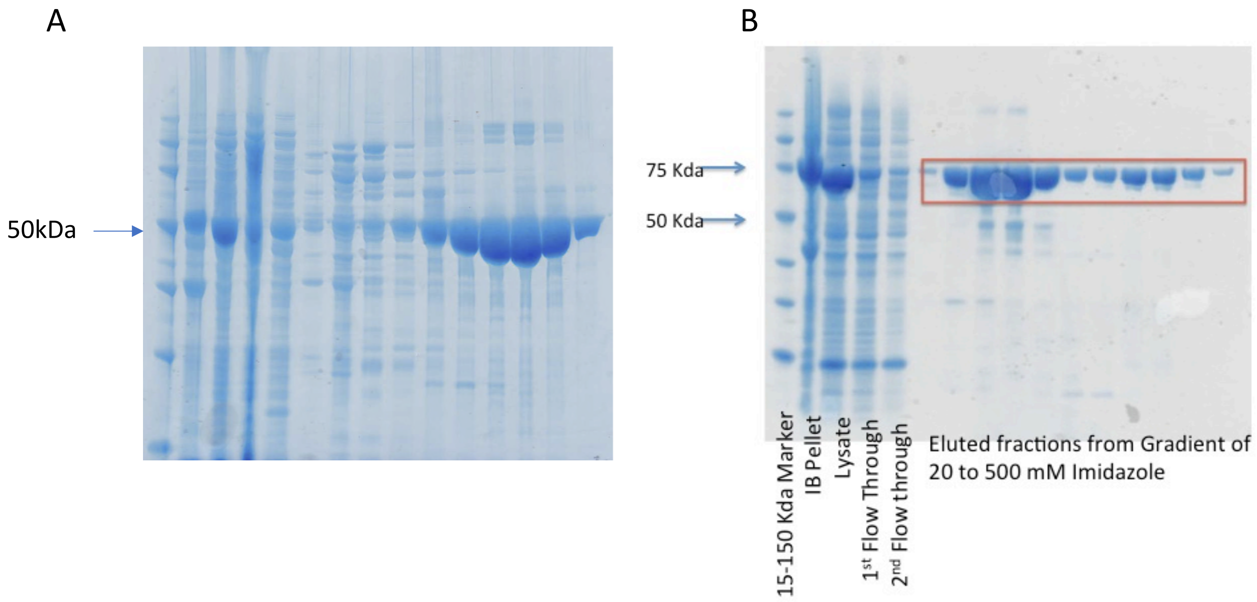

**Supplementary Figure S6. OtsA and OtsB *E. coli* Expression, Purification and Characterisation.**

**A)** 10% Bis-Tris SDS PAGE Analysis of OtsA expression run in MOPS buffer at 200 V for 50 minutes. Analytes: 15-150 KDa Protein marker, IB pellet, Lysate, 1st two flow through, 2nd-11th eluted fractions. **B)** SDS-PAGE Analysis of His-MBP-OtsB protein expression.

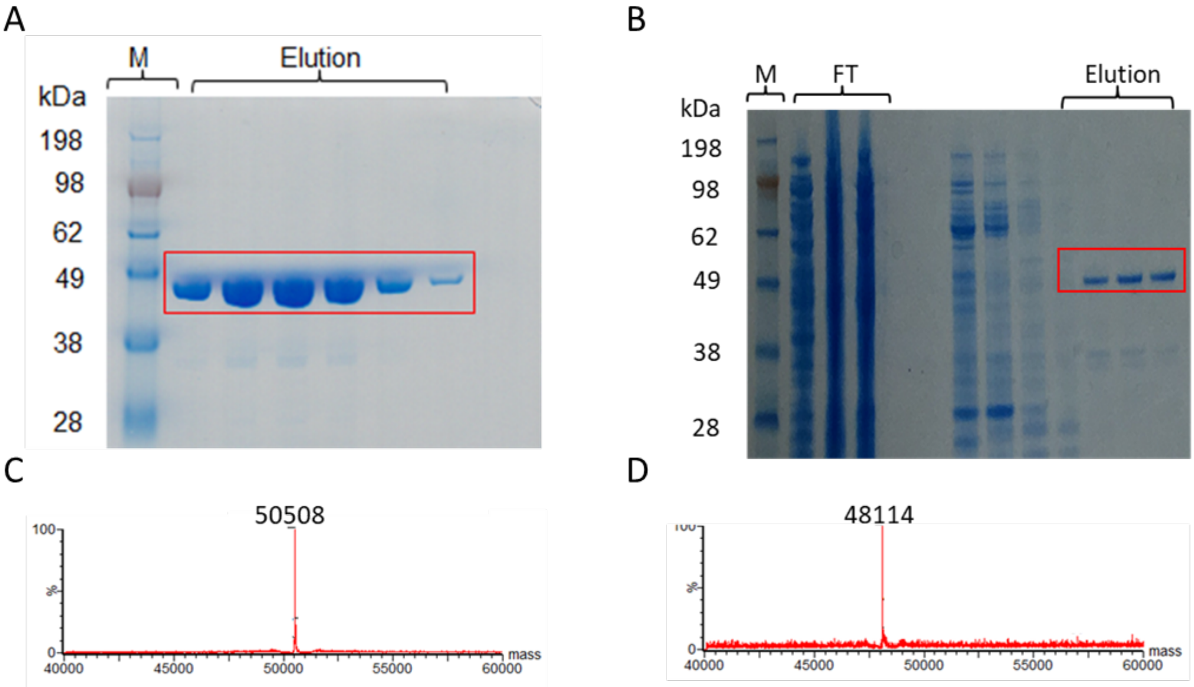

**Supplementary Figure S7. TreT *P. horikoshii* and TreT *T. tenax* Enzyme Characterization and Reaction Analysis.**

Details of TreT homologue purification, characterisation and enzyme reactions. **A**) & **C**). TreT *P. horikoshii* enzyme characterisation by SDS-PAGE and LC-MS. Similarly Figure **B**) & **D**) represent TreT *T. tenax* characterisation by SDS-PAGE and LC-MS.

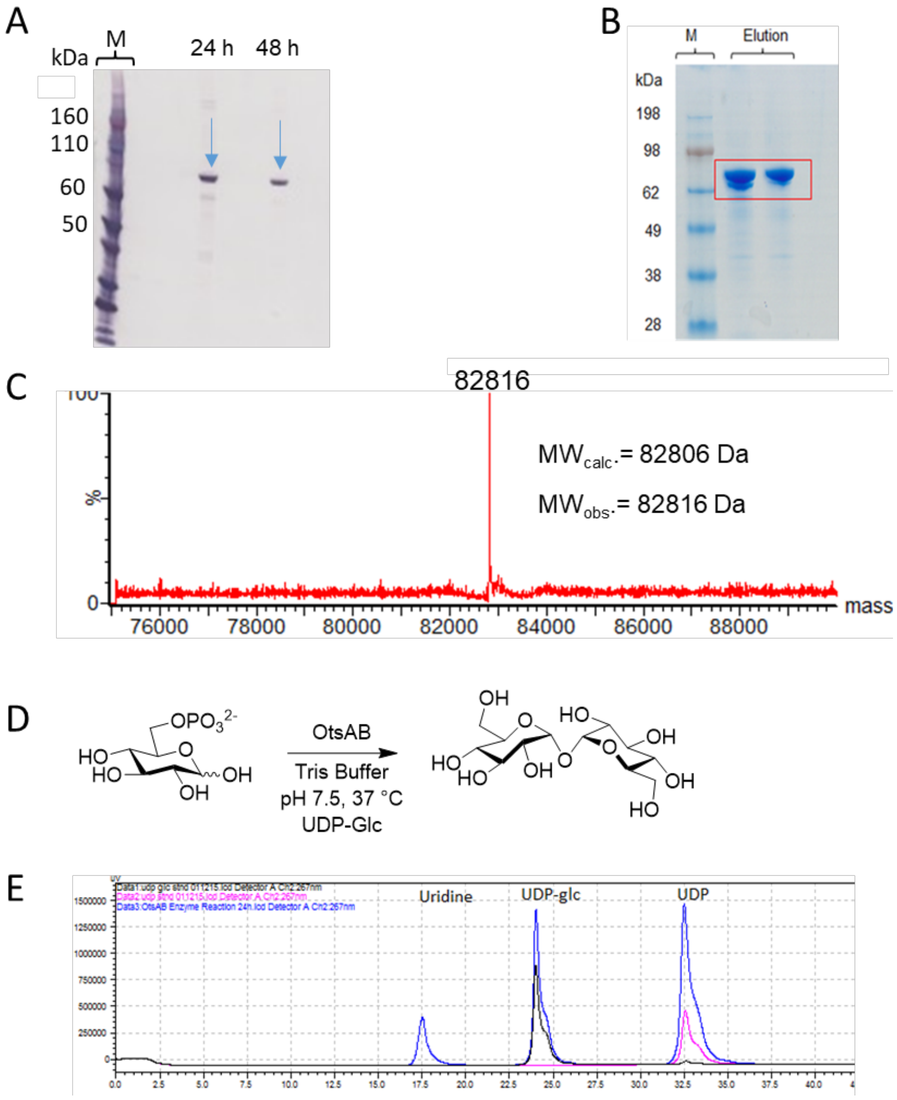

**Supplementary Figure S8. OtsAB fusion Enzyme Optimisation and Expression**.

A) Small scale expression with different IPTG concentrations and $OD_{600}$, values. A His-tagged protein at the expected molecular weight was visualised by Western blotting after probing with anti-His monoclonal antibody after 24 h and 48 h of incubation at 15 °C. IPTG concentration of 0.5 mM and $OD_{600}$ of 0.4 were the relatively best conditions as compared to IPTG concentration 1 mM and $OD_{600}$ 0.6. B) SDS-PAGE characterisation of the large scale (1L) expression of protein. C) LC-MS characterisation of OtsAB fusion enzyme where the expected mass after N-terminus Met cleavage is observed D) Reaction scheme for the enzyme reaction of OtsAB fusion enzyme. and E). HPLC chromatogram of the glucose-6-phosphate reaction in the presence of UDP-Glc and OtsAB fusion enzyme. The reaction monitoring by HPLC confirmed the UDP release which is one of the product of this enzyme catalysis.

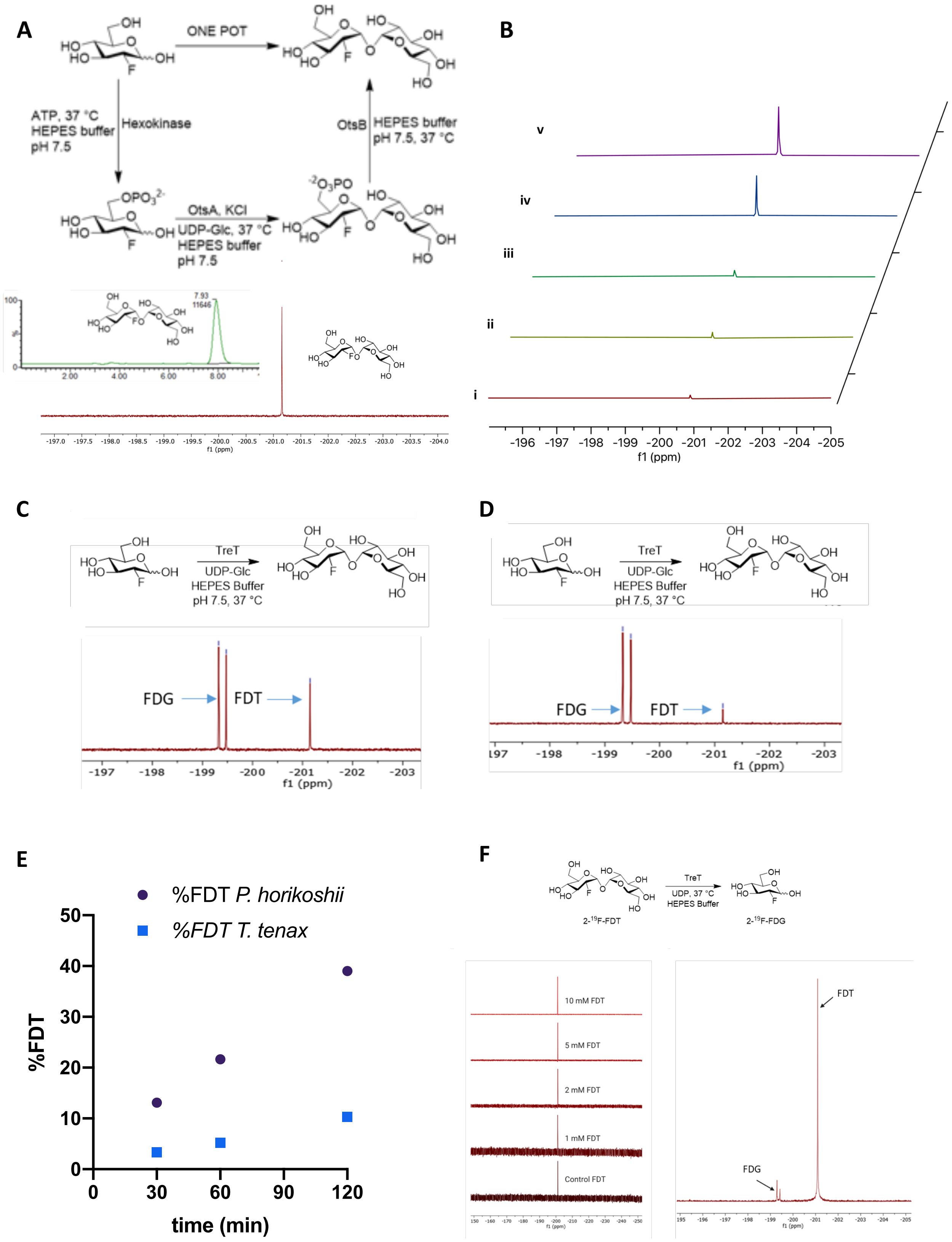
A
ONE POT
ATP, 37 °C
HEPES buffer
pH 7.5
Hexokinase
OtsB
HEPES buffer
pH 7.5, 37 °C
OtsA, KCl
UDP-Glc, 37 °C
HEPES buffer
pH 7.5
7.93
11646
f1 (ppm)
B
v
iv
iii
ii
i
-196 -197 -198 -199 -200 -201 -202 -203 -204 -205
f1 (ppm)
C
TreT
UDP-Glc
HEPES Buffer
pH 7.5, 37 °C
FDG
FDT
-197 -198 -199 -200 -201 -202 -203
f1 (ppm)
D
TreT
UDP-Glc
HEPES Buffer
pH 7.5, 37 °C
FDG
FDT
197 -198 -199 -200 -201 -202 -203
f1 (ppm)
E
%FDT P. horikoshii
%FDT T. tenax
%FDT
time (min)
F
TreT
UDP, 37 °C
HEPES Buffer
2-19F-FDT
2-19F-FDG
10 mM FDT
5 mM FDT
2 mM FDT
1 mM FDT
Control FDT
FDT
FDG
f1 (ppm)

**Supplementary Figure S9. Comparison of different biocatalytic systems for the synthesis of 'cold' FDT on a small scale.**

Routes as described in **Figure 1A**. (**A**) Full conversion of substrate FDG (5.5 mM) into FDT within 45 min as shown by $^{19}$F NMR spectroscopy and LC-MS integrated chromatogram (inset) of [$^{19}$F]FDT reaction sample, showing the formation of the product at rt 7.93 min. (**B**) The reaction was complete on a range of FDG substrate amounts i.e. **i**) 75 µg; **ii**) 100 µg; **iii**) 200 µg; **iv**) 400 µg & **v**) 800 µg, all with full conversion as judged by $^{19}$F NMR spectroscopy to the product FDT within 45 min. (**C & D**) Conversion using two TreT homologue enzyme from *P. horikoshii* and *T. tenax*, respectively (shown at 10 mM) gave an estimated conversion for *P. horikohii* was 40% and for the *T. tenax* ~ 10% as judged by $^{19}$F NMR spectra (shown) when reactions were carried out under the same conditions over 2 h. (**E**) Time course of the reactions shown in (**C**, **D**) using $^{19}$F NMR. (**F**) The *P. horkoshii* TreT enzyme is reversible converting FDT back to FDG, as shown when tested at higher FDT substrate concentrations (20 mM) and higher UDP concentration (80 mM); less 'reversion' was observed under the same conditions at lower UDP (20 mM) and FDT concentrations (1-10 mM).

A

#### OtsA *E. coli N-terminus*

DNA Sequence

ATGAGTCGTTTAGTCGTAGTATCTAACCGGATTGCA
CCACCAGACGAGCACGCCGCCAGTGCCGGTGGCCTT
GCCGTTGGCATACTGGGGGCACTGAAAGCCGCAGGC
GGACTGTGGTTTGGCTGGAGTGGTGAAACAGGGAA

GAGGATCAGCCGCTAAAAAAGGTGAAAAAAGGTAAC

AA Sequence

**M**SRLVVVSNRIAPPDEHAASAGGLAVGILGALKAAG
GLWFGWSGETGNEDQPLKKVKKGN

#### OtsA *E. coli C-terminus*

DNA Sequence

CTGGCGGAACGTATTTCCCGTCATGCAGAAATGCTGGA
CGTTATCGTGAAAAACGATATTAACCACTGGCAGGAGT
GCTTCATTAGCGACCTAAAGCAGATAGTTCCGCGAAGC
GCGGAAAGCCAGCAGCGCGATAAAGTTGCTACCTTTCC

AAAGCTTGCGCTCGAGCACCACCACCACCACCACTGA

AA Sequence

**L**AERISRHAE**M**LDVIVKNDINHWQECFISDLKQIVPRS
AESQQRDKVATFPKLALEHHHHHH-

#### OtsA *Sf9 N-terminus*

DNA Sequence

ATGAGCCGCCTGGTCGTGGTGTCCAACCGCATCGCTCCCCC
TGACGAACATGCCGCTAGCGCTGGTGGTCTCGCTGTCGGTA
TCCTCGGCGCTCTGAAGGCTGCCGGTGGTCTGTGGTTTGGC
TGGAGTGGTGAAACAGGGAATGAGGATCAGCCGCTAAAAAA
GGTGAAAAAA GGCAAC

AA Sequence

**M**SRLVVVSNRIAPPDEHAASAGGLAVGILGALKAAGGLWFG
WSGETGNEDQPLKKVKKGN

#### OtsA *Sf9 C-terminus*

DNA Sequence

CTGGCTGAACGTATCTCCCGCCATGCCGAGATGCTGG
ACGTCATCGTGAAGAACGATATTAACCACTGGCAGGA
GTGCTTCATTAGCGACCTAAAGCAGATAGTTCCGCGA
AGCGCTGAGAGCCAGCAGCGCGACAAGGTGGCCACCT
TCCCTAAGCTGGCCCTGGAGCACCAT
CACCACCACCACGCG

AA Sequence

**L**AERISRHAE**M**LDVIVKNDINHWQECFISDLKQIVPR
SAESQQRDKVATFPKLALEHHHHHH

B

#### OtsB *E. coli N-terminus*

DNA Sequence

ATGGGCAGCAGCCACCACCACCACCACCACGGCACCAAGAC
CGAGGAGGGCAAGCTGGTGATCTGGATCAACGGCGACAAGG
GCTACAACGGCCTGGCCGAGGTGGGCAAGAAGTTCGAGAAG
GACACCGGCATCAAGGTGACCGTGGAGCACCCCGACAAGCT
GGAGGAGAAGTTCCCCCTGGCCGAGATCAAGCCCCACCCCG
ACCAGGTGGTGGTG

AA Sequence

MGSSHHHHHHGTKTEEGKLVIWINGDKGYNGLAEVGKKFEK
DTGIKVTVEHPDKLEEKFPLAEIKPHPDQVVV

#### OtsB *E. coli C-terminus*

DNA Sequence

GGCTTCGCCGTGGTGAACAGGCTGGGCGGCATGAGCG
TGAAGATCGGCACCGGCGCCACCCAGGCCAGCTGGAG
GCTGGCCGGCGTGCCCGACGTGTGGAGCTGGCTGGAG
ATGATCACCACCGCCCTGCAGCAGAAGAGGGAGAACA
ACAGGAGCGACGACTACGAGAGCTTCAGCAGGAGCAT
C

AA Sequence

GFAVVNRLGGMSVKIGTGATQASWRLAGVPDVWSWLE
MITTALQQKRENNRSDDYESFSRSI

#### OtsB *Sf9 N-terminus*

DNA Sequence

ATGAGCTACTACCACCACCACCACCACCACGACTACGACAT
CCCCACCACCGAGAACCTGTACTTCCAGGGCCACATGGTGA
CCGAGCCCCTGACCGAGACCCCCGAGCTGAGCGCCAAGTAC
GCCTGGTTCTTCGACCTGGACGGCACCCTGGCCGAGATCAA
GCCCCACCCCGACCAGGTGGTGGTG

AA Sequence

SYYHHHHHHDYDIPTTENLYFQGHMVTEPLTETPELSAKYA
WFFDLDGTLAEIKPHPDQVVV

#### OtsB *Sf9 C-terminus*

DNA Sequence

GGCTTCGCCGTGGTGAACAGGCTGGGCGGCATGAGCG
TGAAGATCGGCACCGGCGCCACCCAGGCCAGCTGGAG
GCTGGCCGGCGTGCCCGACGTGTGGAGCTGGCTGGAG
ATGATCACCACCGCCCTGCAGCAGAAGAGGGAGAACA
ACAGGAGCGACGACTACGAGAGCTTCAGCAGGAGCAT
C

AA Sequence

GFAVVNRLGGMSVKIGTGATQASWRLAGVPDVWSWLE
MITTALQQKRENNRSDDYESFSRSI

**Supplementary Figure S10. Sequence Comparison of OtsA and OtsB Sources.**

**A)** Comparison of N-terminus and C-terminus residues of OtsA sequences from both *E. coli* and *Sf9*. **B)** Comparison of N-terminus and C-terminus residues of OtsB sequences from both *E. coli* and *Sf9*.

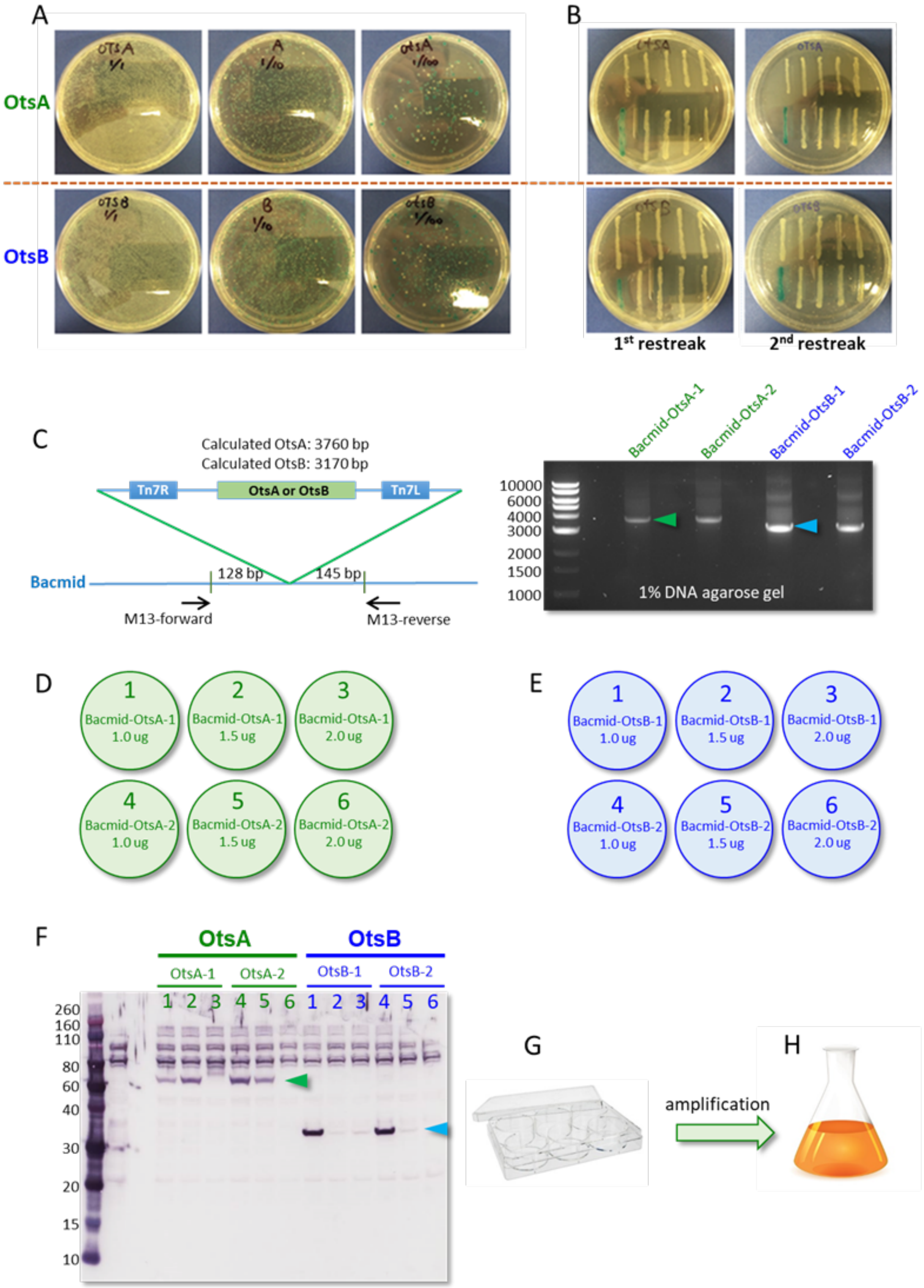
A
B
OtsA
OtsB
1st restreak
2nd restreak
C
Calculated OtsA: 3760 bp
Calculated OtsB: 3170 bp
Tn7R
OtsA or OtsB
Tn7L
Bacmid
128 bp
145 bp
M13-forward
M13-reverse
Bacmid-OtsA-1
Bacmid-OtsA-2
Bacmid-OtsB-1
Bacmid-OtsB-2
10000
6000
4000
3000
2000
1500
1000
1% DNA agarose gel
D
1 Bacmid-OtsA-1 1.0 ug
2 Bacmid-OtsA-1 1.5 ug
3 Bacmid-OtsA-1 2.0 ug
4 Bacmid-OtsA-2 1.0 ug
5 Bacmid-OtsA-2 1.5 ug
6 Bacmid-OtsA-2 2.0 ug
E
1 Bacmid-OtsB-1 1.0 ug
2 Bacmid-OtsB-1 1.5 ug
3 Bacmid-OtsB-1 2.0 ug
4 Bacmid-OtsB-2 1.0 ug
5 Bacmid-OtsB-2 1.5 ug
6 Bacmid-OtsB-2 2.0 ug
F
OtsA
OtsB
OtsA-1
OtsA-2
OtsB-1
OtsB-2
1 2 3 4 5 6 1 2 3 4 5 6
260
160
110
80
60
40
30
20
15
10
G
amplification
H

**Supplementary Figure S11. Schematic of the insect cell (Sf9) expression of OtsA$^{Sf}$ and OtsB$^{Sf}$.**

**A**) Blue/white selection screening of OtsA and OtsB recombinant bacmid DNAs. **B**) Phenotype confirmation by streaking and re-streaking the selected colonies from A). **C**) PCR analysis to confirm the insertion of *otsA* and *otsB* gene into bacmid DNA. **D** & **E**) Generation of P1 baculovirus for OtsA and OtsB enzymes by Sf9 cell transfection with different amounts of bacmid DNA in a 6-well plate. **F**) Western blot analysis using *anti*-His tag antibody to characterize the expression of OtsA and OtsB proteins based on the cell pellets obtained during the harvest of their P1 baculovirus, respectively. **G** & **H**) Amplification of the P1 baculovirus of OtsA and OtsB to generate the P2 baculovirus stock.

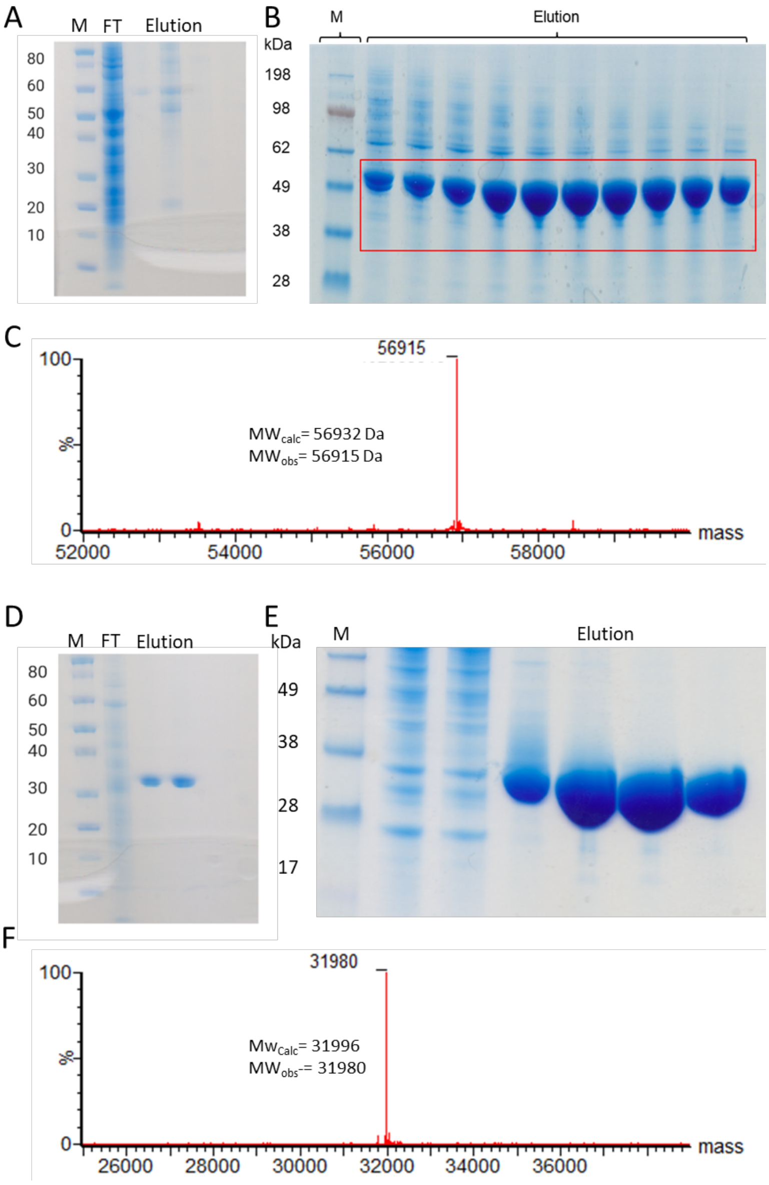

**Supplementary Figure S12. Small and Large scale Expression of OtsA and OtsB enzymes from Insect cells.**

**A**) SDS-PAGE characterization of OtsA enzyme from the pellet of 50 mL culture. **B**) SDS-PAGE characterization of OtsA enzyme from 1 L culture. **C**) LC-MS characterization of OtsA enzyme. **D**) SDS-PAGE characterization of OtsB enzyme from the pellet of 50 mL culture. **E**) SDS-PAGE characterization of OtsB enzyme from the 1L scale expression culture. **F**) LC-MS characterization of OtsB enzyme.

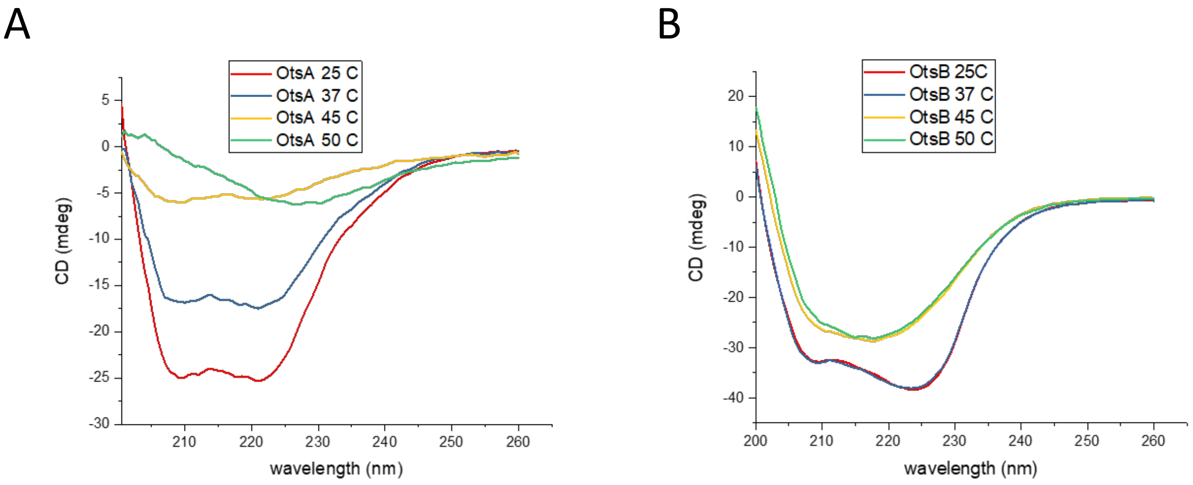

**Supplementary Figure S13. CD spectra and Stability Assessment of OtsA and OtsB enzymes.**

**A**). CD spectra of OtsA enzyme incubated at 25 °C, 37 °C, 45 °C and 50 °C for 1h. **B**). CD spectra of OtsB enzyme at 25 °C, 37 °C, 45 °C and 50 °C under the same conditions. Both enzymes maintained secondary structure at 25 °C and 37 °C, however, at higher temperatures secondary structure of protein was lost.

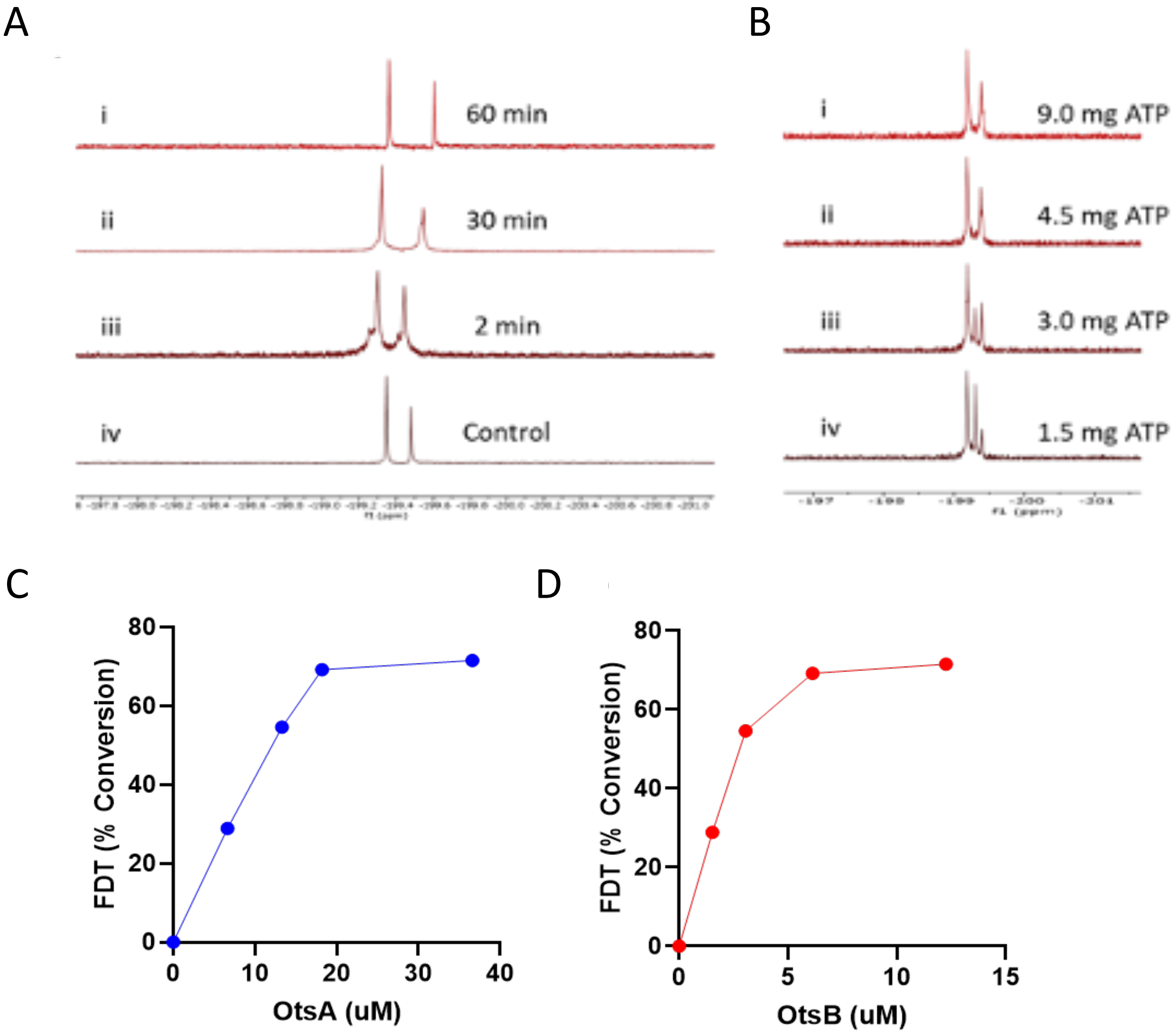

**Supplementary Figure S14. Optimisation Screen during Enzyme Reaction Scale-up.**

**A, B**) Testing the optimal amount of hexokinase and ATP needed for full conversion of FDG to FDG6P. **A**). Reaction monitoring from FDG to FDG6P using 5 mg hexokinase. Reaction completed after 60 min. **B**). Small scale reactions of FDG with ATP to determine the optimal amount of ATP needed for conversion of FDG to FDG6P. The amount of FDG was fixed at 5.6 mM or 1 mg, whereas ATP varied from 2.7 mM (1.5 mg), 5.4 mM (3 mg), 8.1 mM (4.5 mg) and 16.2 mM (9 mg). In conditions i and ii where concentration of ATP was higher than FDG, full conversion to FDG6P observed by $^{19}$F-NMR spectroscopy.

**C, D**) OtsA and OtsB enzyme optimum concentration determination. For optimum enzyme concentration determination, reaction with fixed substrate concentrations and all the other parameters incubated with varying enzyme(s) concentrations. **C**). OtsA concentrations varied from 6.65 µM to 36.6 µM. Optimum concentration of the enzyme was found to be 18.2 µM. **D**). OtsB concentrations varied from 1.53 µM to 12.26 µM. Optimum concentration was 6.1 µM.

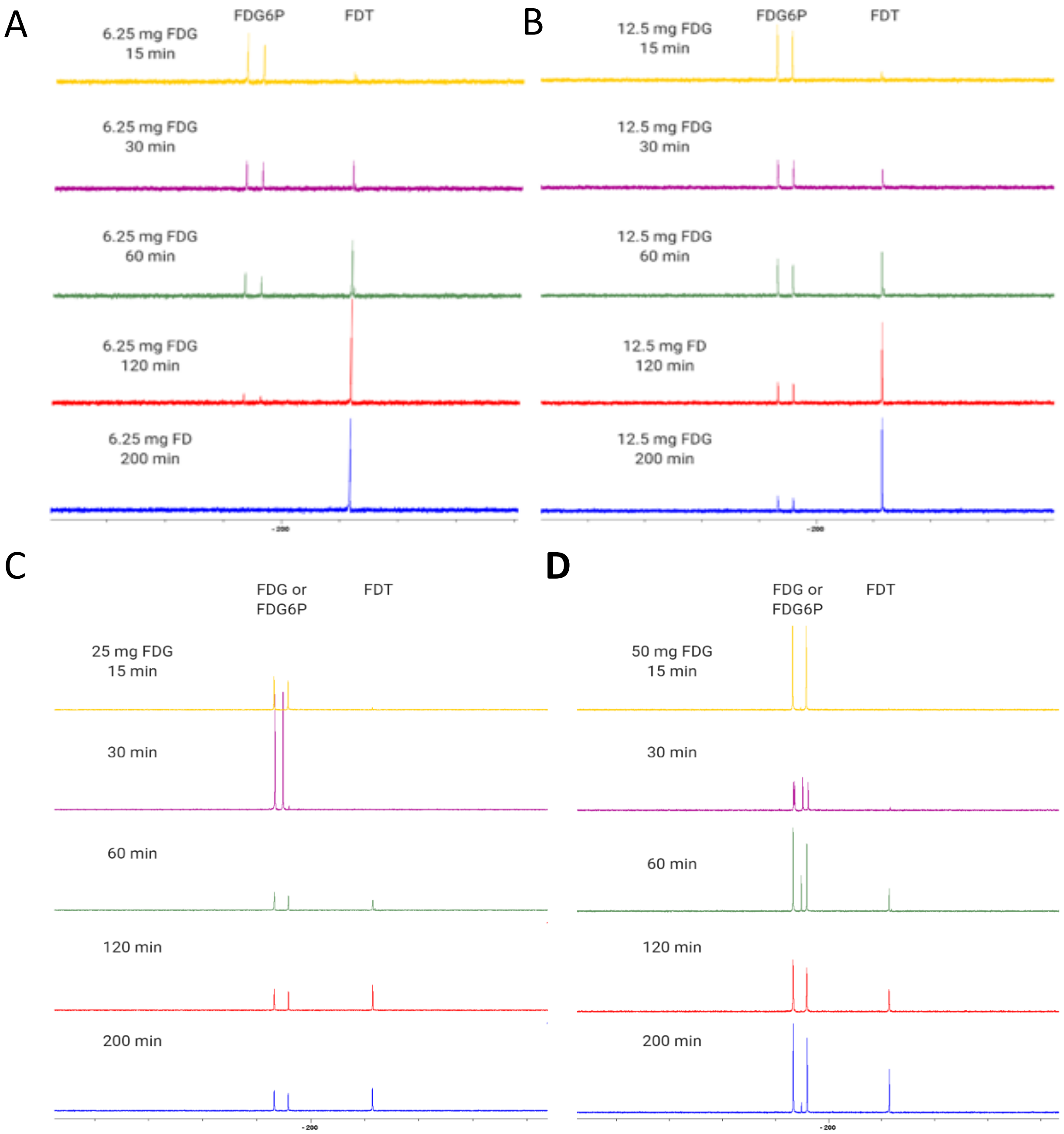

**Supplementary Figure S15. [$^{19}$F]FDT Reaction Optimisation and Monitoring by $^{19}$F-NMR spectroscopy.**

**A**). 6.25 mg FDG reaction catalysis by hexokinase, OtsA and OtsB. Samples were taken 15 min, 30 min, 60 min, 120 min and 200 min post reaction and reaction was more than 95% complete after 2 h. **B**). 12.5 mg FDG sample with similar monitoring time as above. After 3 h, there was 10% of FDG6P remained uncatalysed. **C**). 25 mg FDG sample catalysed and reaction monitored as A, after 3h, 20 min there was about 45% intermediate remained uncatalysed and **D**). 50 mg FDG sample catalysed and analysed as above, the conversion was 30% after 3h, 20 min. The glycosylation reaction was the limiting step and the amount of UDP-Glc was kept constant at 30 mM which was significantly less for 25 mg FDG substrate and 50 mg FDG substrate.

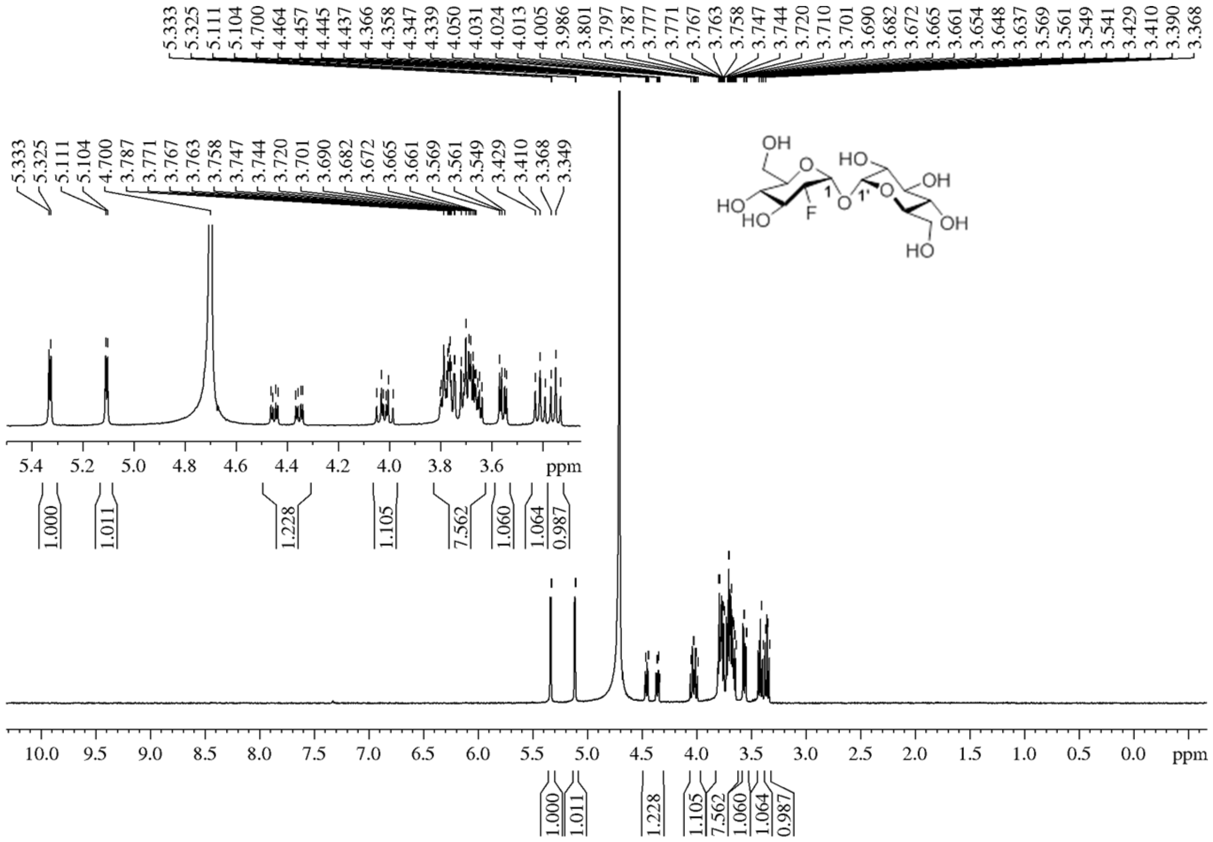

**Supplementary Figure S16. $^{1}$H NMR spectrum of [$^{19}$F]-FDT**

| | |
|---|---|
| Frequency [MHz) | 500.13 |
| SW [ppm] | 19.9947 |
| Origin | AVX500 |
| Nucleus | $^{1}$H |
| NS | 16 |
| Solvent | $D_2O$ |
| Temperature (degree C) | 25 |

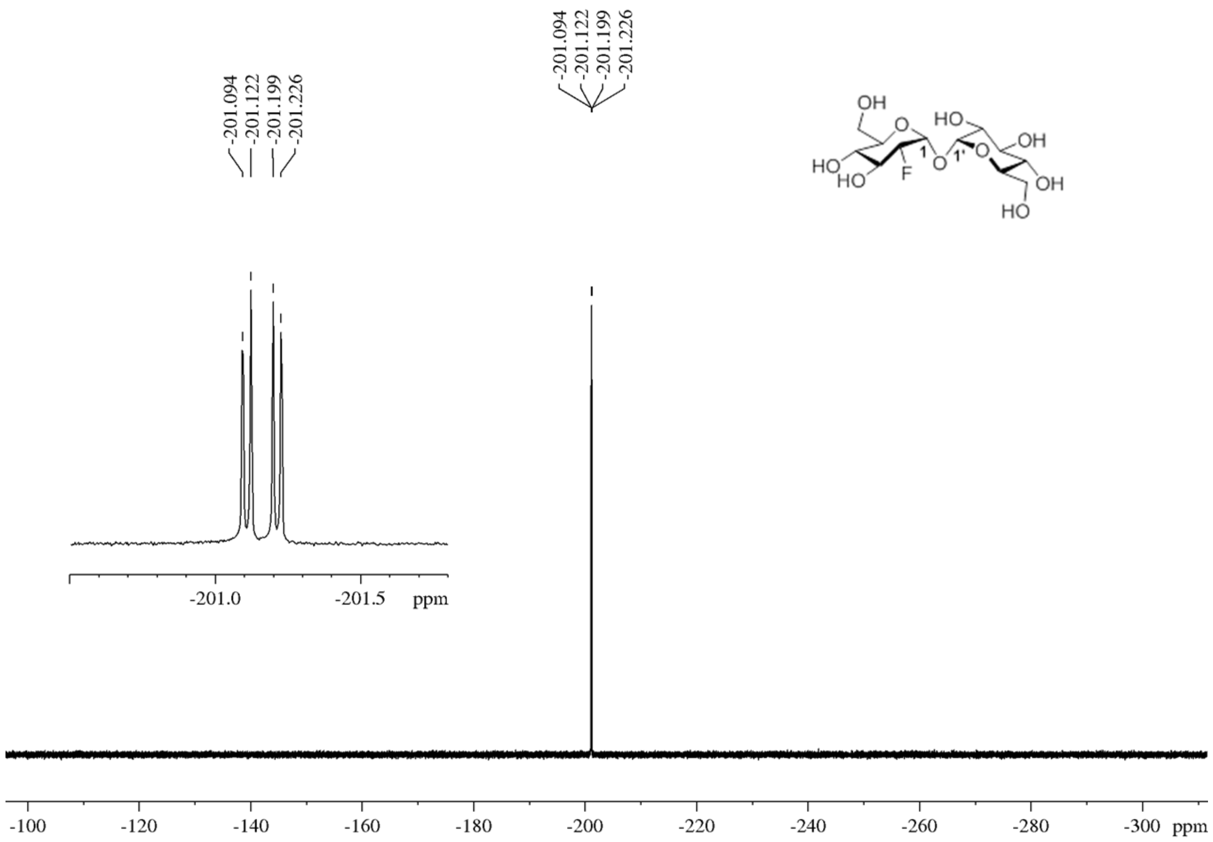

**Supplementary Figure S17. $^{19}$F($^{1}$H coupled) NMR spectrum of [$^{19}$F]-FDT**

| | |
|---|---|
| Frequency [MHz) | 470.13 |
| SW [ppm] | 241.5235 |
| Origin | AVX500 |
| Nucleus | $^{19}$F |
| NS | 256 |
| Solvent | $D_2O$ |
| Temperature (degree C) | 25 |

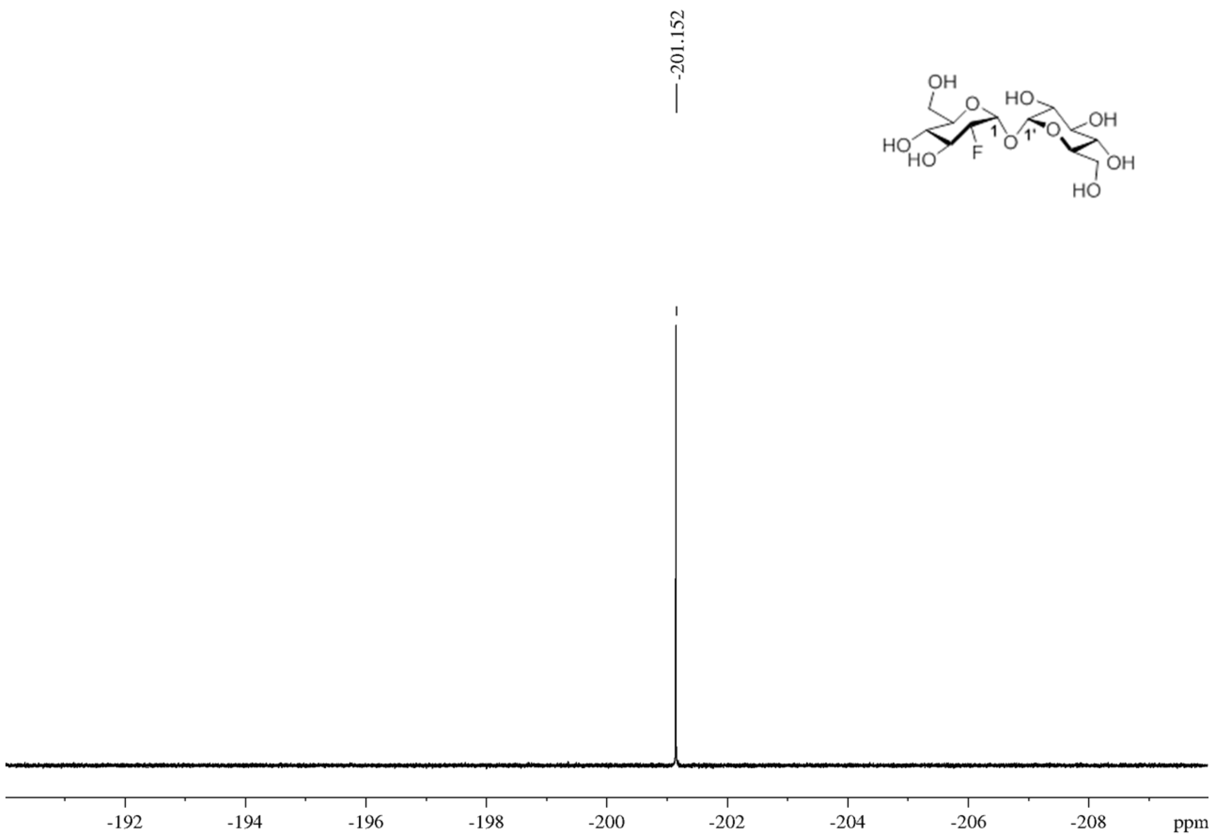

**Supplementary Figure S18. $^{19}$F($^{1}$H decoupled) NMR spectrum of [$^{19}$F]- FDT**

| | |
|---|---|
| Frequency [MHz) | 470.13 |
| SW [ppm] | 19.9756 |
| Origin | AVX500 |
| Nucleus | $^{19}$F |
| NS | 32 |
| Solvent | $D_2O$ |
| D1 (sec) | 10 |
| Temperature (degree C) | 25 |

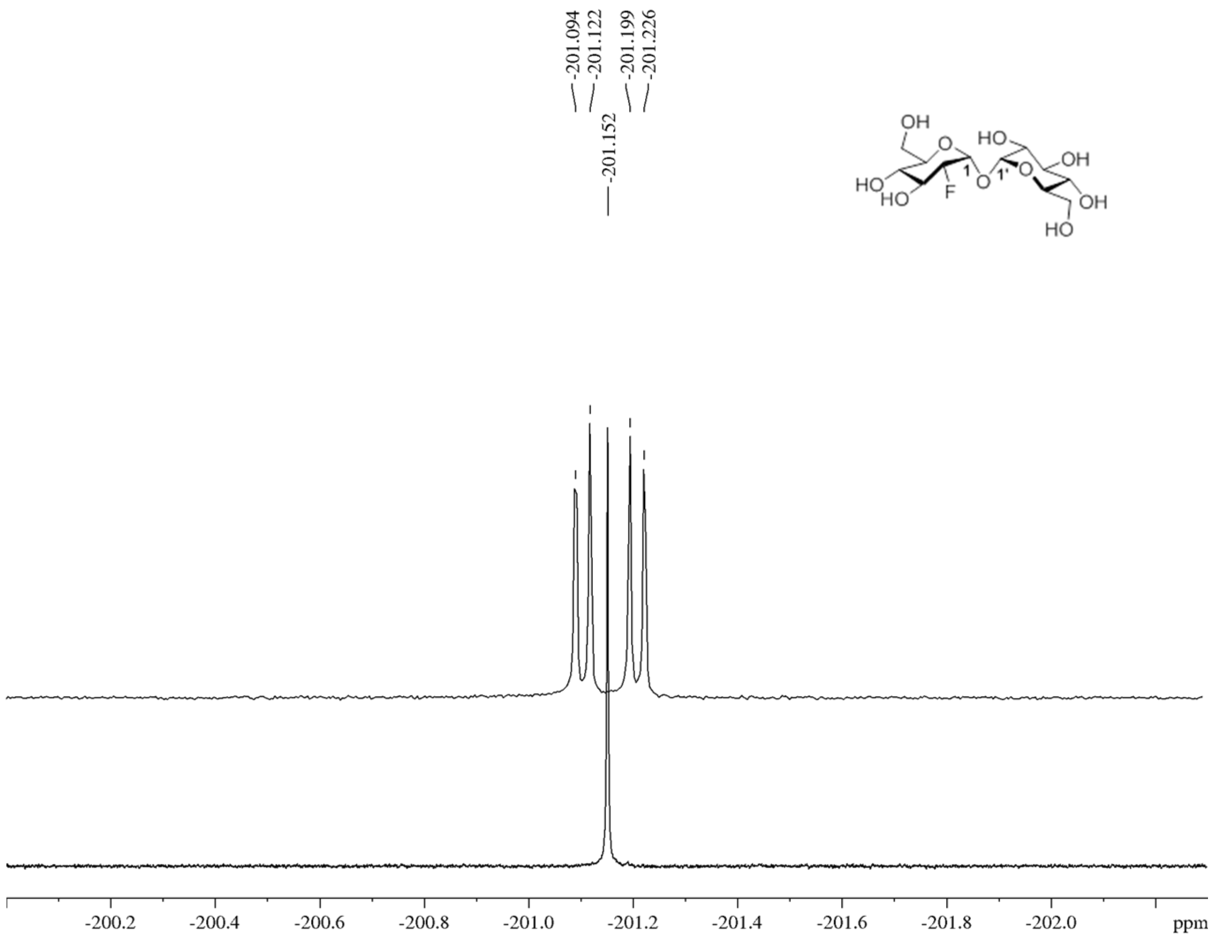

**Supplementary Figure S19. $^{19}$F and $^{19}$F($^{1}$H) Coupled NMR spectrum of [$^{19}$F]-FDT.**

*Both coupled and decoupled spectra are stacked together.*

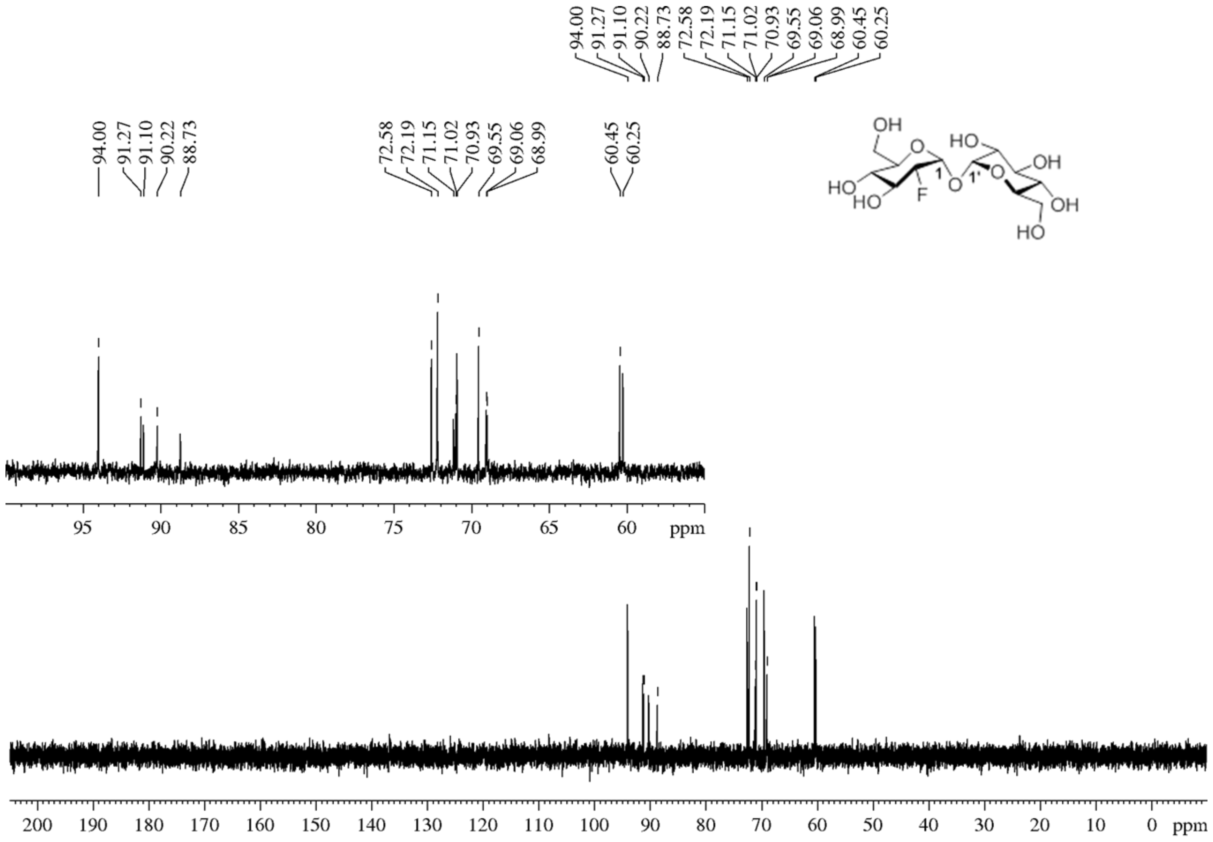

**Supplementary Figure S20. $^{13}$C NMR spectrum of [$^{19}$F]-FDT**

| | |
|---|---|
| Frequency [MHz) | 126 |
| SW [ppm] | 236.64 |
| Origin | AVX500 |
| Nucleus | $^{13}$C |
| NS | 3072 |
| Solvent | $D_2O$ |
| D1 (sec) | 2 |
| Temperature (degree C) | 25 |

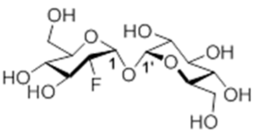

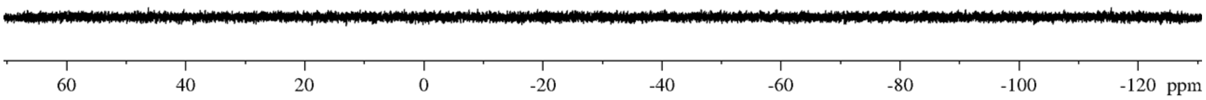

**Supplementary Figure S21. $^{31}$P($^{1}$H Decoupled) NMR Spectrum of [$^{19}$F]-FDT.**

*No signal for either reaction intermediates nor reagents used that contain $^{31}$P nuclei.*

| | |
|---|---|
| Frequency [MHz) | 202 |
| SW [ppm] | 201.3377 |
| Origin | AVX500 |
| Nucleus | $^{31}$P |
| NS | 64 |
| Solvent | $D_2O$ |
| D1 (sec) | |
| Temperature (degree C) | 25 |

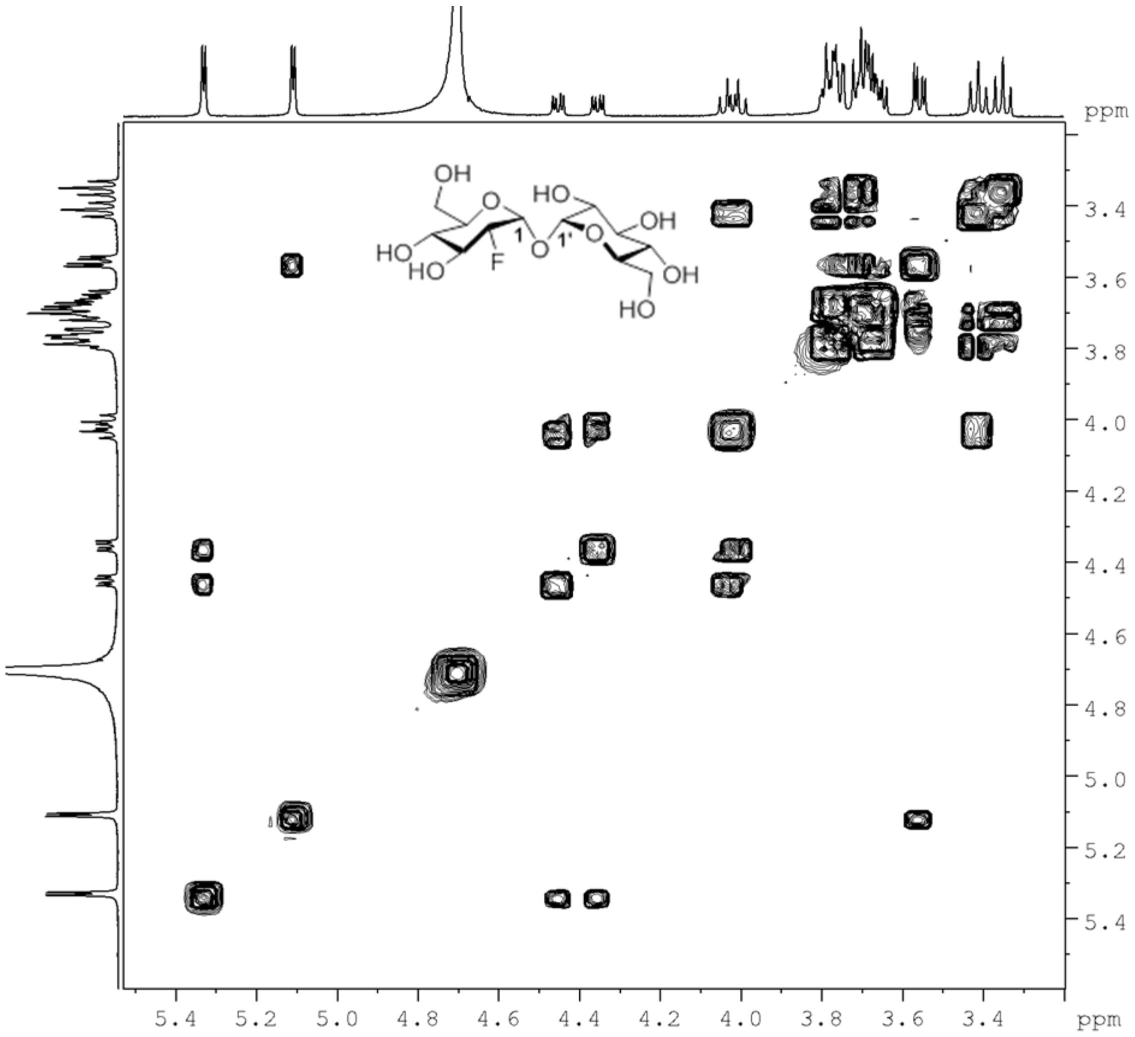

**Supplementary Figure S22. COSY spectrum of [$^{19}$F]-FDT.**

*$^{1}$H-$^{1}$H COSY (Correlated Spectroscopy).*

| | |
|---|---|
| Frequency [MHz) | 500 |
| SW [ppm] | 12.0161 |
| Origin | AVX500 |
| Nucleus | $^{1}$H |
| NS | 8 |
| Solvent | $D_2O$ |

| | |
|---|---|
| D1 (sec) | 2 |
| Temperature (degree C) | 25 |

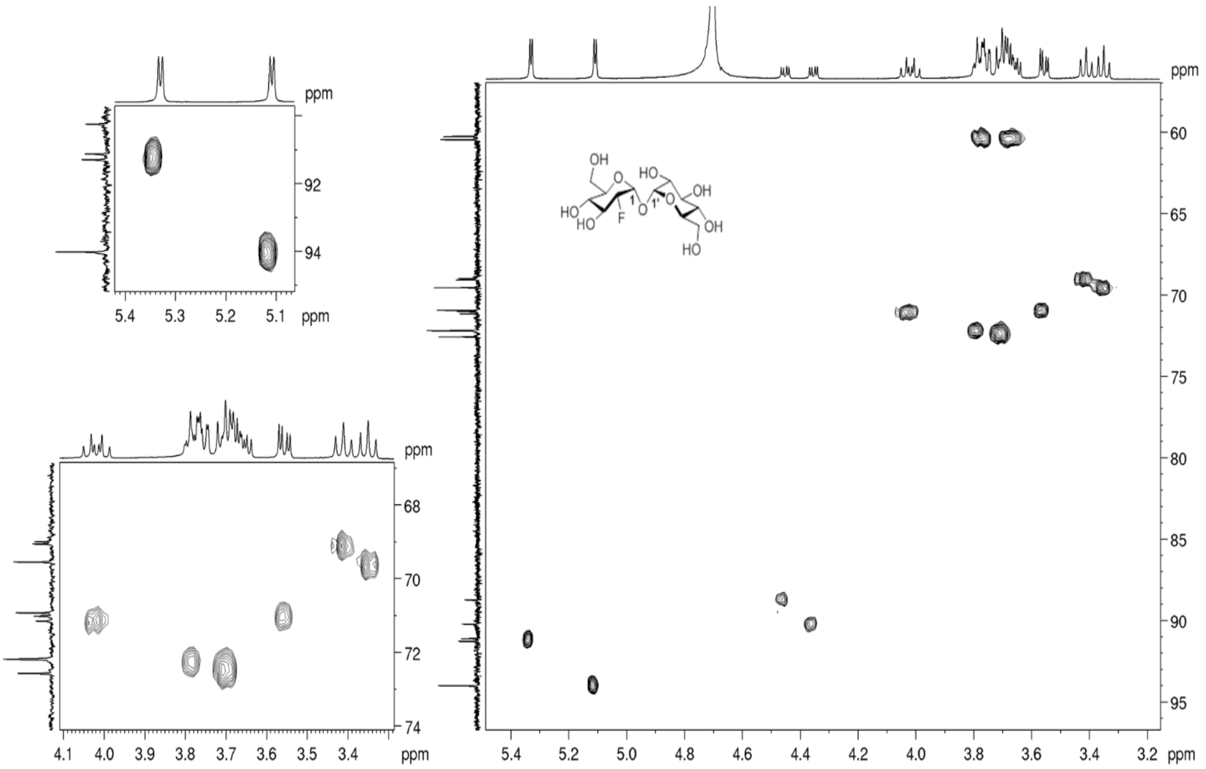

**Supplementary Figure S23. HSQC Spectrum of [$^{19}$F]-FDT.**

Heteronuclear Single Quantum Coherence experiment was used to determine proton-carbon single bond correlations. In the above spectrum $^{13}$C NMR spectrum is on the Y-axis whereas $^{1}$H NMR spectrum is shown on the X-axis.

| | |
|---|---|
| Frequency [MHz) | $^{1}$H 500 & $^{13}$C 126 |
| SW [ppm] | 12.0161 |
| Origin | AVX500 |
| Nucleus | $^{1}$H – $^{13}$C |
| NS | 8 |
| Solvent | $D_2O$ |
| D1 (sec) | 1 |
| Temperature (degree C) | 25 |

X:\data\Jan 19\ESI74212.raw 07/01/2019 1:46 pm

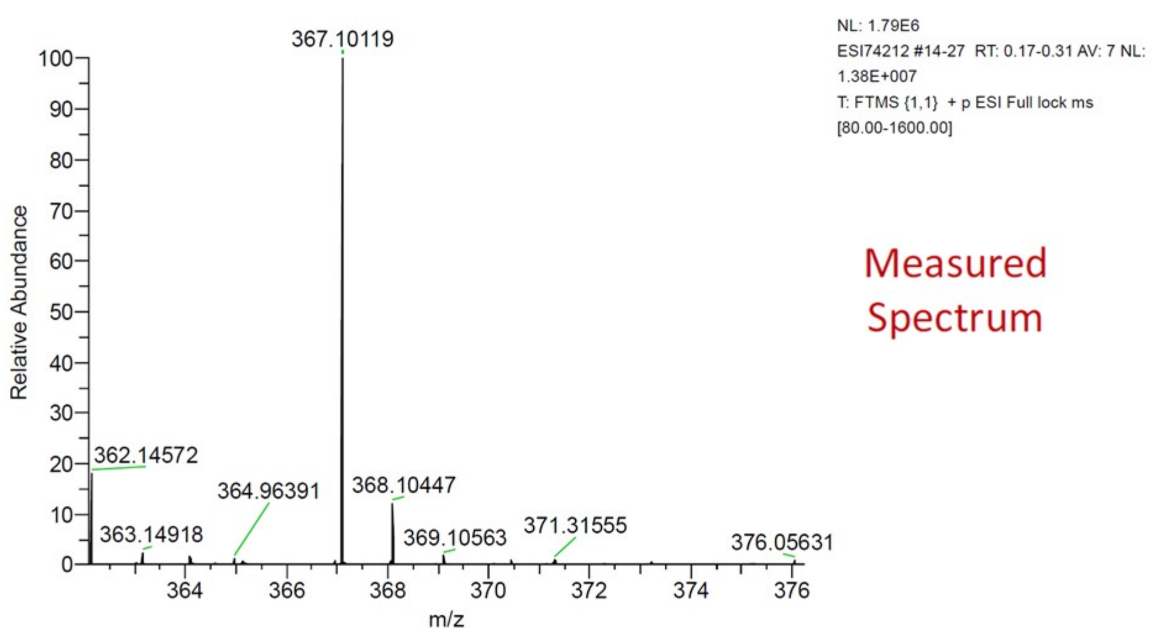

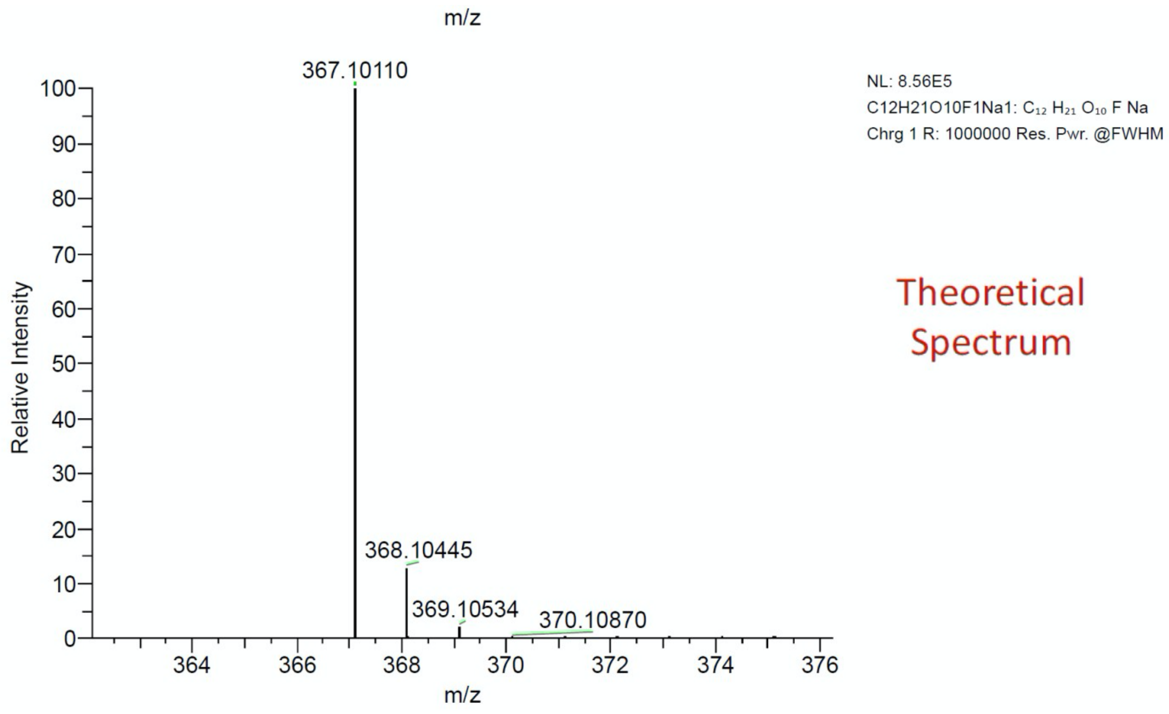

**Supplementary Figure S24. ESI-MS Spectrum of [$^{19}$F]- FDT.**

40 µg of [$^{19}$F]- FDT Sample was dissolved in 1 mL 50:50 methanol de-ionised water and the accurate mass of the sample was determined. The instrument is accurate up to 4 decimal points.

The sample was run on a Thermo Orbitrap Exactive mass spectrometer calibrated with standard calibration solutions that included MRFA (L-methionyl-arginyl-phenylalanyl-alanine acetate.x$H_2O$), Ultramark 1621, caffeine and glacial acetic acid.

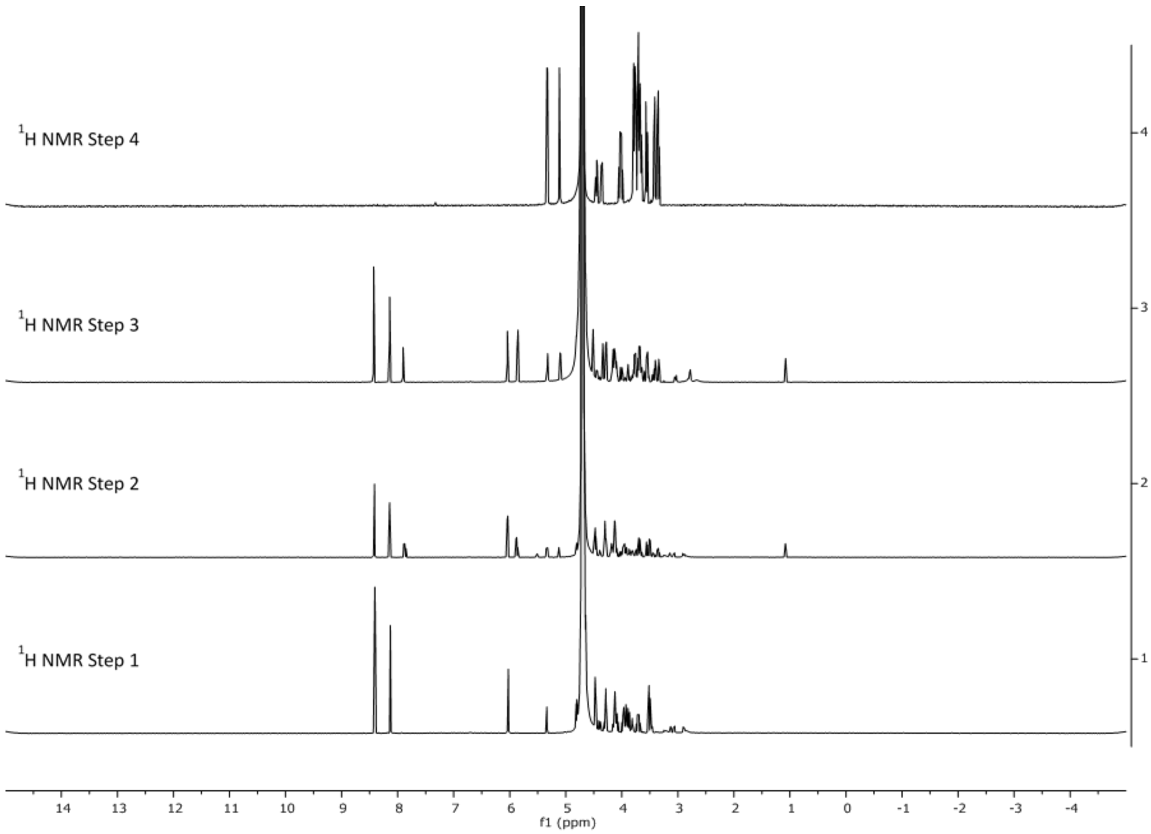

**Supplementary Figure S25. $^1$H-NMR (500 MHz) spectra of crude reaction mixtures during one-pot synthesis and purified product.**

Step 1 involves the conversion of **FDG** to **FDG6P**; Step 2 represents conversion of **FDG6P** to **FDT6P**; Step 3 shows the conversion of **FDT6P** to **[$^{19}$F]- FDT**. Step 4 is for the purified final batch of **[$^{19}$F]-FDT**. $D_2O$ was used as solvent. For all crude reactions, 20 µL sample was diluted in 500 µL of $D_2O$; whereas for the purified sample, 2.3 mg of **[$^{19}$F]-FDT** was dissolved in 500 µL of $D_2O$.

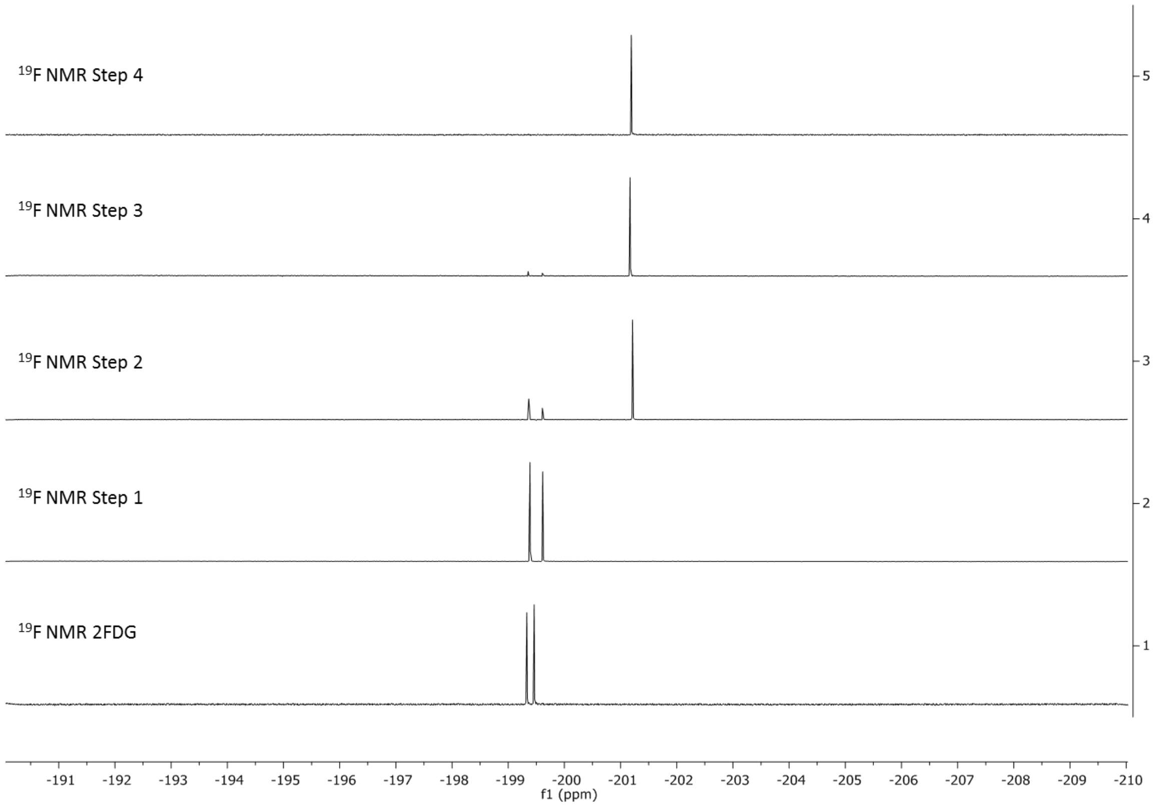

**Supplementary Figure S26. $^{19}$F(Decoupled) NMR (470 MHz) spectra of crude mixture and purified product of [$^{19}$F]-FDT.**

Step 1 involves the conversion of **FDG** to **FDG6P**; Step 2 represents conversion of **FDG6P** to **FDT6P**; Step 3 shows the conversion of **FDT6P** to **[$^{19}$F]-FDT**. Step 4 is for the purified final batch of **[$^{19}$F]-FDT**. $D_2O$ was used as solvent. For all crude reactions, 20 µL sample was diluted in 500 µL of $D_2O$; whereas for the purified sample, 2.3 mg of **[$^{19}$F]-FDT** was dissolved in 500 µL of $D_2O$.

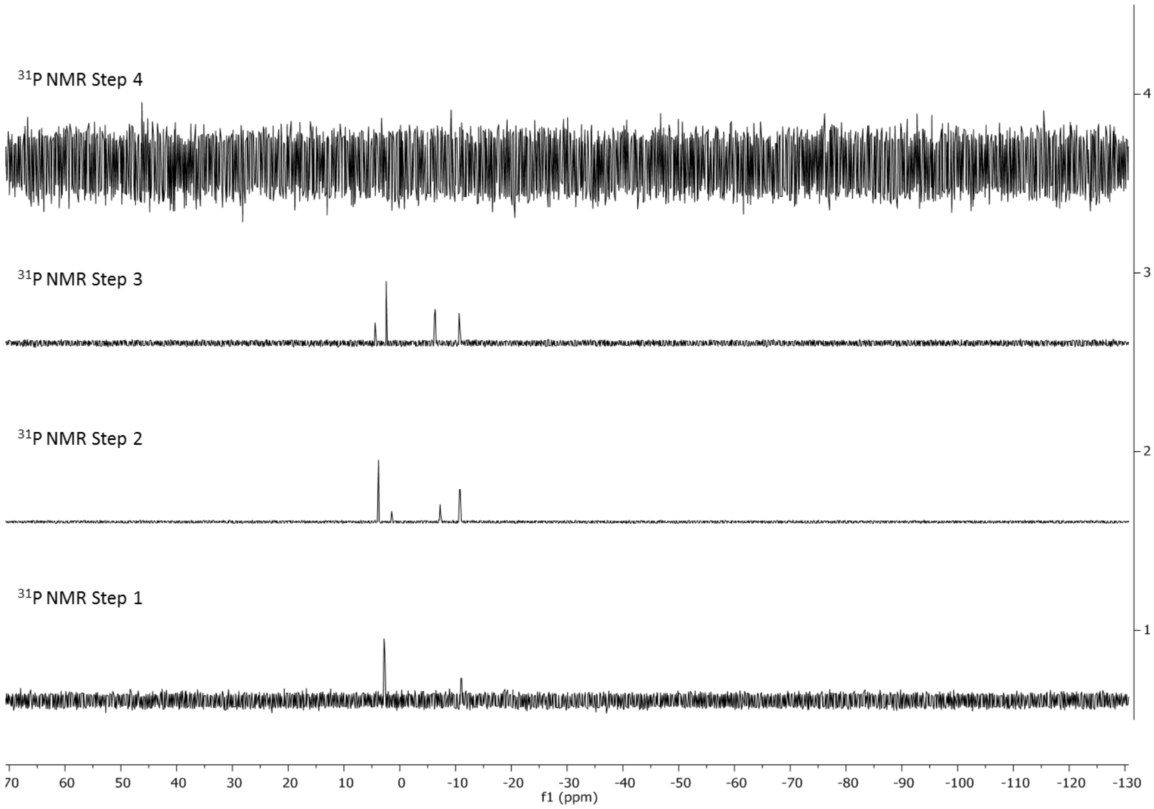

**Supplementary Figure S27. $^{31}$P-NMR (202 MHz) spectra of crude mixture and purified batch of [$^{19}$F]-FDT.**

Step 1 involves the conversion of **FDG** to **FDG6P**; Step 2 represents conversion of **FDG6P** to **FDT6P**; Step 3 shows the conversion of **FDT6P** to **[$^{19}$F]-FDT**. Step 4 is for the purified final batch of **[$^{19}$F]-FDT**. $D_2O$ was used as solvent. For all crude reactions, 20 μL sample diluted in 500 μL of $D_2O$; whereas for the purified sample, 2.3 mg of **[$^{19}$F]-FDT** was dissolved in 500 μL in $D_2O$.

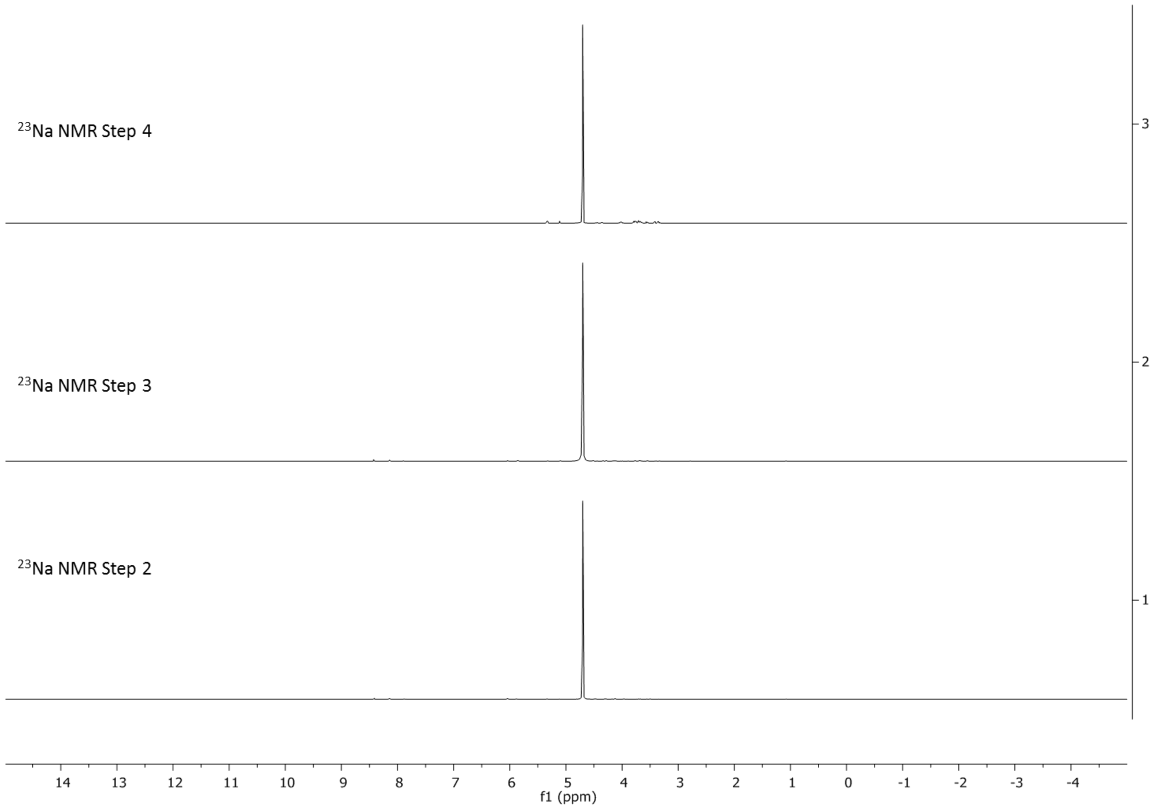

**Supplementary Figure S28. $^{23}$Na-NMR (132 MHz) Spectra of FDT6P & [$^{19}$F]-FDT Crude and [$^{19}$F]-FDT purified batch.**

Step 2 is from **FDT6P** crude mixture; Step 3 is from **[$^{19}$F]-FDT** crude mixture; Step 4 is from the purified product of **[$^{19}$F]-FDT**. All samples were analyzed in $D_2O$.

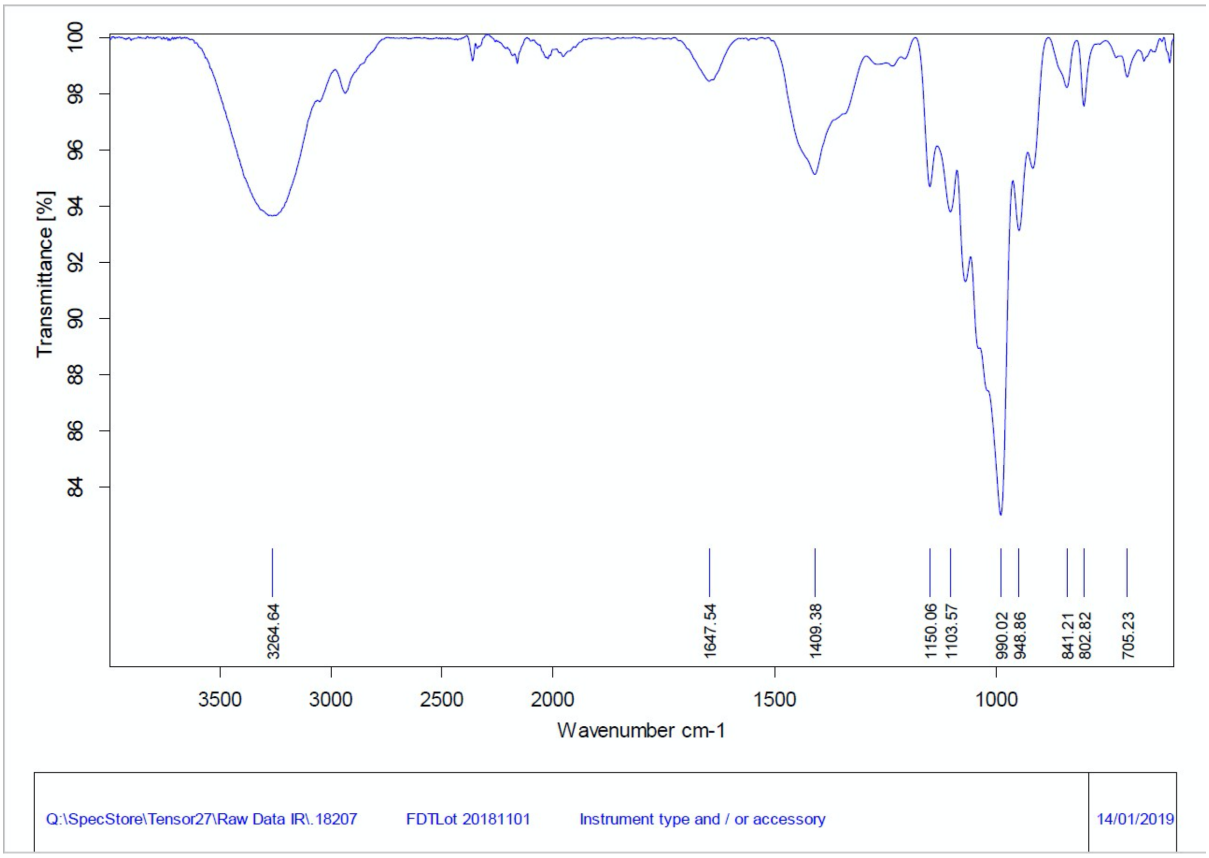

**Supplementary Figure S29. IR Spectrum of [$^{19}$F]-FDT.**

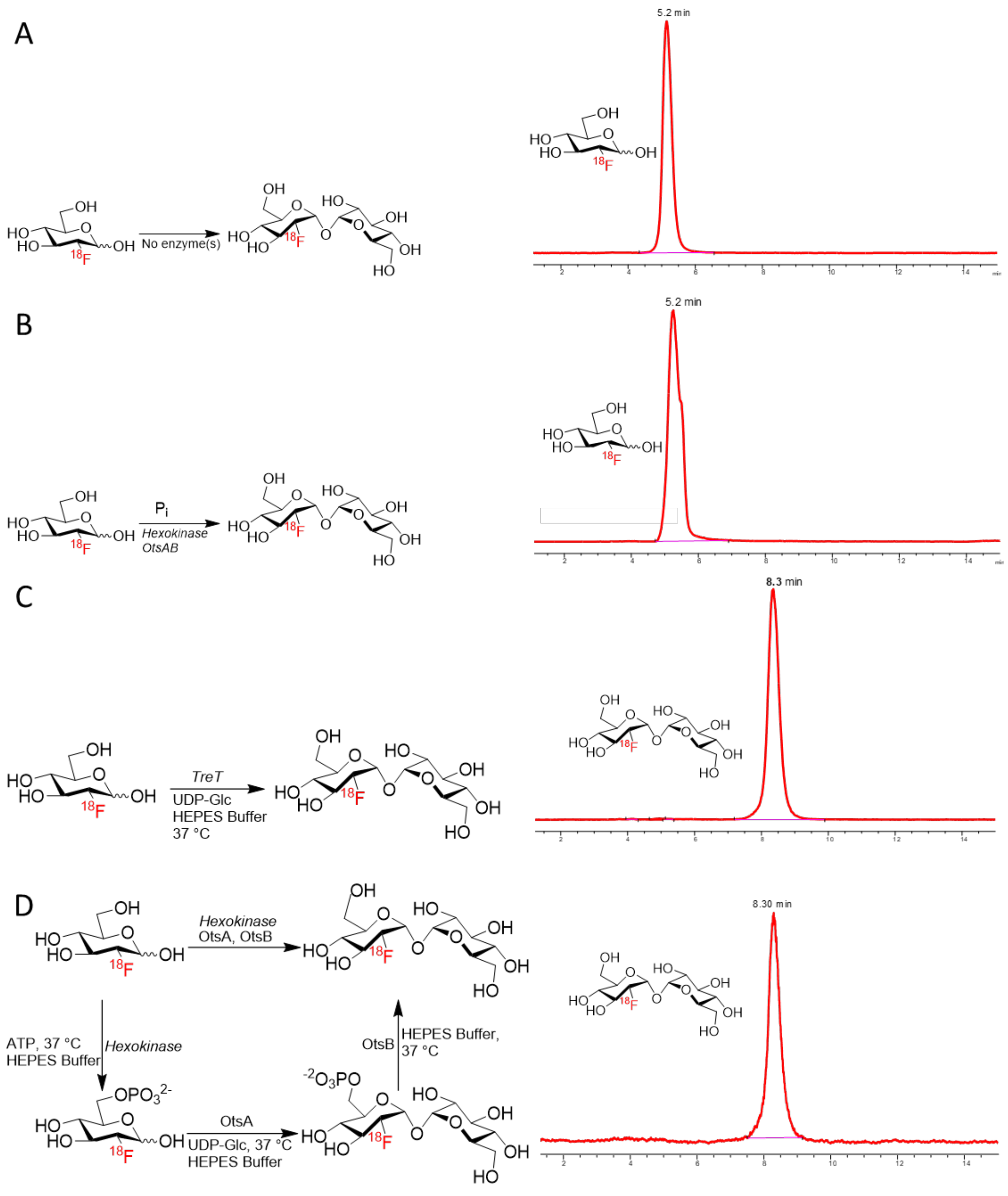

**Supplementary Figure S30. Enzymatic radiosynthesis of [$^{18}$F]FDT.**

**A**) Schematic overview and radio-HPLC chromatogram of control reaction, no enzyme was added in the reaction mixture. **B**) Schematic overview and chromatogram of attempted radiosynthesis of **[$^{18}$F]FDT** using 2-enzyme-2-step one-pot synthesis method which failed to yield any product and only [$^{18}$F]FDG observed after 2 h reaction. **C**) Schematic and radio-

HPLC chromatogram of **[$^{18}$F]FDT** synthesized from TreT enzyme; full conversion to the product observed after 60 min of reaction. **D**). Schematic overview of **[$^{18}$F]FDT** synthesised from hexokinase, OtsA and OtsB enzyme and radio-HPLC chromatogram; full conversion to the product observed after 60 min of reaction.

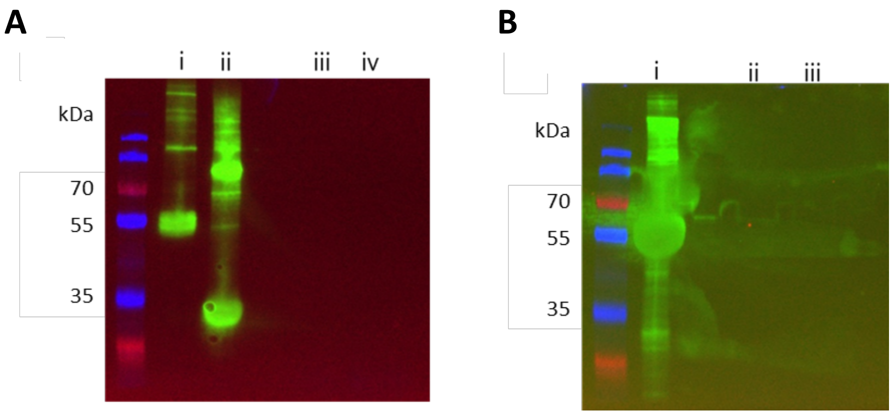

**Supplementary Figure S31. Western Blot Analyses of [$^{18}$F]FDT Samples.**

(**A**) Detection of OtsA and OtsB, using anti-His antibody. Lanes: (**i**) OtsA enzyme; (**ii**) OtsB enzyme; (**iii**) dilute **[$^{18}$F]FDT** sample; (**iv**) concentrated **[$^{18}$F]FDT** sample (**B**). Detection of hexokinase in **[$^{18}$F]FDT** sample using anti-hexokinase antibody. Lanes: (**i**) hexokinase enzyme only; (**ii**) dilute **[$^{18}$F]FDT** sample; (**iii**) concentrated **[$^{18}$F]FDT** sample. No enzymes were detected in the **[$^{18}$F]FDT** samples.

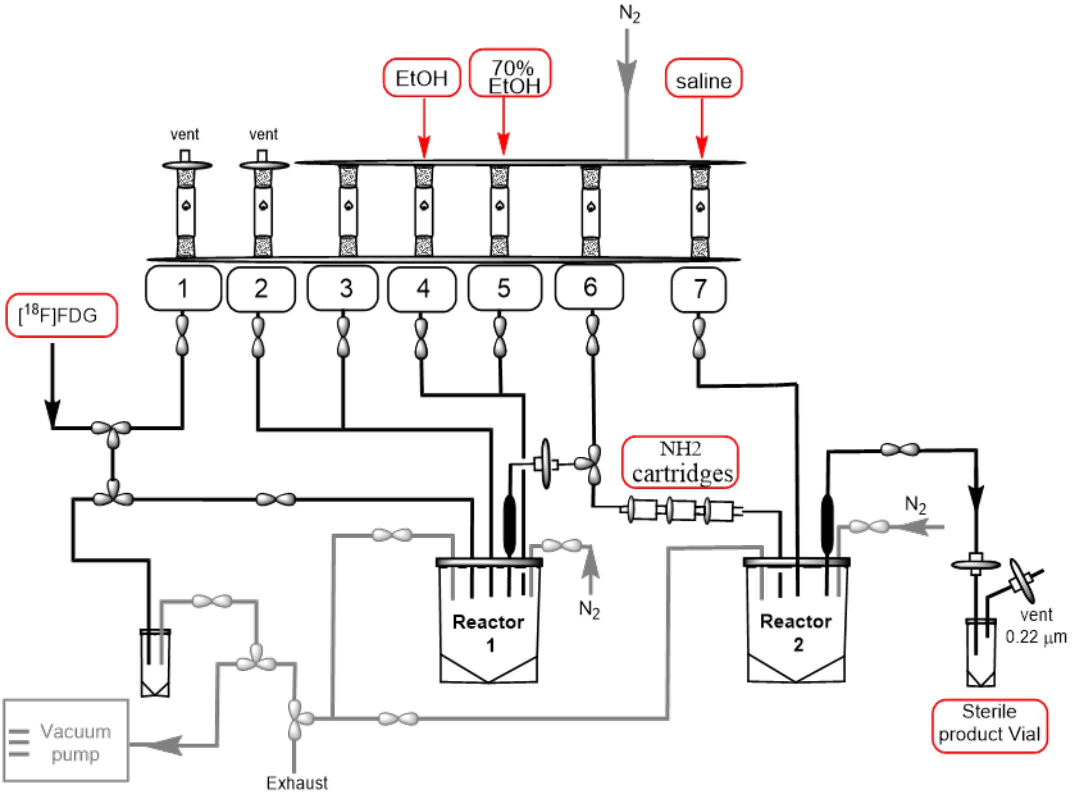

**Supplementary Figure S32. Schematic Diagram of Automated [$^{18}$F]FDT Synthesis on GE-FXN pro module**

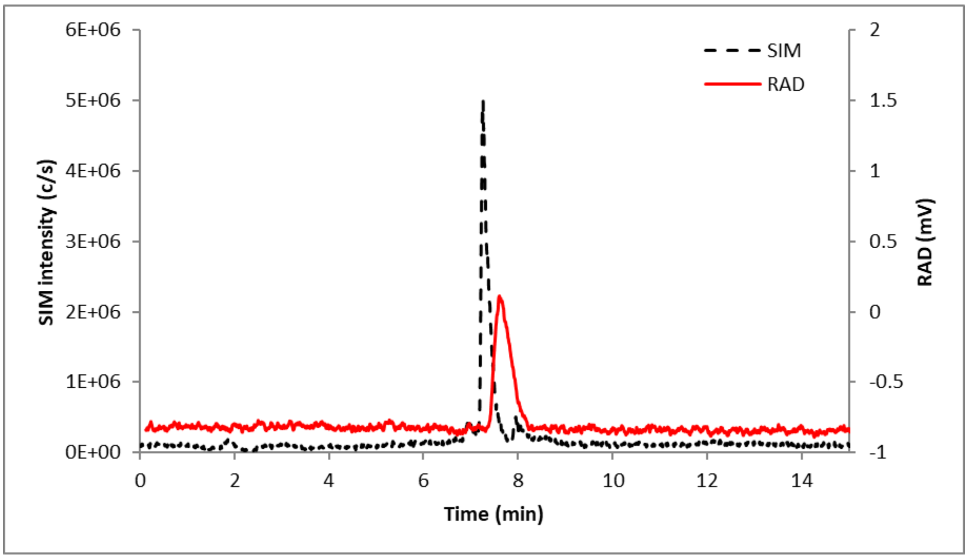

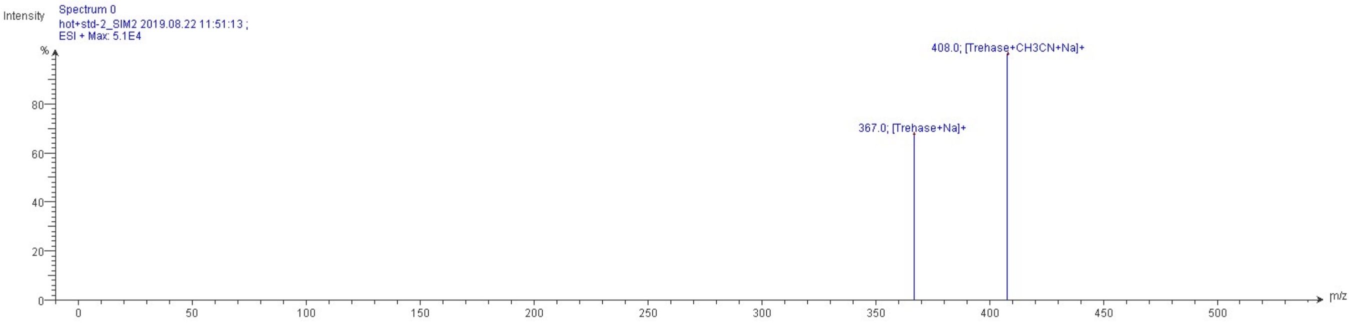

**Supplementary Figure S33. LC-MS analysis of [$^{18}$F]FDT coinjected with nonradioactive standard FDT.**

Solid line, in-line radiodetector; dotted line, UV detector at 254 nm (for **a**–c); for d: dotted line, SIM intensity of *m*/*z* 367. Chromatographic Conditions: Method B: Chromatographic separations were carried out with gradient elution on an Waters XBridge Amide column, 130Å, 3.5 µm, 4.6 mm X 150 mm. The mobile phase was composed of water (A) and acetonitrile (B) each containing 0.1 % ammonium hydroxide. The analytes were eluted from the column by a linear gradient which started at 90% B, then decreased to 50% B within 8 min. The total run time was 10 min. The flow rate was set at 1.0 mL/min. The column oven was kept at 40 °C throughout the analysis. The injection volume was 20 µL.

Mass Spectrometric conditions: Mass spectrometric data were acquired in positive ion mode with the following ESI-MS parameters: Capillary Temperature 250 ˚C, Capillary Voltage 120 V, Source Voltage Offset 25V, Source Voltage Span 50V, Source Gas Temperature 200 ˚C, ESI Voltage 3,500V. Nitrogen was used as a desolvation gas. The analysis was done using scan/single ion monitoring (SIM) switching mode. The parameters for the scan mode was set as follows: scan range 101500 m/z, scan time 376 ms, scan speed 3963 m/z/sec. In SIM mode dwell time was set at 50 ms for all analytes. Non radioactive standard trehalose, FDT (m/z calculated mass for $C_{12}H_{21}FO_{10}Na$ 367.10, found 408 [M+Na+ACN], 367 [M+Na]) at retention time 7.25 min was co-injected with radiolabeled product was used to confirm the identity of the analyte in the sample.

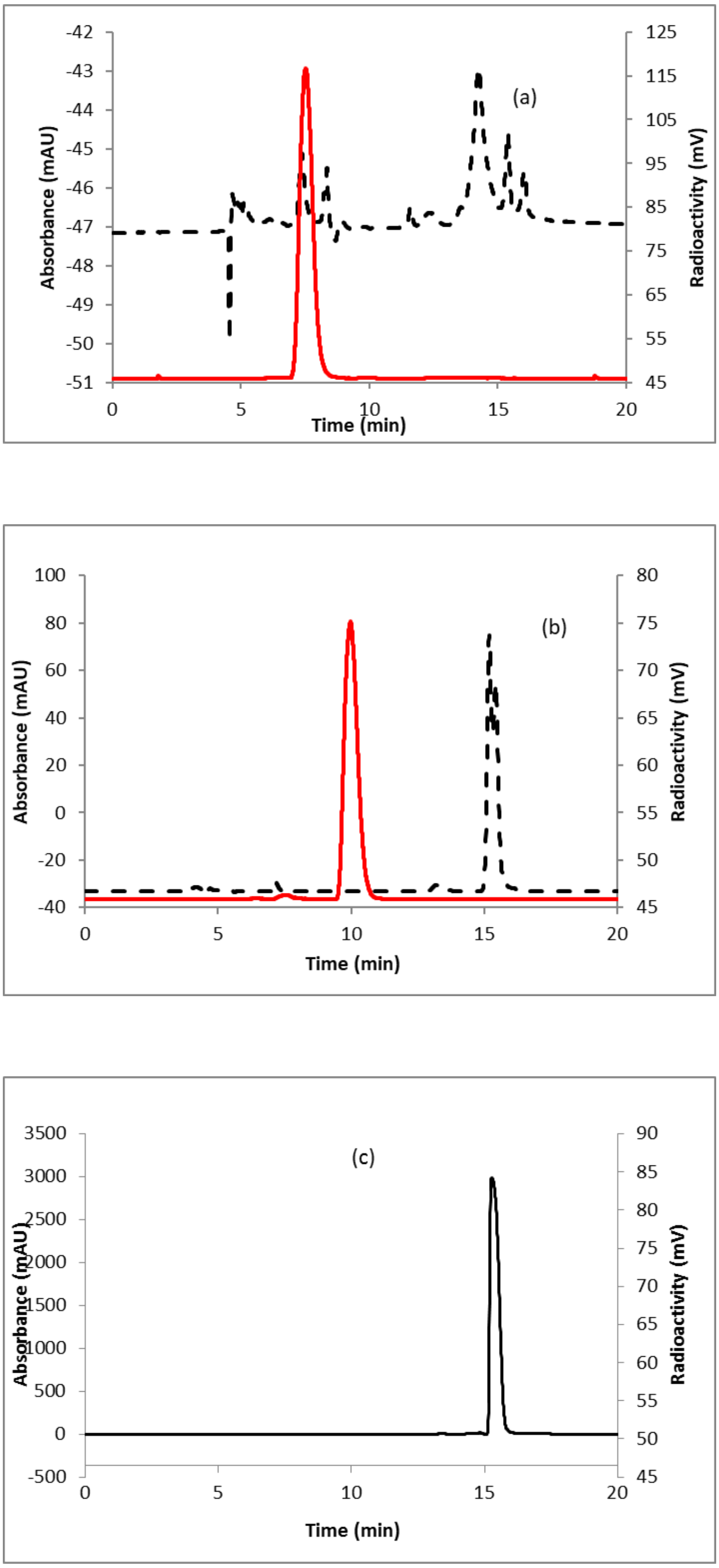
(a)
Absorbance (mAU)
Radioactivity (mV)
Time (min)
-42
-43
-44
-45
-46
-47
-48
-49
-50
-51
125
115
105
95
85
75
65
55
45
0
5
10
15
20
(b)
Absorbance (mAU)
Radioactivity (mV)
Time (min)
100
80
60
40
20
0
-20
-40
80
75
70
65
60
55
50
45
0
5
10
15
20
(c)
Absorbance (mAU)
Radioactivity (mV)
Time (min)
3500
3000
2500
2000
1500
1000
500
0
-500
90
85
80
75
70
65
60
55
50
45
0
5
10
15
20

**Supplementary Figure S34. HPLC Analyses of (A) [$^{18}$F]FDG; (B) [$^{18}$F]FDT; (C) UDP-glucose.**

4.6 x 250 mm, 2.7 µm AdvanceBio Glycan Mapping Column; solvent A = 50 mM ammonium formate, pH 4.5, solvent B = acetonitrile; Flow rate = 0.5 mL/min;gradient 0-15 min 68-62% B; 15-20 min 62-68% B. Retention. Red line, in-line radiodetection; black line, UV detection at 254 nm. The UV peak at ~15 min is for the UDP-glucose, confirmed by comparing the HPLC retention time of separately injected pure UDP glucose (**A**,**C**) and mass spec analysis of the decayed solution of [$^{18}$F]FDT (**B**).

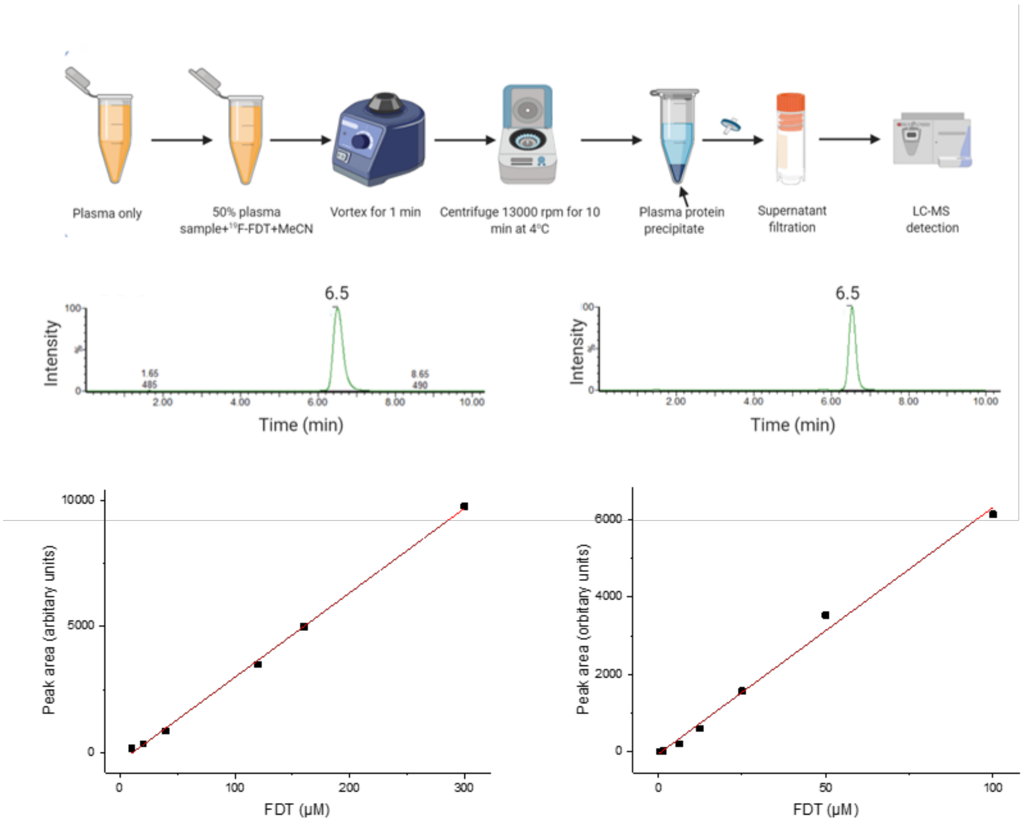

**Supplementary Figure S35. Assessment of [$^{19}$F]FDT Stability in Plasma**

**A**) Experimental set-up of the sample preparation. Commercially available plasma samples were spiked with known concentrations of [$^{19}$F] FDT, plasma proteins were then precipitated by adding up to 50% of MeCN solution and after centrifugation the samples were analysed by LC-MS. **B**) [$^{19}$F]FDT standard sample analysed by LC-MS method 2 with the retention time of 6.5 min. **C**) LC-MS analysis of plasma sample spiked with known amount of FDT showing the similar retention time to that of the standard sample. **D**) Calibration curve of plasma samples when spiked with FDT concentrations from 10 µM – 300 µM with $r^2$ = 0.998 and **E**) Further calibration curve showing the instrument sensitivity of up to 390 nM to accurately detect FDT with $r^2$ = 0.998.

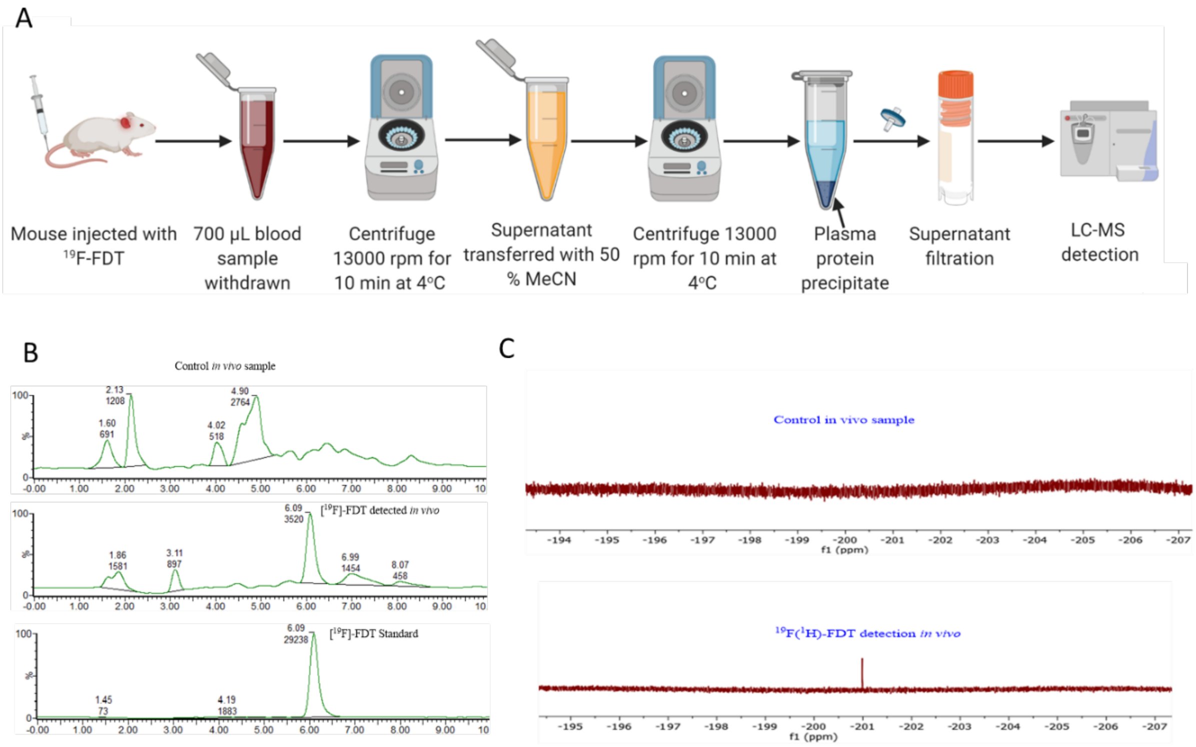

**Supplementary Figure S36. Assessment of [$^{19}$F]FDT Stability *In vivo*.**

**A**). Experimental set-up for the *in vivo* detection; **B**). From top to bottom; control *in vivo* plasma sample; middle [$^{19}$F] FDT detection in plasma *sample in vivo* when 500 µM of the [$^{19}$F] FDT solution injected in mouse and sample taken 5 min post injection; [$^{19}$F] FDT standard sample; **C**). From top to bottom, control *in vivo* sample and bottom [$^{19}$F] FDT detection *in vivo* by $^{19}$F-NMR spectroscopy.

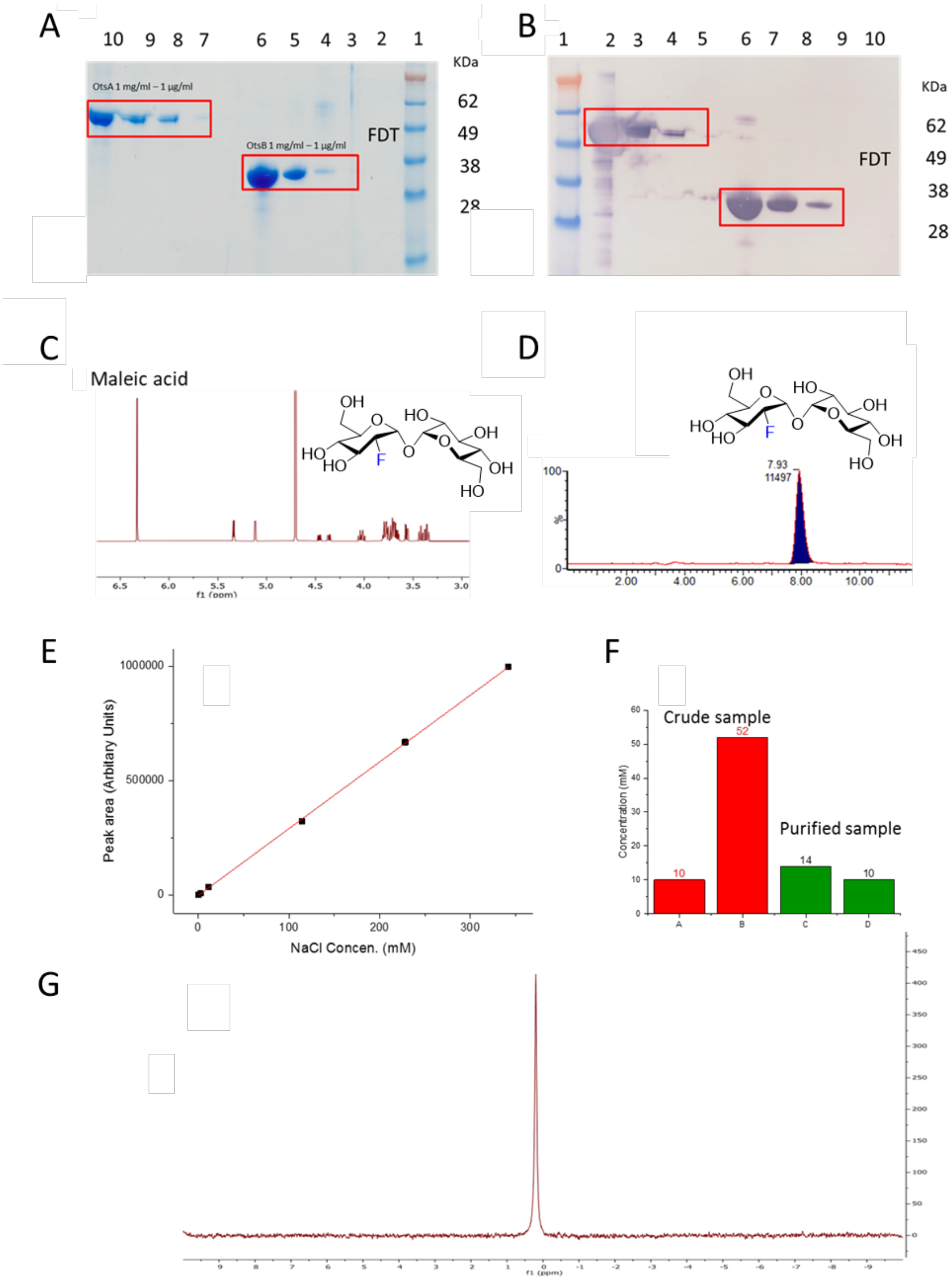

**Supplementary Figure S37. FDT Batch Quantification and Purity Testing.**

(**A**) SDS-PAGE analysis for detection of and enzyme leaching in the final FDT batch. Lanes: OtsA standards (lane 7-10) and OtsB standards (lane 3-6) with 1/10 serial dilutions loaded onto the SDS-PAGE. 25 mg/mL FDT from purified batch also loaded on the gel (lane 2). No enzyme was detected in the FDT sample. (**B**) Western blotting visualisation for the detection of enzyme leaching in the final FDT batch. Lanes: OtsA standard (lane 2-5) and OtsB standards (lane 6-9) enzymes with 1/10 serial dilutions from 1mg/mL to 1 µg/mL loaded onto the SDS-PAGE. 25 mg/mL FDT (lane 10) from purified batch also loaded. Visualisation was done with an anti-His monoclonal antibody. No protein was detected in the FDT sample even at this high concentration. (**C**). Processed quantitative $^{1}$H-NMR spectrum of analyte FDT. Maleic acid as an internal standard. (**D**) LC-MS Chromatogram of FDT. Retention times above; integral below for each noted peak. Peak at 7.93 min is FDT with major ESI$^{-}$ m/z of 343. (**E**) Standard calibration curve for NaCl using $^{23}$Na-NMR; **F**) Concentrations in FDT samples (determined by combined $^{19}$F-NMR and $^{23}$Na-NMR) in the crude reaction mixture (red) and purified (green). Column A is the concentration of FDT in the crude reaction mixture determined by $^{19}$F-NMR; column B shows the quantified value of NaCl as determined by $^{23}$Na-NMR. Columns C & D are the corresponding measurements from the purified sample of FDT**.** All are calculated from samples dissolved in 0.5 mL $D_2O$. Total integrals from the peak area were used for calculation using intercept and slope from the calibration curve. (**G**) Example $^{23}$Na-NMR spectrum for quantitative yield calculation of NaCl in samples.

### Supplementary Tables

**Table S1. Endotoxin test for the one-pot reaction mixture containing OtsA and OtsB from *E. coli* at each purification step.**

Endotoxin assay was carried out using EndosafeTM-PTS detection instrument (Charles River). Detection cartridges were 0.01 – 1.0 endotoxin unit (EU)/mL and 0.05 – 5.0 EU/mL. Protein concentration at each purification step was also carried out at the same time using BCA method (Thermo Scientific). The measured values for endotoxins were not averaged but presented as the range of measured concentrations while protein concentrations were averaged

| Samples | Protein (mg/mL) | Endotoxins (EU/mL) |
|---|---|---|
| Reaction mixture | > 3 | > 5,000 |
| After 1$^{st}$ SAX SPE | 0.43 | 200 – 392 |
| After 2$^{nd}$ SAX SPE | Background level | 0.05 – 5 |
| After 3$^{rd}$ SAX SPE | Background level | 0.01 – 0.051 |
| After HPLC | Background level | < 0.01 |

**Table S2. Effect of amount of [$^{18}$F]FDG on radiochemical yield**

| Type of Synthesis | Entry | [$^{18}$F]FDG mCi | [$^{18}$F]FDG Volume mL | Total Volume mL | % Yield, non-decay corrected |
|---|---|---|---|---|---|
| Automated | 1 | 40 | 0.8 | 1.1 | 26.3 ± 5.8 (n = 2) |
| | 2 | 60 | 1.0 | 1.3 | 26.3 ± 5.8 (n = 2) |
| | 3 | 100 | 1.8 | 2.1 | 41.0 ± 4.2 (n=3) |

**Table S3. LC-MS Analysis Gradient for LCT Classic MS**

| Time (min) | % Solvent A | % Solvent B | Flow rate (mL/min) |
|---|---|---|---|
| 0.0 | 95 | 5 | 0.4 |
| 1.0 | 95 | 5 | 0.4 |
| 7.0 | 5 | 95 | 0.4 |
| 11.0 | 5 | 95 | 0.4 |
| 12.0 | 95 | 5 | 0.4 |
| 17.0 | 95 | 5 | 0.4 |

**Table S4. LC-MS Analysis Gradient for Waters Xevo G2-QS QToF MS**

| Time (min) | % Solvent A | % Solvent B | Flow rate (mL/min) |
|---|---|---|---|
| 0.0 | 95 | 5 | 0.4 |
| 1.0 | 95 | 5 | 0.4 |
| 7.0 | 5 | 95 | 0.4 |
| 8.0 | 5 | 95 | 0.4 |
| 8.1 | 95 | 5 | 0.4 |
| 10.0 | 95 | 5 | 0.4 |

**Table S5: Summary of different HPLC conditions for enzymatic reaction analysis**

| Method | Mobile Phase | Stationary | Gradient | Flow rate | Run | UV |
|---|---|---|---|---|---|---|

| | | Phase | | (mL/min) | time (min) | detection (λ) |
|---|---|---|---|---|---|---|
| 1 | A. $H_2O$ B.Ammonium formate 300 mM | waters spherisorb 5 µm SAX 4.6 x 250 nm | 5-100% B linear gradient | 1 | 30 | 262 |
| 2 | A.$H_2O$ B.$H_2KPO_4$ 100 mM | waters spherisorb 5 µm SAX 4.6 x 250 nm | 5-100% B linear gradient | 1 | 30 | 262 |
| 3 | A. $H_2O$ B. $H_2KPO_4$ 1M | Phenomenex Luna $NH_2$ 4.6 x 250 nm | 0-100% linear gradient over 60 min, 61-65 min, 100% A | 0.6 | 65 | 262 |

**Table S6: Reagents Details for Enzyme Comparison Reaction**

| Enzyme | FDG (mM) | UDP-Glc (mM) | $MgCl_2$ (mM) | Enzyme (μM) | HEPES Buffer (mM) |
|---|---|---|---|---|---|
| TreT *P. horikoshii* | 10 | 20 | 10 | 1.2 | 50 mM, pH 7.5 |
| TreT *T. tenax* | 10 | 20 | 10 | 1.2 | 50 mM, pH 7.5 |

**Table S7. Testing ATP concentrations for the conversion of FDG to FDG6P**

| Entry | FDG (mM) | ATP (mM) | $MgCl_2$ (mM) | Enzyme (mg) | HEPES Buffer (mM) |
|---|---|---|---|---|---|
| 1 | 5.55 | 2.75 | 10 | 5 | 50 mM, pH 7.5 |
| 2 | 5.55 | 5.50 | 10 | 5 | 50 mM, pH 7.5 |
| 3 | 5.55 | 8.25 | 10 | 5 | 50 mM, pH 7.5 |
| 4 | 5.55 | 16.5 | 10 | 5 | 50 mM, pH 7.5 |

**Table S8: Reagent Amounts Used in the Small scale Synthesis of FDT Using a 3-enzyme-one-pot System**

| Entry | FDG (mmol) | ATP (mmol) | $MgCl_2$ (mmol) | KCl (mmol) | UDP-Glc (mmol) | hexokinase (mg) | OtsA (mg) | OtsB (mg) | HEPES (50 mM) | Reaction volume (mL) |
|---|---|---|---|---|---|---|---|---|---|---|
| 1 | 0.1380 | 0.7259 | 5 | 100 | 20 | 1 | 1 | 0.1 | Diluent | 1 |
| 2 | 0.2760 | 0.7259 | 5 | 100 | 20 | 1 | 1 | 0.1 | Diluent | 1 |
| 3 | 0.4140 | 0.7259 | 5 | 100 | 20 | 1 | 1 | 0.1 | Diluent | 1 |
| 4 | 0.5520 | 0.7259 | 5 | 100 | 20 | 1 | 1 | 0.1 | Diluent | 1 |
| 5 | 1.1040 | 7.0780 | 5 | 100 | 20 | 1 | 1 | 0.1 | Diluent | 1 |
| 6 | 2.2082 | 7.0780 | 5 | 100 | 20 | 1 | 1 | 0.1 | Diluent | 1 |
| 7 | 4.4164 | 7.0780 | 5 | 100 | 20 | 1 | 1 | 0.1 | Diluent | 1 |
| 8 | 5.5205 | 7.0780 | 5 | 100 | 20 | 1 | 1 | 0.1 | Diluent | 1 |

**Table S9. LC-MS methods employed in the detection of FDT**

| **Method no.** | **Mobile phase** | **Flow rate (mL/min)** | **Run time (min)** | **Temperature (°C)** | **FDT rt (min)** |
|---|---|---|---|---|---|
| 1 | 30/70 $H_2O$/MeCN | 0.8 | 20 | 25 | 10.5 |
| 2 | 35/65 $H_2O$/MeCN | 0.3 | 12 | 25 | 8.7 |
| 3 | 35/65 $H_2O$/MeCN | 0.4 | 12 | 25 | 6.5 |
| 4 | 35/65 $H_2O$/MeCN | 0.4 | 12 | 60 | 6.1 |
